# Triple tandem trimer immunogens for HIV-1 and influenza nucleic acid-based vaccines

**DOI:** 10.1101/2023.08.27.554987

**Authors:** Iván del Moral-Sánchez, Edmund G. Wee, Yuejiao Xian, Wen-Hsin Lee, Joel D. Allen, Alba Torrents de la Peña, Rebeca Fróes Rocha, James Ferguson, André N. León, Sylvie Koekkoek, Edith E. Schermer, Judith A. Burger, Sanjeev Kumar, Robby Zwolsman, Mitch Brinkkemper, Aafke Aartse, Dirk Eggink, Julianna Han, Meng Yuan, Max Crispin, Gabriel Ozorowski, Andrew B. Ward, Ian A. Wilson, Tomáš Hanke, Kwinten Sliepen, Rogier W. Sanders

## Abstract

Recombinant native-like HIV-1 envelope glycoprotein (Env) trimers are used in candidate vaccines aimed at inducing broadly neutralizing antibodies. While state-of-the-art SOSIP or single-chain Env designs can be expressed as native-like trimers, undesired monomers, dimers and malformed trimers that elicit non-neutralizing antibodies are also formed, implying that these designs could benefit from further modifications for gene-based vaccination approaches. Here, we describe the triple tandem trimer (TTT) design in which three Env protomers are genetically linked in a single open reading frame and express as native-like trimers. Viral vectored Env TTT induced similar neutralization titers but with a higher proportion of trimer-specific responses. The TTT design was also applied to generate influenza hemagglutinin (HA) trimers without the need for trimerization domains. Additionally, we used TTT to generate well-folded chimeric Env and HA trimers that harbor protomers from three different strains. In summary, the TTT design is a useful platform for the design of HIV-1 Env and influenza HA immunogens for a multitude of vaccination strategies.

## Introduction

One of the main goals of the HIV-1 vaccine field is the generation of recombinant envelope glycoprotein (Env) immunogens that can elicit protective broadly neutralizing antibody (bNAb) responses^1–6^. Such bNAbs develop in a subset of chronically infected patients and have been shown to be protective in titers achievable by vaccination^7–9^. Env is a class I fusion trimeric protein that engages CD4 and its CCR5 or CXCR4 coreceptor to mediate target cell fusion^10–14^. The *env* gene encodes a gp160 precursor polyprotein that is synthesized and initially trimerizes in the endoplasmic reticulum. N-linked glycosylation and furin cleavage of the gp160 protein into mature gp120 and gp41 subunits are essential for folding into its native prefusion conformation^15,16^. The Env complexes on the viral surface consist of three heterodimeric gp120-gp41 protomers. The membrane proximal external region (MPER) of gp41 is attached to the membrane by a coiled coil transmembrane domain that aids in trimerization of the complex. The gp120 subunits interact with gp41 by non-covalent interactions^17,18^, which can result in shedding of some of the gp120 and exposure of gp41 ‘decoys’ that participate in immune evasion^19,20^.

The first-generation recombinant HIV-1 Env immunogens consisted of unstable complexes that expose non-neutralizing antibody (non-NAb) epitopes, which are normally not exposed on infectious viral Env^4,21,22^. These non-NAb epitopes are considered undesirable because they attract immunodominant responses and might distract the immune system from targeting the desired neutralizing antibody (NAb) epitopes. Furthermore, these non-native Env immunogens do not properly display several quaternary-dependent antibody epitopes^23–26^.

It took several years of iterative design to generate soluble native-like Env immunogens, including SOSIP trimers^27–30^. SOSIP modifications include the truncation of gp41 at position 664, a disulfide bond (501C-605C) to covalently link the gp120 and gp41 subunits^27^, an Ile-to-Pro mutation (I559P) to prevent conformational transitions to the post-fusion state^29^ (a strategy that was also applied for many COVID-19 vaccines^31^), and an RRRRRR (R6) multibasic motif to enhance furin cleavage^28,32^. The determination of Env SOSIP trimer structures^33,34^ led to a plethora of further structure-based stabilizing mutations and novel Env trimer designs, such as single-chain (SC; also called native flexibly-linked, NFL), and uncleaved prefusion-optimized (UFO) Env trimers^35–37^. These SC and UFO trimers contain a flexible linker that connects the gp120 and gp41 subunits and allows a native-like conformation without the need of furin cleavage^37,38^. Several native-like Env trimers based on the SOSIP and SC designs are currently being tested in phase I clinical trials^31,39^.

Gene-based vaccines, including viral-vectored and nucleic acid immunogens (e.g. mRNA and DNA), have recently gained momentum because of their simplicity, reduced development and manufacturing costs, and advances in delivery methods^40–43^. For instance, viral vectors and mRNA-containing lipid nanoparticles (mRNA-LNPs)^43^ have been shown to efficiently elicit both NAb and cellular immune responses^44,45^. Furthermore, platforms such as self-amplifying RNAs^46,47^ and integrase-deficient lentivirus vectors (IDLVs)^48,49^, which allow for prolonged antigen exposure, may further enhance immune responses. Most recently, nucleic acid vaccines have gained particular importance in combatting the SARS-CoV-2 pandemic^43,50^, building an excellent safety record along the way.

During the production of Env proteins for vaccination, cells transfected with state-of-the-art HIV-1 Env designs, such as SOSIP and SC, produce the desired native-like trimers, but also monomeric, dimeric and malformed trimeric species that expose undesired epitopes **(Fig. S1).** Therefore, size exclusion and/or affinity chromatography purification methods are required to obtain homogeneous protein preparations containing only native-like trimeric species^30,35,36^. As selective purification is impractical during gene-based vaccination, these Env designs may not be optimal for nucleic acid vaccination approaches. One of the strategies to alleviate this problem is the use of heterologous trimerization-inducing domains, like the GCN4 leucine zipper 49^51–53^ or the bacteriophage T4 fibritin foldon^54,55^ motifs, which enhance trimerization of several class I fusion proteins, including HIV-1 Env^23,56–59^ and influenza hemagglutinin (HA)^60–62^. However, these heterologous domains can induce aberrant Env conformations^63^ and also immunodominant antibody responses that might hinder Env- or HA-targeting NAb responses^59^.

A recombinant Env design that expresses mostly as native-like trimers would therefore be a great advance for unlocking the potential of genetic vaccination for induction of bNAbs against HIV-1. Here, we describe a novel Env design, named Triple Tandem Trimer (TTT), in which we genetically fused three Env protomers by flexible linkers. We generated Env TTT constructs that expressed only as trimers, the majority of which were native-like. Using viral vector-based vaccination, we found that TTT constructs induce less non-neutralizing responses than other recombinant Env trimer designs. We also applied the TTT design for generation of influenza hemagglutinin (HA) and chimeric Env and HA constructs that express only as trimers and present native-like features.

## Results

### Design and characterization of a native-like Env triple tandem trimer (TTT)

Ideally, an HIV-1 Env construct intended for nucleic acid-based vaccination should only express trimers and should not require furin cleavage for native-like folding **(Fig. S1).** Therefore, we conceived a triple tandem trimer (TTT) design, encompassing three successive gp140 protomers genetically linked in a single open reading frame **(Fig. 1a,b)**. As proof of concept, we generated a BG505 SOSIP.v8.4 TTT construct. Each BG505 SOSIP.v8.4 (referred to as SOSIP.v8 herein) protomer includes all of the SOSIP.v4.1 mutations^64^, as well as additional modifications (MD39) that improve thermostability and trimerization^65^. The individual mutations are detailed in the *Methods* section and **Table S1.** Moreover, the R6 furin cleavage site^28^ was replaced by a 15-residue flexible linker to assure furin independence **(Fig. 1a,b)**, akin to previous uncleaved single-chain (SC) Env constructs^35,36^. Next, we linked the gp41 C-terminus of each protomer to the gp120 N-terminus of the neighboring protomer **(Fig. 1a,b).** The termini are separated by ∼11 Å **(Fig. 1b)**; therefore, we introduced a relatively long 11-residue flexible linker between the protomers to allow enough rotational freedom for trimer assembly. Additionally, this BG505 TTT construct contained a C-terminal StrepII-Tag for purification and immobilization purposes. The AlphaFold2 software predicted that such a construct would fold with a structure similar to the experimentally determined structure of a native-like BG505 Env trimer **(Fig. S2).**

**Fig. 1.**
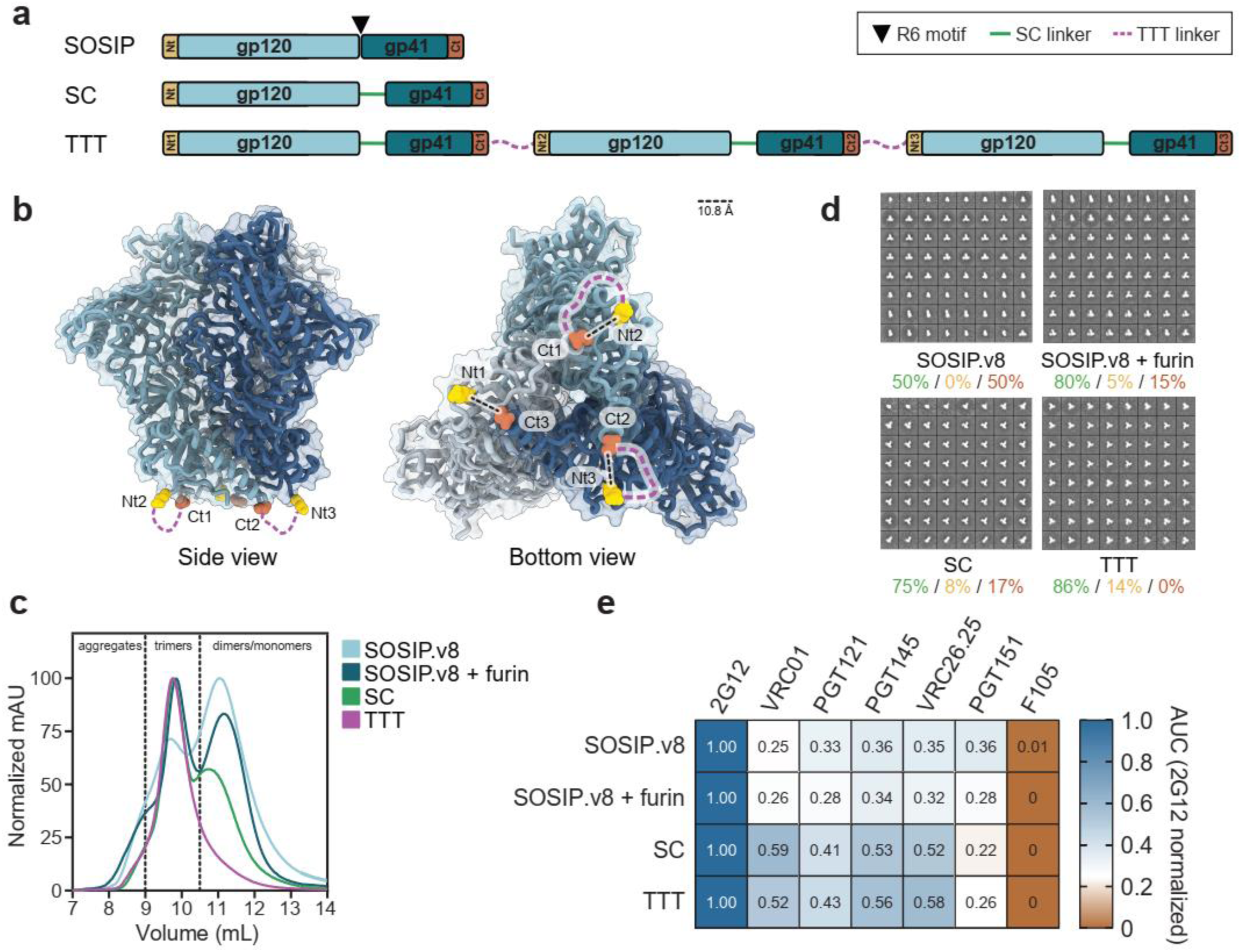
Design and biophysical characterization of a triple tandem trimer Env construct. **a** Linear representation of SOSIP, single-chain (SC) and triple tandem trimer (TTT) Env constructs. The SOSIP construct presents a R6 furin cleavage motif (black triangle) between gp120 and gp140 subunits, which is replaced by SC linkers (green lines) in SC and TTT constructs. The TTT construct is composed of three SC protomers fused by TTT linkers (purple discontinuous lines). N- and C-termini of each protomer are colored in yellow and red, respectively. **b** Expected position of TTT linkers on the structure of a BG505.SOSIP trimer (PDB 5CEZ). The distance, in angstrom (Å), between the C_ɑ_ atoms of N-(Glu32, HxB2 numbering) and C-termini (Asp664) of neighbor protomers is indicated. **c** SEC profiles of StrepTactinXT-purified BG505 SOSIP.v8, SOSIP.v8 + furin, SC and TTT protein preparations on a Superdex 200 Increase 10/300 GL column. **d** 2D class averages generated by nsEM analysis of the StrepTactinXT-purified SOSIP.v8, SC and TTT protein preparations. Percentages of native-like trimers (green), malformed trimers (yellow) and monomers/dimers (red) are indicated. **e** StrepTactinXT ELISA assay with StrepTactinXT-purified SOSIP.v8, SC and TTT protein preparations against a panel of bNAbs (2G12, VRC01, PGT121, PGT145, VRC26.25, PGT151) and non-NAbs (F105). Heatmap values correspond to the ratio between the areas under the curve (AUC) of each Ab and the one of 2G12, calculated using the binding curves in **Fig. S4c.**

To initially assess the expression and antigenicity of BG505 TTT, we performed a StrepTactinXT ELISA assay with unpurified supernatants from HEK293T cells transfected with the BG505 TTT construct, as well as BG505 SOSIP.664, SOSIP.v8 and single-chain SOSIP.v8 (herein termed SC) control constructs **(Fig. S3).** The TTT supernatant interacted well with all tested bNAbs, including those targeting quaternary epitopes (PGT145, VRC26.25 and PGT151^66–69^), while showing weak binding to non-NAb F105.

Next, we purified BG505 TTT from transiently transfected HEK293F suspension cells using StrepTactinXT affinity chromatography, an unbiased purification method that should capture all Env, not specifically native-like trimers. The TTT protein yields (∼2.1 mg/L) were similar to those obtained from cells transfected with SC or SOSIP.v8, without or with furin co-expression (∼2.8 mg/L, ∼2.7 mg/L and ∼3.4 mg/L, respectively) **(Table 1).** However, size-exclusion chromatography (SEC) and PAGE analysis showed that the BG505 TTT preparation consisted of covalently linked trimers exclusively, while SOSIP.v8 and SC controls also contained substantial amounts of dimers and monomers **(Fig. 1c, Fig. S4a).** Negative-stain electron microscopy (nsEM) revealed that >85% of StrepTactinXT-purified TTT trimers displayed a native-like morphology, while StrepTactinXT-purified SC and SOSIP.v8 preparations contained substantial amounts of dimers and monomers **(Fig. 1d**, **Table 1).** We obtained high protein yields (∼16 mg/L) and similar percentages of native-like trimers when we purified BG505 TTT using *Galanthus nivalis* lectin (GNL) chromatography **(Table 1, Fig. S5).**

**Table 1.** Production yields and biophysical characterization of the different Env and HA proteins.

| Protein | Yield<br>(mg/L) (≈) | Native-like<br>(%) (>) | T<sub>m</sub><br>(°C) |
| --- | --- | --- | --- |
| SOSIP.v8<sup>a</sup> | 2.7 | 50 | ND |
| SOSIP.v8 + Furin<sup>a,b</sup> | 3.4<sup>a</sup>/10.8<sup>b</sup> | 80<sup>a</sup> | ND |
| SC<sup>a,b</sup> | 2.8<sup>a</sup>/15.1<sup>b</sup> | 75<sup>a</sup> | ND |
| TTT<sup>a,b</sup> | 2.1<sup>a</sup>/15.7<sup>b</sup> | 86<sup>a</sup>/82<sup>b</sup> | 76.9<sup>b</sup> |
| TTT.GM<sup>b</sup> | 6.9 | 82 | 73.9 |
| chEnv TTT<sup>c</sup> | 0.4 | 89 | 69.3 |
| HA<sup>a</sup> | 7.6 | ND | 49.0 |
| HA GCN4<sup>a</sup> | 6.5 | ND | 58.8 |
| HA TTT<sup>a</sup> | 3.0 | ND | 54.5 |
| H1 monomeric<sup>a</sup> | 1.1 | ND | 56.6 |
| H1 GCN4<sup>a</sup> | 4.9 | ND | 57.6 |
| H2 GCN4<sup>a</sup> | 0.9 | ND | ND |
| H5 GCN4<sup>a</sup> | 6.6 | ND | 55.9 |
| cH125 TTT<sup>a</sup> | 0.2 | ND | 55.9 |
| H1/1 monomeric<sup>a</sup> | 10.3 | ND | 56.7 |
| H1/1 GCN4<sup>a</sup> | 10.0 | ND | 57.2 |
| H2/1 GCN4<sup>a</sup> | 4.3 | ND | 51.3 |
| H5/1 GCN4<sup>a</sup> | 2.0 | ND | 61.0 |
| cH125/111 TTT<sup>a</sup> | 1.2 | ND | 58.7 |

<sup>a</sup> StreptactinXT-purified, <sup>b</sup> GNL-purified, <sup>c</sup> PGT145-purified

Furthermore, the BG505 TTT protein retained favorable antigenicity and engaged quaternary-dependent PGT145 and VRC26.25 more efficiently than SOSIP.v8 controls, while the linkers in BG505 TTT did not disrupt binding of the gp120-gp41 interface-targeting PGT151 bNAb **(Fig. 1e, Fig. S4c).** Nano Differential scanning fluorimetry (nanoDSF) showed that TTT displayed a midpoint melting temperature (T_m_) of 76.9 °C, slightly lower than one of the most stable BG505 SOSIP proteins described to date: BG505 SOSIP.v9.3 protein (80.7 °C)^70^ **(Table 1, Fig. S4b).** Glycosylation plays a major role in the folding, antigenicity and immunogenicity of HIV-1 Env^71^. To interrogate the glycan shield of BG505 TTT, we performed site-specific glycan analysis and compared the glycan abundance and species at each position to those from regular furin-cleaved BG505 SOSIP.v8. This analysis revealed that the glycan occupancy and processing of BG505 TTT was comparable with that of SOSIP trimers^72^, and that key glycan bNAb epitopes are conserved across both formats **(Fig. S4d, Table S2)**.

### BG505 TTT closely mimics the native-like Env conformation

To assess whether the TTT design allows native-like folding at the molecular level, we determined a crystal structure at 5.8 Å of 2G12-purified BG505 TTT in complex with PGT124 Fabs and 35O22 scFvs **(Fig. 2a, Table S3).** The structure of BG505 TTT is similar to that of other native-like BG505 Env trimers, as indicated by a C_ɑ_ root-mean-square deviation (RMSD) of 0.8 Å and similar distance maps **(Fig. 2c,d).** Although the relatively low resolution precluded the assignment of side chains and their interactions, PGT124 and 35O22 bound BG505 TTT with similar angles of approach to those observed with other Env trimers^73^. We did not observe density for the SC and TTT linkers, probably due to their high flexibility. However, the distance between the N- and C-termini of the BG505 TTT protomers (Glu32 to Leu663, ∼12 Å) **(Fig. 2b)** is comparable to their distance in BG505 SOSIP.664 (Glu32 to Asp664, ∼11 Å) **(Fig. 1b).** To evaluate the packing between BG505 TTT protomers, we measured the interprotomer distances between the C_ɑ_ atoms of all Env residues **(Fig. S6a),** including the ones selected by Stadtmueller et al. to compare trimers in various states of ‘openness’ (106, 173, 202, 306 and 542)^74^ **(Fig. 2e).** The interprotomer distances within BG505 TTT are comparable to the closed prefusion BG505 SOSIP trimers, but differ substantially from the distances measured in BG505 trimers in an open conformation **(Fig. 2e, Fig. S6a).** Furthermore, the BG505 TTT disulfide bond network is consistent with other Env trimers (**Fig. S6b)**. Overall, these observations confirm that the TTT design allows for proper folding and oligomerization of the individual protomers resulting in a conformation closely resembling those of other native-like Env trimers.

**Fig. 2.**
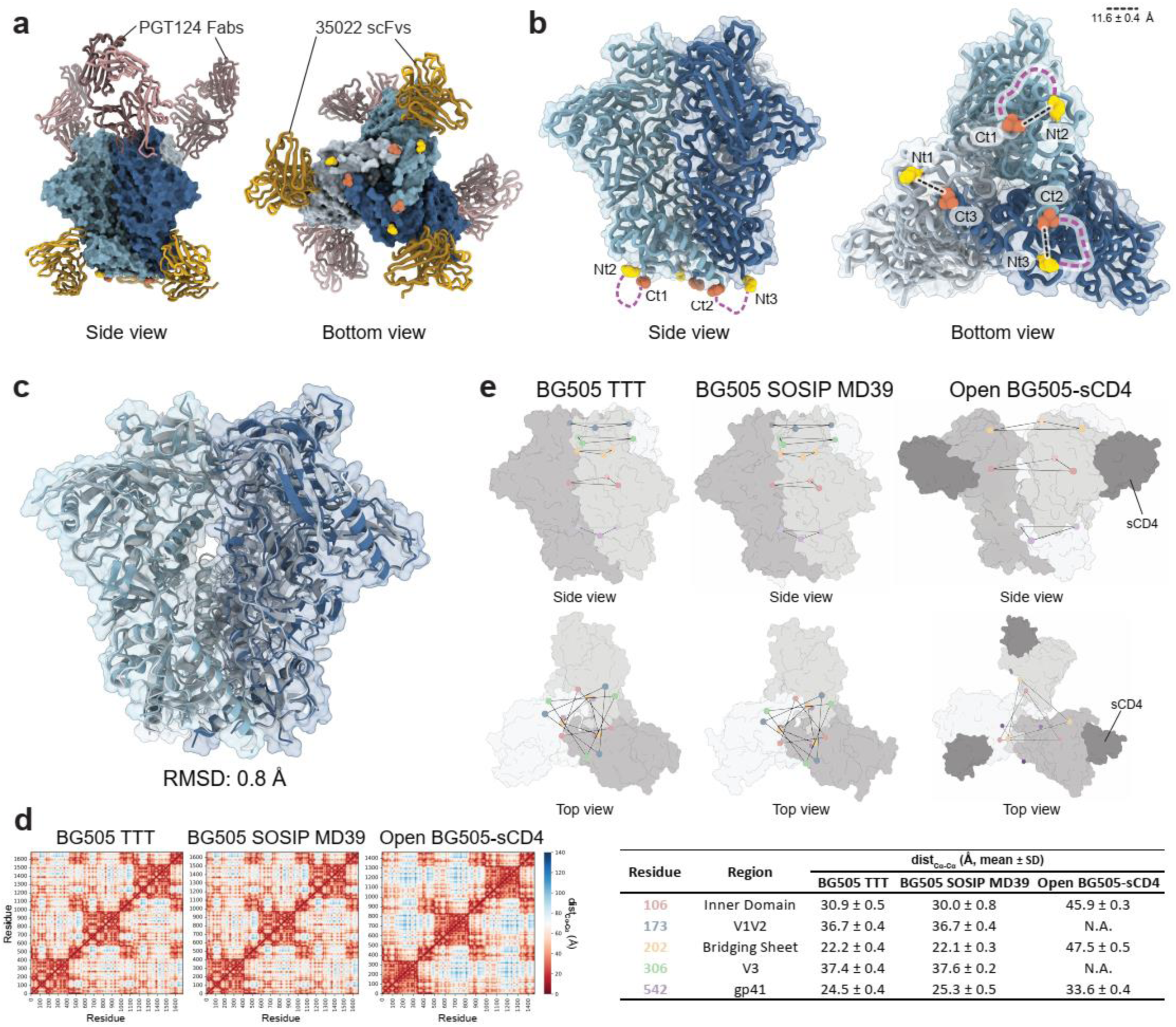
BG505 TTT closely mimics the native-like Env conformation. **a** Crystal structure of 2G12-purified BG505 TTT (blue) in complex with PGT124 Fabs (dark pink) and 35O22 scFvs (dark yellow) at 5.8 Å resolution. **b** Expected position of TTT linkers (purple dashed lines) on the structure BG505 TTT. The distance, in angstrom (Å), between the C_ɑ_ atoms of N-(Glu32, HxB2 numbering, yellow) and C-termini (Leu663, red) of neighbor protomers is indicated. **c** Alignment of BG505 TTT and BG505 SOSIP MD39 (PDB 7L8D) structures, with the corresponding C_ɑ_ root-mean-square deviation (RMSD) value. **d,e** Distance maps **(d)** and interprotomer distances between the C_ɑ_ atoms of residues (106, 173, 202, 306, 542) **(e)** of BG505 TTT, BG505 SOSIP MD39 and a BG505 Env trimer in a CD4-induced open conformation (PDB 5THR).

### BG505 TTT elicits qualitatively improved antibody responses in mice and rabbits

Simian (chimpanzee) adenovirus-derived ChAdOx1^75^ and modified vaccinia Ankara (MVA)^76^ are well-established vector systems for the delivery of immunogens^77–84^. ChAdOx1 was used by AstraZeneca to deliver one of the first effective SARS-CoV-2 vaccines^50^. MVA vaccines were used for smallpox eradication and are currently being used against the 2022 mpox outbreak^80^. Since BG505 TTT immunogens may be especially suitable for gene-based vaccination, including viral vector delivery, we generated ChAdOx1 and MVA vectors encoding either BG505 SOSIP.664 or TTT. Both ChAdOx1.TTT and MVA.TTT efficiently expressed as Env trimers in transduced HeLa cells **(Fig. S7).**

To evaluate the immunogenicity of vector-delivered BG505 TTT, we vaccinated four groups of mice and rabbits at weeks 0, 4 and 8 or weeks 0, 8, 24 and 40, respectively **(Fig. 3a,b).** For both models, two groups of animals were primed with ChAdOx1 (C)- and MVA (M)-delivered SOSIP.664 (group 1, SOSIP.664(CMP)) or TTT constructs (group 2, TTT(CMP)), and boosted with purified SOSIP.664 protein (P). In parallel, group 3 animals were immunized with three doses of purified SOSIP.664 protein (SOSIP.664(PPP)). For the mice, group 4 received a similar TTT(CMP) regimen as group 2. However, groups 3 and 4 mice were co-immunized, eight weeks apart (C8M), with ChAdOx1-(C1C62) and MVA-delivered (M3M4) HIVconsvX immunogens for inducing T cell responses against conserved regions of Gag and Pol^82,85^. Group 4 rabbits were co-immunized with viral-vectored TTT and SOSIP.664 protein at weeks 0 and 8, and were finally boosted with SOSIP.664 protein (TTT(CP.MP.P)). Sera were collected two weeks after each immunization to characterize the polyclonal antibody responses in mice **(Fig. 3c, Fig. S8a,d, Table S4)** and rabbits **(Fig. 3d, Fig. S9, Tables S7,S8)**, as well as the T cell responses in mice **(Fig. S8b,c, Tables S5,S6).**

**Fig. 3.**
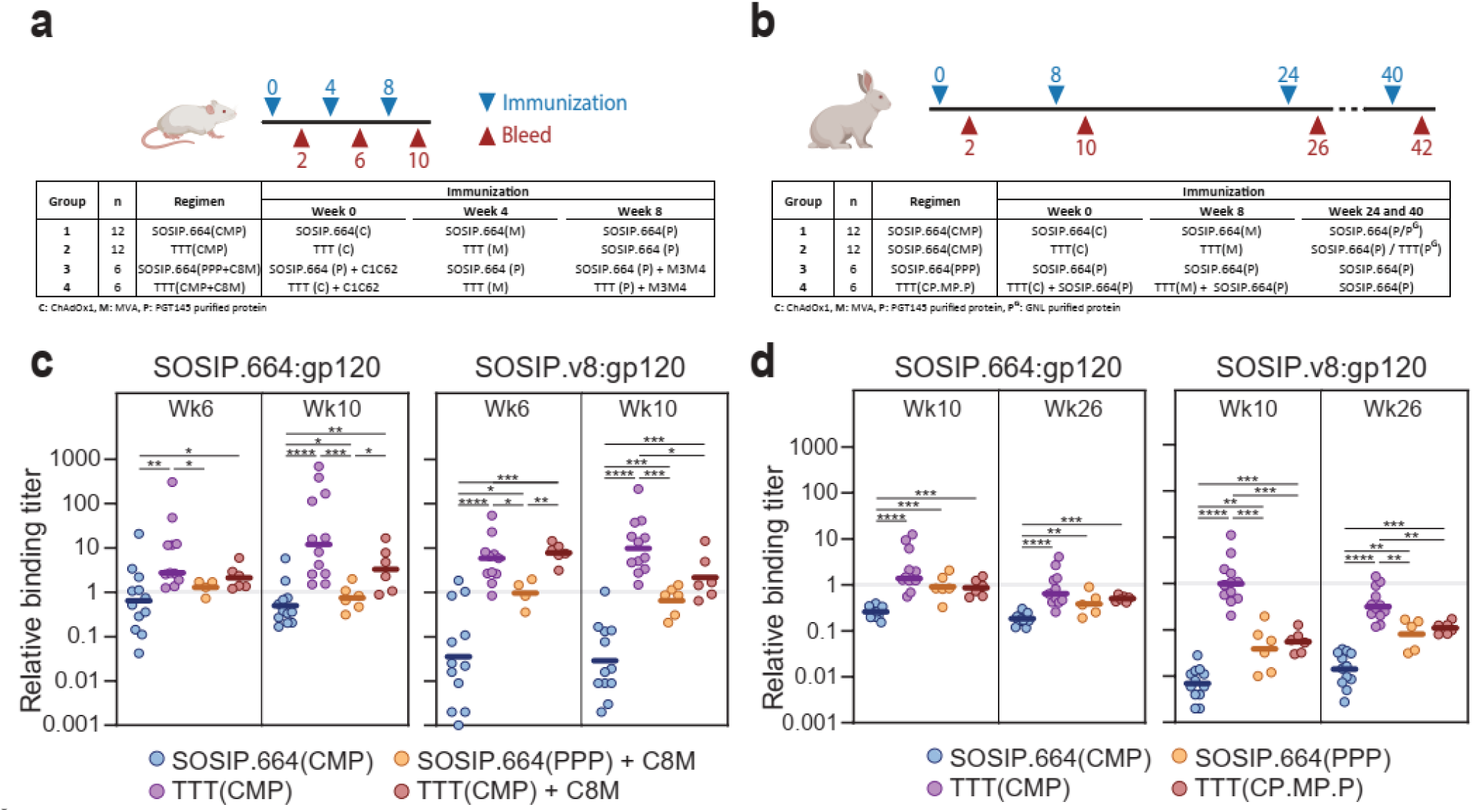
Immunogenicity of viral-vectored BG505 TTT. **a,b** Mice **(a)** and rabbits **(b)** immunization schedules. Four groups of BALB/c mice were immunized (blue arrows) at weeks 0, 4 and 8, and antibody responses were evaluated (red arrows) at weeks 2, 6 and 10. Four groups of New Zealand White rabbits were immunized at weeks 0, 8, 24 and 40, and antibody responses were evaluated at weeks 10, 26 and 42. The tables show the detailed regimen for each of the groups. Groups 3 and 4 mice received a regimen including ChAdOx1-(C1C62) and MVA-delivered (M3M4) HIVconsvX T cell immunogens eight weeks apart (C8M). **c,d** Ratios between trimer (SOSIP.664 or SOSIP.v8)-binding and gp120-binding titers, measured in mice **(c)** and rabbits **(d)**. Median values are represented by horizontal lines. Statistical differences between groups **(c,d)** were determined using a two-tailed Mann-Whitney U test (*p < 0.05, **p < 0.01, ***p < 0.001, ****p < 0.0001).

To determine the quality of the antibody response, we measured the binding of the sera to three different BG505 antigens (gp120, SOSIP.664 and SOSIP.v8) by D7324-capture ELISA **(Fig. S8a,S9a).** The gp120 monomers more readily expose non-NAb epitopes than SOSIP.664 and SOSIP.v8^22,86–88^. SOSIP.664 trimers still display the V3-loop in ELISA^22,86^, while SOSIP.v8 trimers show negligible V3-targeting Ab reactivity^87,88^ due to the presence of stabilizing mutations^64,65^. Therefore, a higher proportion of anti-trimer (SOSIP.664- and SOSIP.v8) binding is indicative of higher-quality Ab responses. Both in mice and rabbits, SOSIP.664 and TTT (groups 1 and 2) elicited similar anti-trimeric binding titers **(Fig. S8a,S9a).** However, SOSIP.664 elicited stronger anti-gp120 responses than TTT, consistent with undesired species participating in the SOSIP.664-induced immune response. Additionally, TTT-primed animals (group 2) elicited higher titers against the more stabilized SOSIP.v8 trimers than those primed with SOSIP.664 (group 1), especially after the SOSIP.664 protein boost. While adding C8M to the TTT regimen (group 4 versus group 2) did not significantly affect the binding responses **(Fig. S8a)**, it did result in the induction of Gag-Pol T cell responses **(Fig. S8b,c).** In rabbits, priming with viral vector and protein co-immunization (group 4) induced the highest and most consistent binding titers to all three antigens **(Fig. S9a).**

Next, we calculated the ratios between the binding titers to each trimeric protein (SOSIP.664 or SOSIP.v8) and gp120 monomer **(Fig. 3c,d).** At week 10, the SOSIP.664:gp120 and SOSIP.v8:gp120 ratios were ∼33-fold (p<0.0001) and ∼574-fold (p<0.0001) higher, respectively, for TTT-primed (group 2) than for SOSIP.664-primed (group 1) mice **(Fig. 3c)**. In rabbits, week 26 ratios were ∼3.5-fold (p<0.0001, SOSIP.664:gp120) and ∼20-fold (p<0.0001, SOSIP.v8:gp120) higher for group 2 than for group 1 **(Fig. 3d)**. This confirms that priming with TTT, which expresses less undesired monomeric and dimeric species, steers responses away from gp120-specific non-NAb epitopes and towards epitopes that are better presented on Env trimers. Of note, mice primed with viral-vectored TTT and T cell immunogen co-immunization (group 4) presented equal or higher trimer-to-gp120 ratios than those that received three doses of SOSIP.664 protein (group 3) **(Fig. 3c)**, which is further testament of the ability of TTT to elicit more trimer-specific responses.

We measured the autologous NAb titers against a BG505/T332N pseudovirus in a standard Tzm-bl assay. None of the mice showed autologous NAb activity **(Fig. S8d)**, consistent with earlier reports that mice do not easily generate NAbs against BG505 trimers^89^. In rabbits, despite the higher trimer-specific binding responses, BG505 TTT elicited autologous NAb responses similar to those elicited by SOSIP.664 **(Fig. S9b).** An explanation might be that the dominant autologous BG505 NAb epitopes are present on gp120 and BG505 gp120 can in some cases induce BG505 NAbs^22^. Furthermore, we did not detect cross-neutralizing responses in the two rabbits (37194 and 37212) that developed the highest autologous NAb titers at week 42 **(Fig. S9c).**

In summary, BG505 TTT efficiently elicits trimer-focused antibody responses in both mice and rabbits. In rabbits, the BG505 TTT-induced responses are able to neutralize the parental BG505 virus.

### Glycans effectively mask immunodominant epitopes on BG505 TTT

BG505 *env* lacks conserved PNGS on positions 241 and 289, which creates a large immunodominant glycan hole that attracts unwanted strain-specific Ab responses^90–94^. Therefore, we generated a <u>g</u>lycan hole <u>m</u>asked (GM) BG505 TTT construct (TTT.GM) that includes potential N-linked glycosylation sites (PNGS) at positions 241 and 289 **(Fig. S10a).** The introduction of these PNGS did not affect the overall properties of the TTT.GM construct, which expressed only as trimers with conformation, thermostability and antigenicity comparable to those of the original TTT **(Fig. S10b-f, Table 1).** Site-specific glycan analysis revealed that the newly incorporated N241 and N289 sites in TTT.GM were efficiently populated by oligomannose-type glycans **(Fig. S10g, Table S9)**, which explained the complete abrogation of the binding of a 241/289-targeting NAb (10A^92^) **(Fig. S10e).**

All soluble Env proteins, including BG505 TTT, present a base glycan hole that constitutes an immunodominant neo-epitope and attracts a high proportion of undesired non-NAb responses^89–91^. In the context of TTT.GM, we attempted to cover the trimer base with glycans by adding PNGS in the TTT linkers, but were so far unsuccessful as these sites were largely unoccupied **(Fig. S10g, Table S9)** and RM19B1 and RM20A2^95^ base-targeting Abs were still reactive **(Fig. S10e)**. The presence of the SC and TTT linkers did, however, block the epitopes of two other base-targeting Abs (RM19R, RM20G^95^) **(Fig. S10e)**.

To evaluate the influence of the 241/289 glycan masking strategy on the immunogenicity of TTT constructs, we immunized rabbits with BG505 TTT.GM following a CMP schedule **(Fig. S11a).** We found no significant differences in binding responses and trimer-to-gp120 ratios between the TTT and TTT.GM groups **(Fig. 11b,c, Table S7).** However, none of the rabbits vaccinated with BG505 TTT.GM developed detectable NAb titers against the parental BG505 pseudovirus **(Fig. 11d, Table S8),** consistent with the efficient masking of the immunodominant 241/289 glycan hole.

Thus, we demonstrate that the TTT design can be adapted to accommodate glycans that shield some of the undesired epitopes on soluble native-like Env trimers.

### The TTT design can be applied to influenza hemagglutinin

Since class I viral fusion glycoproteins share a similar architecture^96^, we hypothesized that some of them, other than Env, might be amenable for the TTT design. A structural screening revealed that the location of the N- and C-termini of the influenza hemagglutinin (HA) protomers is analogous to that of HIV-1 Env. Therefore, we generated a proof-of-concept HA TTT construct encoding three NL03 (A/Netherlands/213/2003(H3N2)) protomers connected by TTT flexible linkers and with a C-terminal StrepII-Tag. The distance between the N- and C-termini of neighboring HA protomers is ∼23 Å **(Fig. 4b)**, i.e., substantially longer than the ∼11 Å for HIV-1 Env **(Fig. 1b)**. Consequently, we designed a longer 22-residue TTT linker to connect the different HA protomers **(Fig. 4a,b).** As controls, we also generated the corresponding HA and HA fused to a GCN4 trimerization domain (HA GCN4) **(Fig. 4a).** The HA TTT construct was predicted by AlphaFold2 to acquire a conformation similar to the experimentally-determined structure of a native-like H3 HA trimer **(Fig. S12).**

**Fig. 4.**
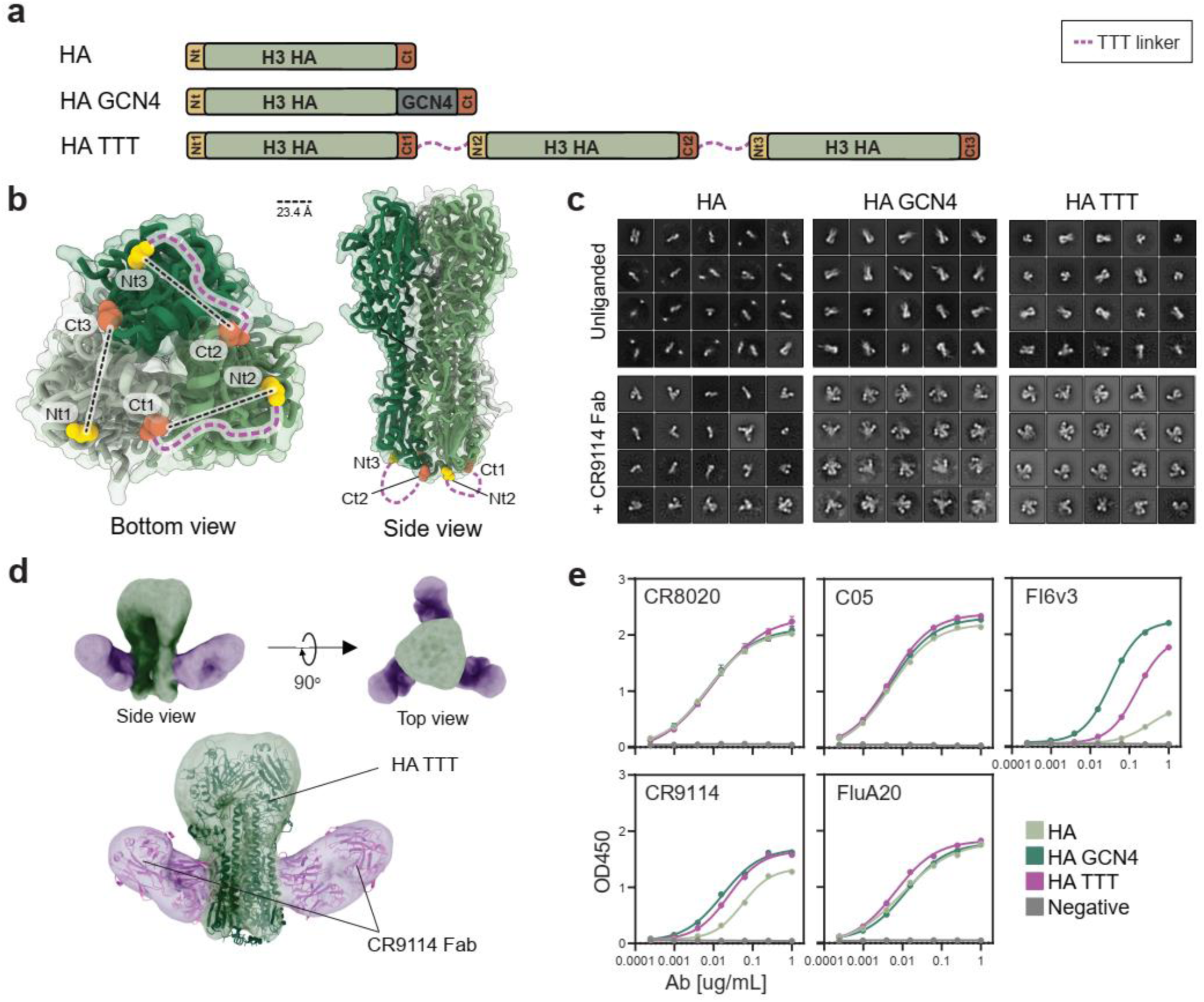
Design and biophysical characterization of a triple tandem trimer HA construct. **a** Linear representation of standard (HA), GCN4-trimerized (HA GCN4) and triple tandem trimer (HA TTT) HA constructs. The TTT construct is composed of three HA protomers fused by TTT linkers (discontinuous purple lines). The N- and C-termini of each protomer are colored in yellow and red, respectively. **b** Expected position of TTT linkers on the structure of a HK05 trimer (A/Hong Kong/4443/2005(H3N2), PDB 2YP7). The distance, in angstrom (Å), between the C_ɑ_ atoms of N-(Asn8, PDB numbering) and C-termini (Lys503) of neighbor protomers are indicated. **c** 2D class averages generated by nsEM analysis of the StrepTactinXT-purified H3 HA, HA GCN4 and HA TTT protein preparations, both unliganded and in complex with a CR9114 Fab (+CR9114 Fab). **d** nsEM-generated 3D model of the purified HA TTT protein in complex with CR9114 Fab, independently (up) and superimposed with a previously determined structure of a similar HK68 trimer (A/Hong Kong/1/1968(H3N2)) + CR9114 complex (PDB 4FQY) (down). **e** StrepTactinXT ELISA with StrepTactinXT-purified HA, HA GCN4 and HA TTT protein preparations against a panel of head- (C05, FluA-20) and stem-specific (CR8020, CR9114, FI6v3) antibodies. Dots and error bars represent the average and standard deviation of values obtained in two independent experiments.

The three HA constructs were expressed in HEK293F suspension cells and purified by StrepTactinXT affinity chromatography. HA TTT showed somewhat lower purification yields than HA and HA GCN4 (∼3.0 mg/L versus ∼7.6 mg/L and ∼6.5 mg/L, respectively) **(Table 1).** In SEC, both HA TTT and HA GCN4 eluted as a single peak at a volume consistent with the size of HA trimers, while HA eluted as a single peak at a higher volume, indicating the presence of non-trimeric HA forms **(Fig. S13a).** The bands on SDS-PAGE confirmed that HA TTT consisted of three covalently linked HA protomers **(Fig. S13b).** By nsEM, purified HA TTT and HA GCN4 appeared homogeneous and contained only trimers, while HA appeared as a heterogeneous mix of monomers, dimers and some trimers **(Fig. 4c).** However, the size and shape of HA trimers complicate the reliable quantification of monomers, dimers and trimers. We therefore complexed the three proteins with Fabs of the bNAb CR9114^97^ **(Fig. 4c),** which binds a conserved conformational epitope on the HA stem. nsEM-generated 2D class averages of these complexes revealed that all HA TTT and HA GCN4 particles presented three Fabs bound, as expected for trimers, while most HA particles had a single Fab bound, indicating that monomers were the predominant species. A nsEM-generated 3D model confirmed that HA TTT-CR9114 complexes assume a conformation consistent with a previously determined high-resolution structure of a native-like H3 HA trimer in complex with CR9114 Fabs^97^ **(Fig. 4d)**.

To assess the antigenicity of HA TTT, we evaluated the binding of several bNAbs, including the head-specific C05^98^ and FluA-20^99^, and the stem-specific CR8020^100^, CR9114^97^ and FI6v3^101^ **(Fig. 4e).** FI6v3 contacts a quaternary epitope located in the stem region between two neighboring protomers Thus, we considered that FI6v3 and CR9114 binding are appropriate proxies for proper folding and resistance to unwanted post-fusion conversion. While the binding of CR8020, C05 and FluA-20 was similar for the three proteins, FI6v3 and CR9114 showed stronger binding to HA TTT and HA GCN4 than to the HA protein **(Fig. 4e).** These findings suggest that HA TTT not only forms trimers, but also trimers that are folded in the proper conformation. The nanoDSF-determined T_m_ value of HA TTT (54.5 °C) was substantially higher than that of HA (49.0 °C), but lower than that of HA GCN4 (58.8 °C) **(Fig. S13c, Table 1).** Furthermore, site-specific glycan analysis revealed that the glycosylation of HA TTT corresponds well with that of a native-like H3 HA trimer analyzed previously **(Fig. S13d, Table S10)**^102,103^.

Overall, we show that the HA TTT format yields high quality and stable HA trimers and demonstrates the broad application of the TTT design for class I fusion proteins.

### The TTT platform allows for expression of chimeric Env immunogens

We hypothesized that the TTT design would facilitate the controlled generation of chimeric immunogens that contain three different protomers in a single Env trimer. As proof of concept, we generated a chimeric Env TTT (chEnv TTT) construct encoding single-chain BG505, ConM and AMC011 protomers connected by 11-residue TTT flexible linkers **(Fig. 5a,b)**. AlphaFold2 predicted that the chEnv TTT sequence would fold with a structure comparable to native-like BG505, ConM and AMC011 Env trimers **(Fig. S14).**

**Fig. 5.**
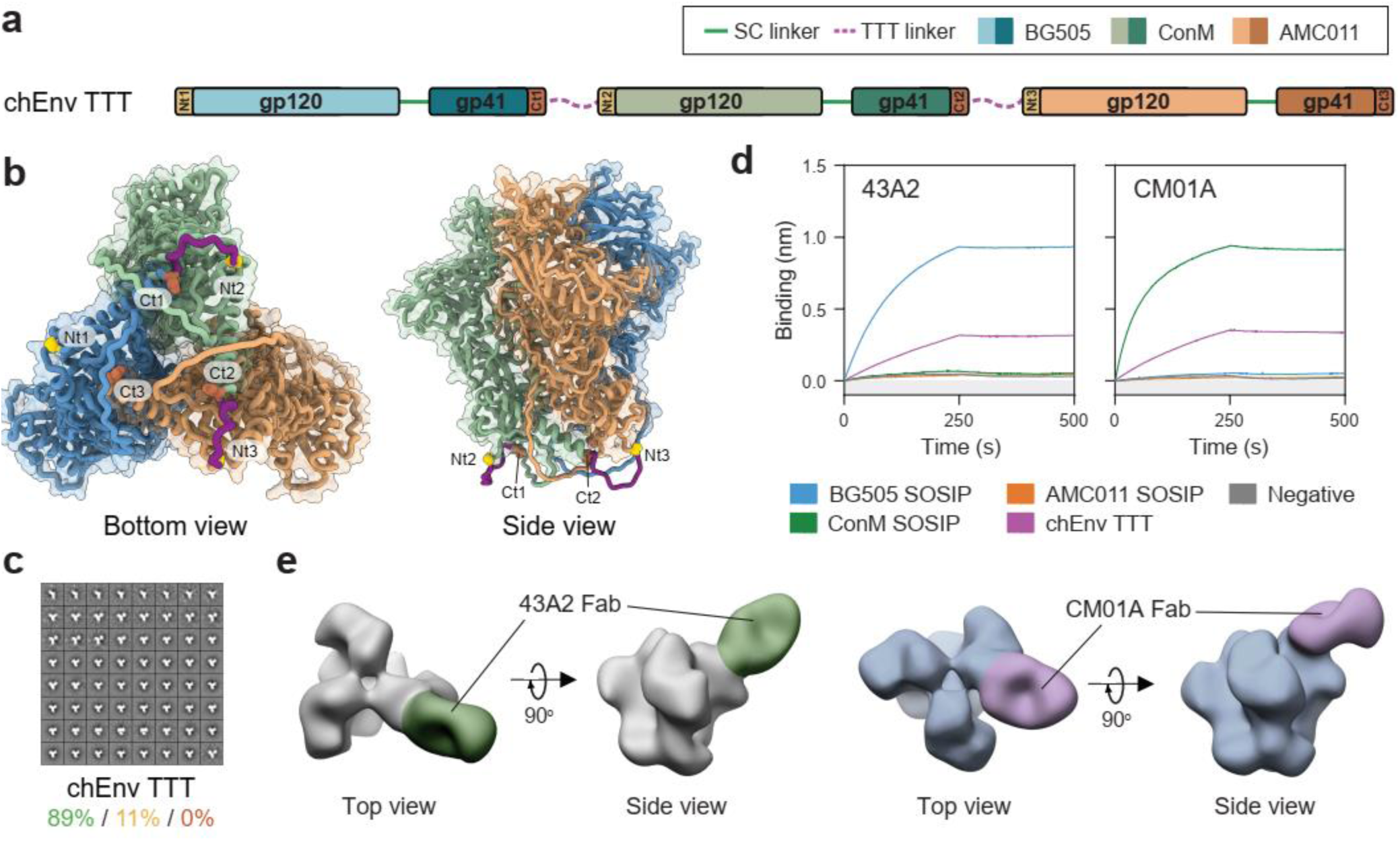
Design and biophysical characterization of a chimeric Env TTT (chEnv TTT) construct. **a** Linear representation of the chEnv TTT construct, composed of single-chain BG505, ConM and AMC011 protomers connected by TTT linkers (purple discontinuous lines). N- and C-termini of each protomer are colored in yellow and red, respectively. **b** Position of TTT linkers on the AlphaFold2-predicted structure of chEnv TTT **(Fig. S14). c** nsEM-generated 2D class averages of the PGT145-purified chEnv TTT protein. Percentages of native-like trimers (green), malformed trimers (yellow) and monomers/dimers (red) are indicated. **d** ProtA BLI assay with PGT145+SEC-purified chEnv TTT protein and BG505-(43A2) and ConM-specific (CM01A) antibodies. PGT145-purified BG505 SOSIP.v5.2, ConM SOSIP.v7 and AMC011 SOSIP.v9 were included as homotrimeric controls. The experiment was performed in duplicate, and the curves shown correspond to one of these repetitions. **e** nsEM-generated 3D models of the chEnv TTT protein in complex with 43A2 and CM01A Fabs.

Subsequently, we expressed chEnv TTT in HEK293F suspension cells and purified the resulting protein by PGT145 affinity chromatography, obtaining a yield of 0.4 mg/L **(Table 1)**. The purified protein preparation contained 89% of covalently linked native-like trimers and no monomers or dimers **(Fig. 5c, Fig. S15a)** and presented a nanoDSF-measured T_m_ of 69.3 °C **(Fig. S15b, Table 1)**. This T_m_ is lower than for BG505 TTT (76.9 °C), but comparable to those previously reported for BG505 SOSIP.v4.2 (70.7 °C)^104^, ConM SOSIP.v7 (67.8 °C)^73^ and AMC011.v5.2 (67.0 °C)^105^. After using SEC to obtain a homogeneous trimer preparation depleted of aggregates **(Fig. S15c)**, a BLI assay was performed to assess the antigenicity of chEnv TTT **(Fig. 5d, Fig. S15d).** The antibody panel included bNAbs (2G12, VRC01, PGT121, PGT145, VRC026.25, PG16, PGT151) and non-NAbs (F105 and 19b), but also BG505-(43A2)^106^ and ConM- (CM01A)^107^ specific antibodies. PGT145-purified BG505 SOSIP.v5.2, ConM SOSIP.v7 and AMC011 SOSIP.v9 trimers were included as homotrimeric controls. chEnv TTT was reactive to all bNAbs tested and did not bind non-NAbs F105 and 19b. Except for PGT151, whose binding is affected by the presence of the SC and TTT linkers, and 2G12, which was used as a loading control, all bNAbs showed a binding signal that accounted for approximately the average of the binding signals observed for the three homotrimeric controls **(Fig. S15d),** consistent with a well-folded chimeric protein. Furthermore, the 43A2 and CM01A showed reduced (approximately a third) binding to the chEnv TTT protein compared to the BG505 and ConM homotrimeric controls, respectively **(Fig. 5d)**, indicating that only one of the protomers in chEnv TTT is reactive to these strain-specific antibodies and thus validating the chimeric nature. nsEM-generated 3D models of the chEnv TTT in complex with 43A2 and CM01A Fabs confirmed that only one of the protomers of the chimeric protein was reactive to these strain-specific antibodies **(Fig. 5e)**. Site-specific glycan analysis of chEnv TTT revealed a glycosylation profile **(Fig. S15e, Table S11)** comparable to BG505 TTT **(Fig. S4d)** and other native-like Env trimers.

### The TTT platform allows for expression of chimeric influenza HA immunogens

In the influenza field, the combination of immunogens from diverse viral strains is well established^108–110^. For instance, most licensed seasonal influenza vaccines are quadrivalent, i.e., contain attenuated/inactivated viral particles or recombinant HA proteins from four different strains to increase the protective coverage^111^. Furthermore, the location of the most conserved epitopes on the stem region of the HA protein has led to the design of head-chimeric proteins with head and stem regions from different viral strains^112^. The combination of several proteins with the same stalk but different head regions results in broad stalk-specific immune responses^112^.

We designed TTT constructs encoding both protomer- (cH125 TTT) and head-chimeric (cH125/111 TTT) HA trimers. The cH125 TTT construct encoded three HA protomers of influenza viruses from different group 1 subtypes, namely NL09 (A/Netherlands/602/2009(H1N1)), SG57 (A/Singapore/1/1957(H2N2)) and IN05 (A/Indonesia/5/2005(H5N1)) **(Fig. 6a,b).** On the other hand, the head chimeric cH125/111 TTT construct contained three protomers with identical PR8 (A/Puerto Rico/8/34/Mount Sinai(H1N1)) stems but different NL09, SG57 and IN05 heads **(Fig. 6e,f)**. Similar to the H3 HA TTT construct, protomers were connected via 22-residue flexible linkers. AlphaFold2 predicted both chimeric constructs could acquire a conformation similar to the experimentally determined structure of native-like H1, H2 and H5 HA trimers **(Fig. S15,S16).** As controls, we generated monomeric constructs encoding the first protomer of both cH125 TTT and cH125/111 TTT, i.e., monomeric NL09 (H1) **(Fig. 6a)** and monomeric NL09-head and PR8-stem (H1/1) **(Fig. 6e).** Additionally, we generated GCN4-trimerized controls encoding each of the three protomers of both chimeras, named H1 GCN4, H2 GCN4 and H5 GCN4 **(Fig. 6a)** and H1/1 GCN4, H2/1 GCN4 and H5/1 GCN4 **(Fig. 6e)**, respectively.

**Fig. 6.**
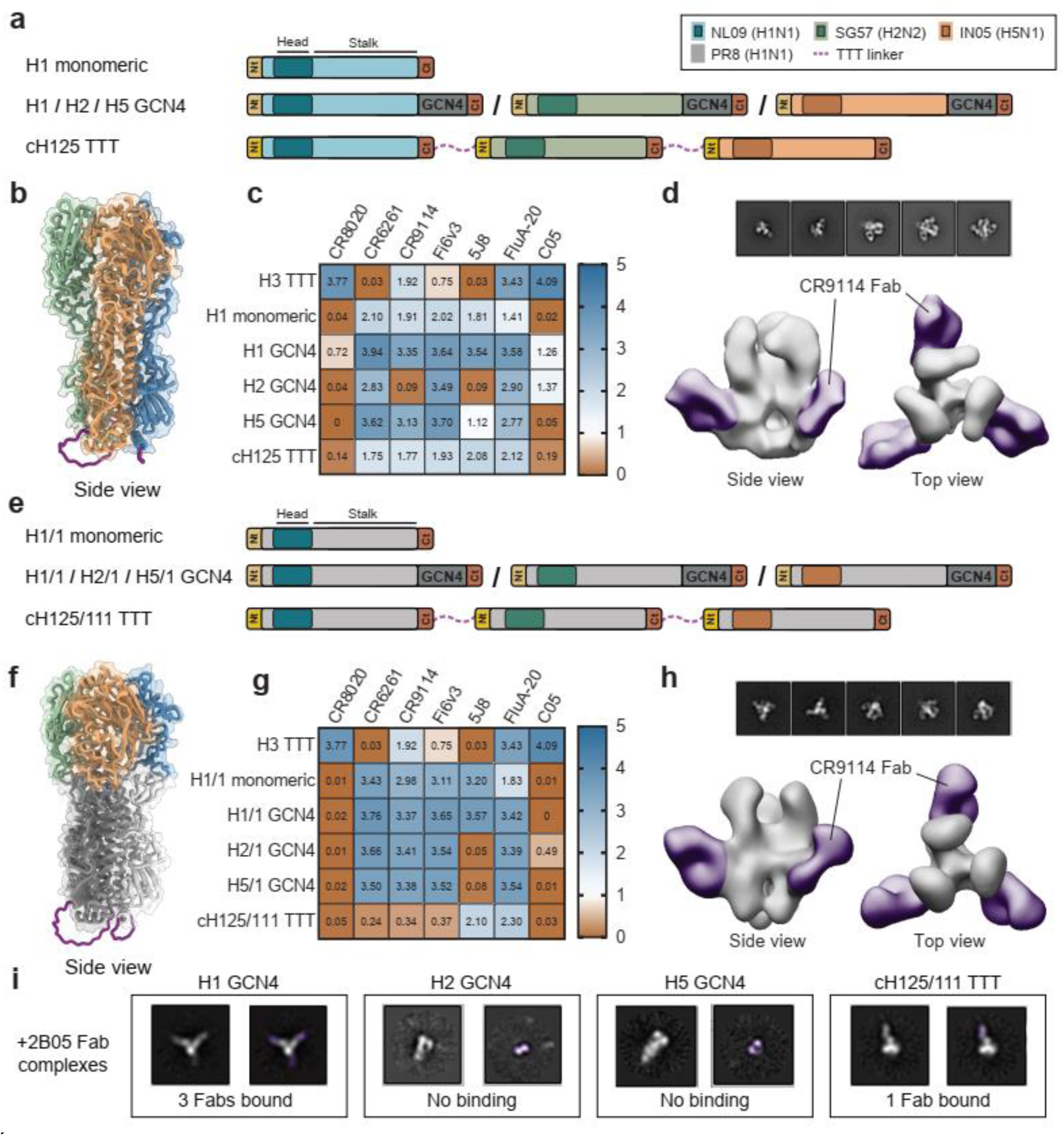
Design and biophysical characterization of chimeric HA TTT constructs. **a,e** Linear representations of chimeric cH125 TTT **(a)** and cH125/111 TTT **(e)** constructs, and the corresponding monomeric and GCN4-trimerized controls. The cH125 TTT construct is composed of NL09 (A/Netherlands/602/2009(H1N1)), SG57 (A/Singapore/1/1957(H2N2)) and IN05 (A/Indonesia/5/2005(H5N1)) protomers connected by TTT linkers (purple discontinuous lines). The cH125/111 TTT construct is composed of protomers with NL09, SG57 and IN05 heads and PR8 (A/Puerto Rico/8/1934/Mount Sinai(H1N1)) stalks. N- and C-termini of each protomer are colored in yellow and red, respectively, and the GCN4 trimerization motifs are colored in dark grey. **b,f** AlphaFold2-predicted structure of cH125 TTT **(b)** and cH125/111 TTT **(f) (Fig. S16,S17). c,g** StrepTactinXT ELISA with StrepTactinXT-purified cH125 TTT **(c)** and cH125/111 TTT **(g)** proteins against a panel of HA head- (5J8, FluA-20, C05) and stalk-specific (CR8020, CR6261, CR9114, FI6v3) antibodies. Heatmap values correspond to the areas under the curve (AUC) of the binding curves in **Fig. S19b,d. d,h** nsEM-generated 3D models of the purified cH125 TTT **(d)** and cH125/111 TTT **(h)** proteins in complex with CR9114 Fabs. **i** nsEM-generated 2D class averages of the homotrimeric H1 GCN4, H2 GCN4 and H5 GCN4 proteins and the chimeric cH125/111 TTT protein in complex with Fabs of the H1-specific 2B05 antibody (in purple).

First, we performed a StrepTactinXT ELISA assay with the supernatant of HEK293T cells transfected with cH125 TTT, cH125/111 TTT and the corresponding controls, including also the H3 HA TTT homotrimeric construct **(Fig. S18).** We evaluated the binding of both stem- (CR8020^100^, CR6261^113^, CR9114^97^, FI6v3^101^) and head-specific (5J8^114^, FluA-20^99^, C05^98^). All samples showed protein expression and, overall, the antigenic profile matched the specificities of the tested antibodies. CR8020 is a group 2-specific antibody^100^ and thus bound only to the H3 HA TTT sample. CR6261 is, on the contrary, a group 1-specific antibody^113^ and was reactive to the H1, H2 and H5 samples, but no to the H3 HA TTT control. Antibodies that are reactive to most influenza A viruses, such as CR9114^97^, FI6v3^101^ and FluA-20^99^, interacted well with all supernatant samples. 5J8 interacted only with constructs encoding H1 heads, as expected^114^. C05 has been reported to bind certain H1, H2, H3 heads^98^, but might not bind the specific H1 head (NL09) that we used in this case. The lower binding signals observed for CR6261, CR9114 and FI6v3 against cH125 TTT and cH125/111 TTT might be due to the stem breathing observed using nsEM **(Fig. 6d,h)**

Next, we purified both chimeric HA TTT proteins and corresponding controls from transiently transfected HEK293F cells using StrepTactinXT affinity chromatography. As in the case of H3 HA TTT construct, both cH125 TTT (∼0.2 mg/L) and cH125/111 TTT (∼1.2 mg/L) presented lower purification yields than the monomeric and GCN4-trimerized controls **(Table 1).** However, both chimeric constructs expressed only covalently linked trimers, as evidenced by the single band of ∼250 KDa observed on reducing SDS-PAGE **(Fig. S19a).** Furthermore, StrepTactinXT ELISA assay with the purified protein preparations showed an antigenic profile matching the one observed with unpurified supernatants **(Fig. 6c,g, Fig. S19b,d).** Also, the nanoDSF-measured T_m_ values of cH125 TTT (55.9 °C) and cH125/111 TTT (58.7 °C) were comparable to H3 HA TTT (54.5 °C) and monomeric (56.6 °C and 56.7 °C for H1 and H1/1, respectively) and GCN4-trimerized controls (57.6 °C, 55.9 °C, 57.2 °C, 51.3 °C and 61.0 °C for GCN4-trimerized H1, H5, H1/1, H2/1 and H5/1, respectively) **(S19c,e, Table 1).** Site-specific glycan analysis of cH125 TTT and cH125/111 TTT revealed that most PNGS were effectively populated by glycans **(Fig S19f, Table S12)**, and only the N20/21 site on the first protomer of cH125 TTT and the N483 sites of all three protomers of cH125/111 TTT presented some unoccupancy. The majority of sites were dominated by complex glycans, probably due to the lower density of glycosylation sites compared to the H3 HA TTT protein^115^.

To assess the conformation of the purified proteins, we imaged them in complex with CR9114 Fabs **(Fig. 6d,h, Table S13)**. nsEM-generated 2D class averages showed that the H1 and H1/1 preparations contained mostly monomers, while the chimeric TTT and GCN4-trimerized preparations contained only trimers with three CR9114 Fabs bound **(Fig. 6d,h, Table S13).** Reconstructed 3D models of these complexes showed that the head domains of both cH125 TTT and cH125/111 TTT appeared in an open conformation, exposing the conserved trimer interface region, which has been suggested to be a target of protective Abs^116^. To prove the chimeric nature of the cH125/111 TTT protein, we imaged it in complex with Fabs of the H1 head-specific 5J8 and 2B05^117^ antibodies. The resulting complexes were unstable and difficult to image. However, we were able to obtain 2D class averages in which H1 GCN4 presented three 2B05 Fabs bound, while cH125/111 TTT presented only one Fab bound, indicating that this latter protein contained only a single H1 head **(Fig. 6i, Table S13).**

In summary, the TTT design can be used for the controlled generation of chimeric Env and HA TTT constructs that express only as trimers and present the expected binding of broad as well as strain-specific antibodies.

## Discussion

Expression of many recombinant class I fusion glycoprotein constructs yields a heterogeneous mix of trimers, dimers and monomers, complicating their use in genetic vaccination approaches including those based on viral vectors and mRNA. Here, we describe a novel design, termed ‘Triple Tandem Trimer’ (TTT), and validate it for the generation of HIV-1 Env and influenza HA immunogens that express only as trimers.

Class I viral glycoprotein trimers, such as HIV-1 Env, influenza HA, SARS-CoV-2 S protein and RSV F protein, generally form dimers/monomers or adopt a post-fusion conformation when their transmembrane domain is removed. Therefore, heterologous trimerization domains, such as the GCN4 leucine zipper^52,53,118^, the bacteriophage T4 fibritin foldon^54,55^ and the molecular clamp^119^ domains, are used to generate soluble versions of these glycoprotein trimers that acquire stable pre-fusion trimeric conformation. However, even with such trimerization domains, not all resulting glycoprotein molecules assemble as trimers^59^. Furthermore, these trimerization domains might negatively affect the conformation^63^ or antigenicity^117^ of the trimerized proteins. Lastly, their presence results in the elicitation of undesirable trimerization domain-specific antibody responses^59^. For example, the clamp trimerization domain is based on HIV-1 gp41 and was used to generate a SARS-CoV-2 spike vaccine, which elicited anti-clamp antibodies that cross-reacted with widely used HIV diagnostics when tested in a phase I trial^120^. Therefore, the development of this vaccine candidate was halted. The nature of the TTT design largely circumvents these various disadvantages and promotes the formation of stable trimers.

Protein fusion is a method widely used in the generation of recombinant proteins, as it allows for the merging of several functionalities and/or properties in a single molecule^121–125^. In most of these applications, the fused proteins are originally encoded by different genes and do not interact to form multimers. Moreover, the flexible linkers usually act as mere linkages that provide the spacing necessary for independent folding. The TTT fusion strategy here described is one of the first examples in which the fused proteins are protomers of a multimeric protein and closely interact and depend on each other for proper folding and conformation. This strategy could be used in the future to assure the multimerization of any multimeric protein, provided that the N- and C-termini of neighbor protomers are relatively close to each other (like in Env and HA) or the addition of longer (and probably rigid) linkers is possible and does not affect the function or conformation of the protein.

Env and HA TTT constructs efficiently express as well-folded trimers that display fewer potentially distracting epitopes than conventional designs, and therefore represent promising candidates for gene-based vaccination. In this study, we provide proof of concept for the use of Env TTT using adenovirus and poxvirus vectors. The combination with other state-of-the-art and emerging delivery technologies, such as mRNA-LNPs^43^, self-amplifying RNAs^46,47,126^ and integrase-deficient lentivirus vectors (IDLVs)^48,49^, might further improve the induction of NAb responses.

The glycan masking of the immunodominant BG505-specific 241/289 glycan hole is an established strategy to reduce the otherwise dominant BG505 strain-specific hole-directed NAb responses^70,127^. In the present study, rabbits vaccinated with 241/289-masked (TTT.GM) immunogens did not develop detectable BG505 NAb responses, consistent with the effective silencing of the 241/289 glycan hole. This modification could therefore facilitate immunofocusing to more desirable sites in epitope-focused and germline-targeting vaccine approaches^128–131^.

The combination of spike proteins from different viral strains or that target different steps of bNAb evolution is a promising approach for the generation of bNAbs against HIV-1 and influenza^4^. However, the diverse proteins are usually administered sequentially, in cocktail or as mosaic nanoparticles^109,132–134^. The chimeric Env and HA TTT immunogens here described can include three different immunogens in every single trimer, which might influence the immune profile compared to other combination vaccines. Furthermore, chimeric TTT containing protomers to target several steps of bNAb evolution might help to simplify sequential-based germline-targeting vaccination strategies, as well as accelerate the screening, approval and manufacture of novel immunogens.

As gene-based vaccination approaches have recently gained much momentum, in part because of the success of mRNA and viral vector COVID-19 vaccines, methods that ensure the expression of predominantly correctly folded antigens from these platforms should facilitate capitalizing on this momentum for HIV-1, influenza and other pathogens. The TTT platform might therefore be particularly suitable for next generation gene-based vaccines aimed at induction of bNAbs.

## Methods

### Construct design

The BG505 SOSIP.664 construct (SOSIP.664), derived from the BG505 *env* gene (Genbank ADM31403.1), has been described elsewhere^86^. Briefly, SOSIP.664 constructs include a set of amino acid changes (HxB2 numbering) to improve the expression and stability of soluble Env trimers: a tPA signal peptide (MDAMKRGLCCVLLLCGAVFVSPSQEIHARFRRGAR); 332N in gp120 (introduction of epitopes of N332 glycan-dependent Abs); 501C-605C (gp120-gp41_ECTO_ disulfide bond); 559P in gp41_ECTO_ (trimer-stabilizing); REKR to RRRRRR (R6) in gp120 (furin cleavage enhancement); and a stop codon after residue 664 in gp41_ECTO_.

The BG505 SOSIP.v8.4 gene (referred to as SOSIP.v8 throughout most of this manuscript for simplicity) was derived from the previously described BG505 SOSIP.v4.1 sequence^64^ by introducing several MD39 trimer-stabilizing mutations (106E, 271I, 288L, 304V, 319Y, 519S, 568D, 570H and 585H)^65^ **(Table S1).** To generate the single-chain SOSIP.v8 (SC) construct, the R6 furin motif in SOSIP.v8 was replaced by a 15-residue flexible linker (GGSGGGGSGGGGSGG), similarly to previous studies^35,36^, and a secondary RCKR furin motif was eliminated (500K). The BG505 triple tandem trimer construct (BG505 TTT) was obtained by concatenating three SOSIP.v8 SC protomers via 11-residue flexible linkers (GSGGSGGSGSG).

The glycan masked TTT construct (TTT.GM) was obtained by introducing several modifications to the BG505 TTT construct: 1) NxT PNGS motifs in every BG505 glycosylation site, except for 137 and 156 (NxS), in order to increase the overall glycan occupancy^72^; 2) PNGS motifs (241N, 291T) and compensatory mutations (240T, 290E) to close the BG505-specific 241/289 glycan hole that attracts narrow-NAb responses^92^; 3) PNGS motifs by introducing G/S to N changes in the TTT linkers (GSGGSNGSGSG) to mask the immunodominant base glycan hole.

The chimeric Env TTT construct (chEnv TTT) comprises three protomers encoding SC versions of BG505 SOSIP.v4.1, ConM SOSIP.v7 and AMC011 SOSIP.v7, fused via 11-residue TTT linkers (GSGGSGGSGSG). The 241/289 strain-specific glycan hole of the BG505 protomer was masked as described above. Both ConM and AMC011 SOSIP.v7 sequences were designed similarly to previously described^73^, by addition of the SOSIP.v5.2 (73C-561C)^104^ and TD8 mutations (47D, 49E, 65K, 106T, 165L, 429R)^135^ to the ConM and AMC011 SOSIP.v4.2 constructs described elsewhere **(Table S1)**^73,136^.

The HA, HA TTT and HA GCN4 constructs were based on the HA sequence of the NL03 virus isolate (A/Netherlands/213/2003(H3N2), Genbank ADM31403.1), positions 8-503 (H3 numbering). Flexible 22-residue TTT linkers (GSGGSGGSGSGGSGGSGGSGSG) were used to connect the three HA protomers comprising the TTT construct. SC linkers were not required as this HA sequence does not contain peptidase cleavage motifs. The HA GCN4 construct was generated by fusing a GCN4-pII isoleucine zipper trimerization motif (RMKQIEDKIEEILSKIYHIENEIARIKKLIGER)^52,53,59^ to the C-terminus of the HA construct using a flexible spacer linker (<u>GS</u>GGSGGSGGSGGS).

The protomer-chimeric HA TTT construct (cH125 TTT) is composed of NL09 (A/Netherlands/602/2009(H1N1), Genbank CY039527.2), SG57 (A/Singapore/1/1957(H2N2), Genbank L11142.1) and IN05 (A/Indonesia/5/2005(H5N1), Genbank CY116646.1) protomers, while the head-chimeric HA TTT construct (cH125/111 TTT) contains three protomers with identical PR8 (A/Puerto Rico/8/34/Mount Sinai(H1N1), Genbank AAM75158.1) stalks but different NL09, SG57 and IN05 heads. In both constructs, the protomers were connected via 22-residue TTT linkers (GSGGSGGSGSGGSGGSGGSGSG) and included a 98F mutation in the receptor binding site (RBS) region, which prevents non-specific hemagglutination while preserving the epitopes of known RBS-targeting bNAbs^137^. Similar to others, we replaced the QRESRRKKR multibasic cleavage site in the H5N1 sequence with QRETK^138–140^. Additionally, we deleted a furin motif in the H5N1 sequence (599K).

All constructs comprised the above-described sequences preceded by tissue plasminogen activator (MDAMKRGLCCVLLLCGAVFVSPSQEIHARFRRGAR) or synthetic leader (MDRAKLLLLLLLLLLPQAQA) signal peptides. Untagged Env constructs presented a STOP codon after position 664. Strep-tagged constructs included an additional Twin-Strep-Tag amino acid sequence (<u>GS</u>GGSSAWSHPQFEKGGGSGGGSGGSAWSHPQFEKG) after positions 664 (Env) or 503 (HA), except for the GCN4-trimerized constructs, which included a Strep-TagII (GGRSGWSHPQFEK) after the GCN4 trimerization domain. D7324-tagged gp120, SOSIP.664 and SOSIP.v8 proteins used for assessing antibody binding responses were expressed from constructs with a C-terminal D7324 tag (<u>GS</u>APTKAKRRVVQREKR) after position 664. In every case, the underlined <u>GS</u> residues were encoded by a BamHI restriction site useful for cloning purposes.

Genes were codon-optimized and synthesized by Genscript (Piscataway, USA) or Integrated DNA Technologies (Coralville, USA) and cloned by restriction-ligation into a pPI4 plasmid. For TTT constructs, especially the ones intended to be delivered by ChAdOx1 and MVA viral vectors, we applied specific optimization considerations and manually optimized each of the three protomer units differently to avoid repeat-mediated gene instability. Subcloning into ChAdOx1 and MVA vectors was achieved by priorly introducing 5’ KpnI/SmaI and 3’ NotI restriction sites.

### Protein expression

Proteins encoded by the described constructs were expressed either in adherent HEK293T or suspension HEK293F cells, as previously described^64,70^. HEK293T cells (ATCC, CRL-11268) were cultured in Dulbecco’s Modified Eagle’s Medium (DMEM) supplemented with 10% fetal calf serum (FCS), penicillin (100 U/mL), and streptomycin (100 µg/mL); and transfected as previously described^64^. Three days after transfection, HEK293T supernatants were harvested, filtered (0.22 µm pore size) and directly used in supernatant ELISA and BLI assays. Suspension HEK293F cells (Invitrogen, cat no. R79009) were cultured in FreeStyle Expression Medium (Gibco) and, 5-7 days post-transfection, supernatants were harvested, centrifuged and filtered using 0.22 µm Steritops (Millipore, Amsterdam, The Netherlands) before protein purification. For transfection, Env- or HA-encoding plasmids were mixed with PEImax (Polysciences Europe GmBH, Eppelheim, Germany) in a 3:1 (w/w) PEImax to DNA ratio. Co-transfection with a furin protease-encoding plasmid was performed in a 4:1 Env to furin ratio only when explicitly indicated.

### Protein purification and profiling

Untagged proteins were purified by Galanthus Nivalis Lectin (GNL) affinity chromatography. Briefly, filtered HEK293F supernatants containing the expressed proteins were flowed (0.5 -1.0 mL/min) over Econo-Column chromatography columns (Bio-Rad, CA, USA) containing 1-2 mL of agarose bound GNL (Vector Laboratories), at 4 °C. After washing with five column volumes of PBS, proteins were eluted with a 1M α-Methyl-D-mannopyranoside (Acros Organics) pH8 solution.

Supernatants containing proteins with C-terminal Strep-Tag peptides were purified by StrepTactinXT affinity chromatography. First, supernatants were mixed with 1 mL of BioLock Biotin blocking solution (IBA GmbH, Göttingen, Germany) and 110 mL of 10x Buffer W (1 M Tris/HCl, pH 8.0 1.5 M NaCl, 10 mM EDTA) per liter of supernatant. Then, the mixes were incubated for a minimum of 15 min at 4 °C before passing them (0.5 - 1.0 mL/min) through StrepTactinXT Superflow (IBA GmbH, Göttingen, Germany) resin beds packed in Econo-Column chromatography columns (Bio-Rad, CA, USA). Columns were washed with ten column volumes of 1x Buffer W and elution of the retained proteins was performed with 1x BXT elution buffer (IBA GmbH, Göttingen, Germany).

The BG505 TTT protein for X-ray crystallography and the chEnv TTT protein were purified by immunoaffinity chromatography using 2G12 and PGT145 bNAbs, respectively, similarly to previously described^70^. Briefly, unpurified proteins contained in the supernatant of transfected HEK293F cells were immobilized on 2G12 or PGT145 functionalized CNBr-activated Sepharose 4B column (GE Healthcare) by flowing at 1 mL/min or overnight incubation at 4 °C on a roller bank. After washing with three column volumes of 0.5 M NaCl and 20 mM Tris HCl pH 8.0, proteins were eluted with 3 M MgCl_2_ pH 7.5.

All eluted proteins were subsequently buffer exchanged to PBS or TBS by using Vivaspin20 ultrafiltration units (Sartorius, Gӧttingen, Germany) with MWCO 50 kDa (Env) or 10 kDa (HA). Protein concentrations were calculated using A_280_ values measured on a NanoDrop2000 device (Thermo Fisher Scientific) and the molecular weight and extinction coefficient values estimated by the ProtParam Expasy webtool.

Some of the affinity chromatography-purified proteins were run through a Superdex200 Increase 10/300 GL column integrated in an NGC chromatography system (Bio-Rad, CA, USA) to obtain a profile of the content in different species (aggregates, trimers, dimers, monomers) in the protein preparations or to remove non-Env impurities from the GNL-purified proteins used in immunization experiments. A Superose6 Increase 10/300 GL column was used to remove aggregates from the PGT145-purified chEnv TTT protein preparation.

### SDS-PAGE and BN-PAGE

Proteins were run through Novex Wedgewell 4-12% Tris-Glycine (Thermo Fisher Scientific) and NuPAGE 4-12% Bis-Tris (Thermo Fisher Scientific) polyacrylamide gels for their SDS-PAGE and BN-PAGE analysis, respectively^64^. Subsequently, gels were stained with PageBlue Protein Staining Solution (Thermo Scientific) or Colloidal Blue Staining Kit (Life Technologies), respectively. Proteins presented in each gel were processed in parallel.

### Enzyme-linked immunosorbent assay (ELISA)

Lectin-capture ELISA experiments were performed as previously described. Briefly, *Galanthus nivalis* lectin (Vector Laboratories) at 20 µg/mL in 0.1 M NaHCO_3_ pH 8.6 was immobilized on half-area 96-well plates (Corning) by overnight incubation. After blocking with Casein Blocker in PBS (Thermo Fisher Scientific), 100 µL of purified proteins in TBS (1 µg/mL) or unpurified cell supernatants were dispensed in the corresponding wells for protein immobilization by a 2 h incubation at room temperature. Subsequent steps to measure binding of the test antibodies were performed similarly to previously described^70^. Briefly, following a double wash step with TBS to remove unbound proteins, serial dilutions of test primary antibodies in Casein Blocker were added and incubated for 2 h. After 3 washes with TBS, HRP-labeled goat anti-human IgG (Jackson Immunoresearch) diluted 1:3000 in Casein Blocker was added and incubated for 1 h, followed by 5 washes with TBS/0.05% Tween20. A developing solution (1% 3,3’,5,5’-tetramethylbenzidine (Sigma-Aldrich), 0.01% H_2_O_2_, 100 mM sodium acetate and 100 mM citric acid) allowed the colorimetric reaction, which was stopped by the addition of 0.8 M H_2_SO_4_. Finally, color development (absorption at 450 nm, OD_450_) was measured to obtain the binding curves.

StrepTactinXT ELISA assays were performed following the same procedures as lectin-capture assays, with the difference that StrepTactinXT coated microplates (IBA GmbH, Göttingen, Germany) did not require any functionalization or blocking steps prior to protein immobilization.

Endpoint antibody titers from mouse and rabbit sera samples were determined in a D7324-capture ELISA, using half-area 96-well plates (Greiner) precoated with anti-D7324 antibody (Aalto Bioreagents) at 10 µg/mL in 0.1 M NaHCO_3_ pH 8.6 overnight, similarly to previously described^22^. The procedures were similar to the ones described for lectin-capture ELISA, with the exception that primary and secondary antibodies were replaced by sera diluted in TBS/2% skimmed milk/20% sheep serum and an HRP-labeled goat anti-rabbit IgG (Jackson Immunoresearch), respectively.

### Biolayer interferometry (BLI)

An Octet K2 (ForteBio) device was used to measure the binding of antibodies at 30 °C and 1000 rpm agitation. Purified proteins were diluted in BLI kinetics buffer (PBS/0.1% bovine serum albumin/0.02% Tween20) to 100nM. First, test antibodies diluted in kinetics buffer were loaded on protein A sensors (ForteBio) to a final interference pattern shift of 1 nm. To obtain a baseline prior to protein association, sensors were equilibrated in kinetics buffer for 60 s. Association and dissociation of glycoproteins were measured for 300s. Binding data were pre-processed and exported using the Octet software.

BLI experiments on cell supernatants diluted in BLI kinetics buffer (PBS/0.1% bovine serum albumin/0.02% Tween 20) were conducted similarly to described above. However, to avoid non-specific binding to other supernatant components, we loaded antibodies to a binding threshold of 2 nm and replaced the kinetics buffer for a mock transfected supernatant for baseline and dissociation steps. When needed, the baseline step was prolonged to 300 s to allow the signal to level out.

### Nano Differential scanning fluorimetry (nanoDSF)

Protein thermostability was evaluated with a Prometheus NT.48 instrument (NanoTemper Technologies). Proteins at a concentration of 1 mg/mL were loaded to the grade capillaries and the intrinsic fluorescence signal was measured while temperature was increased by 1°C/min, with an excitation power of 40%. The temperatures of melting (T_m_) were determined using the Prometheus NT software.

### Sample preparation and analysis by LC-MS

For protein samples to be analyzed by LC-MS, three separate 50 μg aliquots were denatured for 1 h in 50 mM Tris/HCl, pH 8.0 containing 6 M of urea and 5 mM of dithiothreitol (DTT). Next, the proteins were reduced and alkylated by adding 20 mM iodoacetamide (IAA) and incubated for 1 h in the dark, followed by incubation with DTT to get rid of any residual IAA. The alkylated proteins were buffer-exchanged into 50 mM Tris/HCl, pH 8.0 using Vivaspin columns (3 kDa) and digested separately overnight using trypsin, chymotrypsin (Mass Spectrometry Grade, Promega) or alpha lytic protease (Sigma Aldrich) at a ratio of 1:30 (w/w). The peptides were dried and extracted using C18 Zip-tip (Merck Milipore). The peptides were dried again, re-suspended in 0.1% formic acid and analyzed by nanoLC-ESI MS with an Easy-nLC 1200 (Thermo Fisher Scientific) system coupled to an Orbitrap Fusion mass spectrometer (Thermo Fisher Scientific) using stepped higher energy collision-induced dissociation (HCD) fragmentation (15, 25, 45%). Peptides were separated using an EasySpray PepMap RSLC C18 column (75 µm × 75 cm). A trapping column (PepMap 100 C18 3 μm (particle size), 75 μm × 2 cm) was used in line with the LC prior to separation with the analytical column. The LC conditions were as follows: 275 min linear gradient consisting of 0-32% acetonitrile in 0.1% formic acid over 240 min followed by 35 min of 80% acetonitrile in 0.1% formic acid. The flow rate was set to 200 nL/min. The spray voltage was set to 2.7 kV and the temperature of the heated capillary was set to 40 °C. The ion transfer tube temperature was set to 275 °C. The scan range was 400−1600 m/z. The HCD collision energy was set to 50%. Precursor and fragment detection were performed using Orbitrap at a resolution MS1 = 100,000. MS2 = 30,000. The AGC target for MS1 = 4^5^ and MS2 = 5^4^ and injection time: MS1 = 50 ms MS2 = 54 ms.

### Data processing of LC-MS data

Glycopeptide fragmentation data were extracted from the raw file Byos (Version 3.5; Protein Metrics Inc.). The following parameters were used for data searches in Byonic: The precursor mass tolerance was set at 4 ppm and 10 ppm for fragments. Peptide modifications included in the search include: Cys carbamidomethyl, Met oxidation, Glu pyroGlu, Gln pyroGln and N deamidation. For each protease digest, a separate search node was used with digestion parameters appropriate for each protease (Trypsin RK, Chymotrypsin YFW and ALP TASV) using a semi-specific search with 2 missed cleavages. A 1% false discovery rate (FDR) was applied. All three digests were combined into a single file for downstream analysis. All charge states for a single glycopeptide were summed. The glycopeptide fragmentation data were evaluated manually for each glycopeptide; the peptide was scored as true-positive when the correct b and y fragment ions were observed along with oxonium ions corresponding to the glycan identified. The protein metrics 309 mammalian N-glycan library was modified to include sulfated glycans and phosphorylated mannose species, although no phosphorylated mannose glycans were detected on any of the samples analyzed. The relative amounts (determined by comparing the XIC of each glycopeptide, summing charge states) of each glycan at each site as well as the unoccupied proportion were determined by comparing the extracted chromatographic areas for different glycotypes with an identical peptide sequence. Glycans were categorized according to the composition detected. HexNAc(2), Hex(9−3) was classified as M9 to M3. HexNAc(3)Hex(5−6)Neu5Ac (0-4) was classified as Hybrid with HexNAc(3)Hex(5-6)Fuc(1)NeuAc(0-1) classified as Fhybrid. These glycans constituted the high mannose category. Complex-type glycans were classified according to the number of processed antenna and fucosylation. Complex glycans are categorized as HexNAc(3)(X), HexNAc(3)(F)(X), HexNAc(4)(X), HexNAc(4)(F)(X), HexNAc(5)(X), HexNAc(5)(F)(X), HexNAc(6+)(X) and HexNAc(6+)(F)(X).

### In silico protein modeling

Protein molecular graphics and root-mean-square deviation (RMSD) values were generated with the UCSF ChimeraX 1.4 software^141^. The AlphaFold v2.0 software (DeepMind)^142^ was used to predict protein structures. Protein distance maps were generated with a Python script using the following packages: Bio.PDB^143^ for PDB parsing, NumPy^144^ and pandas^145^ for data processing, and seaborn^146^ and matplotlib^147^ for data visualization.

### Negative-stain electron microscopy (nsEM)

Purified proteins or immune complexes were imaged by nsEM essentially as described elsewhere^117,148^. Immune complexes were prepared by incubating Fabs and Env or HA proteins at greater than a 3:1 molar ratio for 2 h at room temperature. Subsequently, 3 μL of purified proteins or immune complexes (∼0.02 mg/mL) were applied onto glow discharged (20 mA for 30s) carbon-coated 400 mesh copper grids (Electron Microscopy Sciences). After 5 s, they were negatively stained with 2% w/v uranyl formate for 60 s. Samples were imaged using a FEI Tecnai T12 electron microscope operating at 120 keV. Micrographs were collected with Leginon^149^ and single particles were processed using Appion^150^, Relion^151^ and XQuartz. The UCSF Chimera software was used to map the footprints and make the figures. Fabs on 2D class average and 3D model figures were colored using Photoshop.

### Crystallization and data collection

To determine the structure of BG505 TTT, trimers were complexed with PGT124 Fabs and 35O22 single-chain variable fragments (scFvs) (in a 3.5:3.5:1 molar ratio of Fab:scFv:trimer) at room temperature for 30 min. The resulting complexes were digested with EndoH glycosidase (New England Biolabs) at 37 °C for 30 min. Subsequently, the digested complexes were purified by size exclusion using a Superdex200 16/600 column (GE Healthcare) and concentrated to 10.7 mg/mL, before being subjected to crystallization screening at both 4 °C and 20 °C using the high-throughput CrystalMation robotic system (Rigaku) at The Scripps Research Institute (TSRI). High-quality crystals were obtained via sitting drops in 0.1 M sodium acetate, 0.2 M zinc acetate, 11.5 % (w/v) PEG 3000, pH 4.83, at 4 °C. These crystals were harvested with 15 % glycerol as cryoprotectant and were immediately cryo-cooled in liquid nitrogen. The diffraction data were collected at the SSRL12-1 beamlines **(Table S3).**

### Structure determination and refinement

The crystals of BG505 TTT + PGT124 Fab + 35O22 scFv complex diffracted to 5.8 Å. The dataset was indexed, integrated and scaled using HKL2000^152^ in a C2 space group. The structures of BG505 SOSIP.664 gp140 (PDB 5CEZ), PGT124 (PDB 4R26) and 35O22 scFv (PDB 6MTJ) were used as search models for molecular replacement. The complex structure was then built and refined via different iterations of Coot^153^ and Phenix^154^. The final R_cryst_ and R_free_ values were 25.5% and 29.2%, respectively, with 97.8% completeness **(Table S3).** The Fab/scFv residues were numbered according to Kabat et al.^155^ and Env residues using HxB2 numbering.

### Construction of the ChAdOx1- and MVA-vectored vaccines

Vaccine vector ChAdOx1 is derived from simian (chimpanzee) adenovirus isolate Y25 of adenovirus group E^75^. The recombinant ChAdOx1 were generated similarly as described elsewhere^81,85^. Briefly, the SOSIP.664, TTT and TTT.GM constructs were inserted into the E1 locus of the adenovirus genome under the control of the human cytomegalovirus immediate-early promoter, while the adenovirus genome was stably integrated in a bacterial artificial chromosome. The recombinant ChAdOx1 vaccines were rescued by transfection of purified excised genomic DNA into HEK293A T-REX (tetracycline-sensitive repressor of transgene expression used for preparation of ChAdOx1-vectored vaccines) cells (Thermo Fisher Scientific, Waltham, USA). The transgene presence and absence of contaminating empty parental adenovirus in the virus stock was confirmed by PCR, and the virus was titered and stored at -80 °C until use.

To generate recombinant MVA vaccines^81,85^, the SOSIP.664, TTT and TTT.GM constructs were inserted directly into the MVA genome by homologous recombination in chicken embryo fibroblast cells. The expression cassette was directed under the control of the modified H5 poxvirus promoter and into the thymidine kinase locus of the MVA genome. Finally, the co-inserted EGFP marker was removed by trans-dominant recombination to generate markerless MVA vaccines. The virus was plaque purified, expanded, purified on a 36% sucrose cushion, tittered, the expression of the transgene product was confirmed on Western Blot and the virus stock was stored at -80 °C until use.

### Western blot to detect TTT expression

Eighty percent confluent HeLa cells in six-well plates were infected with ChAdOx1.BG505 TTT or MVA.BG505 TTT for 1 h and incubated subsequently at 37 °C, 5% CO_2_ for 24 h before the cells were lysed with 300 µl M-PER Mammalian Protein Extraction Reagent (Thermo Scientific). A volume of 10 µl was resolved on a 3-8% Tris Acetate gel, electroblotted onto nitrocellulose and probed with anti-HIV-1 Env mouse monoclonal antibody ARP3119 (Centre for AIDS Reagents, National Institute for Biological Standards and Control). Secondary antibody was a peroxidase-conjugated goat anti-mouse antibody and detection was by adding a chemiluminescent substrate to peroxidase.

### Mouse immunization

Four groups of six-week-old female BALB/c mice were purchased from Envigo (UK) and housed at the Functional Genomics Facility, University of Oxford. Mice were immunized intramuscularly under general anesthesia either with 10^8^ infectious units (IU) of recombinant ChAdOx1, 5 × 10^8^ plaque-forming units (PFU) of recombinant MVA, or 10 μg of protein adjuvanted with AddaVax (Invivogen)^156,157^. Each pair of HIVconsvX-encoding ChAdOx1 and MVA vaccines (C1C62 and M3M4, respectively) were administered together as half doses so that the combined IU and PFU of each pair constituted a full IU dose for ChAdOx1 and a full PFU dose for MVA.

All mouse immunization procedures and care were approved by the local Clinical Medicine Ethical Review Committee, University of Oxford, and conformed strictly to the United Kingdom Home Office Guidelines under the Animals (Scientific Procedures) Act 1986. Experiments were conducted under project license PP1892852 held by T.H.

### Rabbit immunization

The rabbit immunization experiment was performed by Pocono Rabbit Farm & Lab (Canadensis, USA) in female New Zealand rabbits distributed in seven groups (six rabbits per group). Rabbits were immunized at 3 different timepoints (weeks 0, 8 and 24) with either 5 × 10^10^ IU of recombinant ChAdOx1,10^8^ PFU of recombinant MVA, or 30 μg of PGT145- or GNL-purified Env protein adjuvanted with Squalene emulsion (SE) adjuvant (Polymun, Klosterneuburg, Austria) in a 1:1 Env in PBS to SE ratio (v/v). Each immunization consisted of two intramuscular injections (2 × 250 µL) on both quadriceps of each animal. Immunization procedures complied with the relevant ethical regulations and protocols of the Pocono Institutional Animal Care and Use Committee (IACUC), with approval code 9623.

### Neutralization assays

TZM-bl neutralization assays were performed essentially as described elsewhere^86,158^. A BG505/T332N pseudovirus was used to assess the autologous neutralizing responses. The TZM-bl reporter cell line^159^ was obtained from John C. Kappes, Xiaoyun Wu and Tranzyme Inc. through the NIH AIDS Research and Reference Reagents Program, NIAID, NIH. The ID_50_ values were determined as the sera dilution at which infectivity was inhibited by 50 %.

Mice IgGs were purified from sera of immunized animals essentially as described elsewhere^90^. Briefly, sera were incubated with protein A/G Sepharose resin (GE Healthcare) overnight at 4 °C, at a ratio of 1 mL packed resin per milliliter of undiluted serum. The resin was washed 3 times with 10 column volumes of PBS, before eluting the IgGs with 5-10 volumes of 0.1 M glycine pH 2.5. Samples were immediately neutralized with 1 M Tris-HCL pH 8. Finally, buffer was exchanged to PBS (same volume of the initial serum) using 10 kDa MWCO Vivaspin6 filters (Sartorius, Gӧttingen, Germany) and the resulting preparations were used to measure BG505/T332N neutralization as described above.

### IFN-γ ELISPOT assay

Interferon (IFN)-γ Enzyme-linked ImmunoSpot (ELISPOT) assay was performed as described previously^160^ using the Mouse IFN-γ ELISpot kit (Mabtech, Stockholm, Sweden) according to the manufacturer’s instructions. Immune splenocytes were collected and tested separately from individual mice in triplicate wells. Peptides were used at 2 µg/ml each, and splenocytes at 10^5^ cells/well were added to 96-well high-protein-binding Immobilon-P membrane plates (Millipore, UK) that had been precoated with 5 µg/ml anti-IFN-γ monoclonal antibody (mAb) AN18 (Mabtech). The plates were incubated at 37 °C in 5 % CO2 for 18 h and washed with PBS before the addition of 1 µg/ml biotinylated anti-IFN-γ mAb (Mabtech) at room temperature for 2 h. The plates were then washed with PBS, incubated with 1 µg/ml streptavidin-conjugated alkaline phosphatase (Mabtech) at room temperature for 1 h, washed with PBS, and individual spot-producing units (SFU) were detected as dark spots after a 10-min reaction with 5-bromo-4-chloro-3-idolyl phosphate and nitro blue tetrazolium using an alkaline-phosphatase-conjugate substrate (Bio-Rad, CA, USA). SFUs were counted using the AID ELISpot Reader System (Autoimmun Diagnostika). The frequencies of responding cells were expressed as SFU/10^6^ splenocytes after subtracting the background no-peptide frequencies.

### Phenotyping vaccine-elicited follicular T cells

Splenocytes were stimulated with the peptide pools described in the memory phenotypic assay and supplemented with anti-CD28 and anti-CD49d mAbs (ThermoFisher Scientific) both at 1.0 μg/ml or a tissue culture medium with 1% DMSO as a negative control. The cells were incubated at 37 °C, 5% CO_2_ for 2 h prior to the addition of Brefeldin A and monensin (BD Biosciences). After being left in culture overnight, cells were centrifuged briefly, washed in PBS plus 5% BSA (Sigma-Aldrich) and the pellet re-suspended in 40 µl of CD16/32 with LIVE/DEAD fixable aqua stain (Molecular Probes, Invitrogen). Subsequently, cells were washed and stained by incubation at 4 °C for 30 min with 40 µl of mAb anti-membrane marker mix containing CD4 APC/Cy7 (Biolegend), CD19 FITC, CD14 FITC, CD16 FITC, CD3 PerCP-eFluor710, CD8a eFluor 450, CD44 BV605, CCR7 Alexa Fluor 700, CXCR5 PE/Cy7 and PD-1 PerCP-eFluor610 (all from ThermoFisher Scientific). Cells were permeabilized using Fix/Perm solution (BD Biosciences) for 20 min at 4 °C and then washed with Perm Wash buffer (BD Biosciences) Next, they were stained with a mastermix of anti-cytokine/transcription factor mAbs containing BCL6 PE (ThermoFisher Scientific) at 4 °C for 30 min, washed and resuspended in Perm Wash buffer prior to running on an LSRII flow cytometer (Becton-Dickinson). All antibodies were used at pre-titrated, optimal concentrations and the data are presented after subtracting the background.

### Data representation and statistical analyses

All data representation and statistical analyses were performed using Graphpad Prism 9.1.2. Groups were compared using unpaired two-tailed Mann–Whitney U test.

## Data availability

The data that support the findings in this study are available from the corresponding author upon reasonable request. The coordinates and structure factors for the BG505 TTT structure in complex with PGT124 Fabs and 35O22 scFvs have been deposited in the Protein Data Bank (PDB code 8TGO).

## Acknowledgments

We thank Emma Reiss, Marlies van Haaren and Marit van Gils for providing the CM01A and 10A monoclonal antibodies; Albert Cupo and John Moore for kindly supplying PGT145-purified SOSIP.664 protein; and Michel Nussenzweig, Hermann Katinger, Mark Connors, James Robinson, Dennis Burton, John Mascola, Peter Kwong, and William Olson for donating antibodies and reagents directly or through the NIH AIDS Research and Reference Reagent Program. We thank Dietmar Katinger and Ehsan Suleiman for providing the squalene emulsion adjuvant.

This project has received funding from the European Union’s Horizon 2020 research and innovation program under grant agreement No. 681137 (to R.W.S. and M.C.). This work was also supported by the U.S. National Institutes of Health Grant P01 AI110657 (to A.B.W., I.A.W. and R.W.S.); by the International AIDS Vaccine Initiative (IAVI); by the Bill and Melinda Gates Foundation through the Collaboration for AIDS Vaccine Discovery (CAVD), grants OPP1111923 and OPP1132237 (to R.W.S.) and OPP1115782 (A.B.W.); by the Aids Fonds Netherlands, Grant #2016019 (to R.W.S.); and by the Fondation Dormeur, Vaduz (to R.W.S.). R.W.S. is a recipient of a Vici grant from the Netherlands Organization for Scientific Research (NWO). We thank EMBO for the Short-Term Fellowship (STS-7814) awarded to S. Kumar.

We are grateful to the staff of the Stanford Synchrotron Radiation Lightsource (SSRL) beamline 12-1 for assistance. This research used resources of the SSRL, SLAC National Accelerator Laboratory, which is supported by the U.S. Department of Energy, Office of Science, Office of Basic Energy Sciences under contract no. DE-AC02–76SF00515. The SSRL Structural Molecular Biology Program is supported by the DOE Office of Biological and Environmental Research and by the National Institutes of Health – National Institute of General Medical Sciences (including P41GM103393).

Molecular graphics and analyses performed with UCSF Chimera, developed by the Resource for Biocomputing, Visualization, and Informatics at the University of California, San Francisco, with support from NIH P41-GM103311. AlphaFold2 predictions were obtained using the Lisa Computer Cluster, made available by SURF, Science Park 140, 1098XG Amsterdam, Netherlands. Graphics on Figures 2,S1 and S11 were generated using BioRender.com. The electron microscopy data were collected at Electron Microscopy Facility of The Scripps Research Institute.

## Author contributions

Conceptualization: I.d.M-S., K.S., R.W.S. Conceived and designed the experiments: I.d.M-S., E.G.W, Y.X., W.L., J.D.A., A.T.d.l.P, R.F.R., J.F., A.N.L., E.E.S., J.H., M.C., G.O., A.B.W., I.A.W., T.H., K.S., R.W.S. Performed the experiments: I.d.M-S., E.G.W, Y.X., W.L., J.D.A., A.T.d.l.P, R.F.R., J.F., A.N.L., S.K., E.E.S., J.A.B., S.K., R.Z., M.B., M.Y., G.O., K.S. Analyzed the data: I.d.M-S., E.G.W, Y.X., W.L., J.D.A., A.T.d.l.P, R.F.R., J.F., A.N.L., E.E.S., M.Y., G.O., K.S. Wrote the paper: I.d.M-S., E.G.W, Y.X., J.D.A., T.H., K.S., R.W.S. Edited or revised the paper: All authors commented on the manuscript and approved the final version.

