## Supplementary Material for "Triple tandem trimer immunogens for HIV-1 and influenza nucleic acid-based vaccines"

del Moral-Sánchez et al.

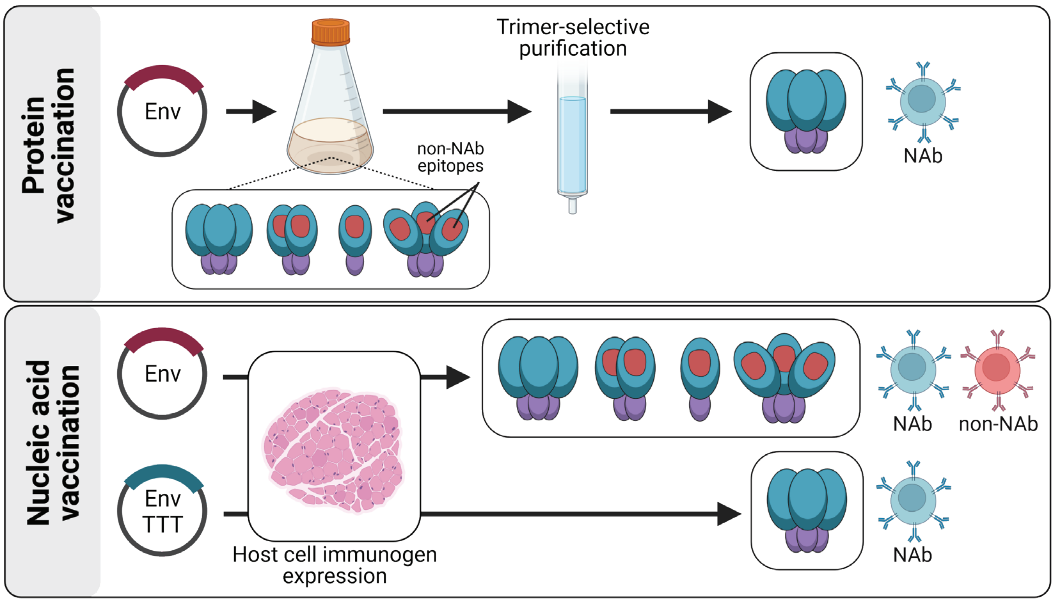


**Fig. S1 Value of constructs that express only as trimers for nucleic acid vaccination approaches.** Compared to other HIV-1 envelope constructs, such as SOSIP and SC (Env, in purple), triple tandem trimer constructs (Env TTT, in blue) express only as trimers, avoiding the participation of inner non-NAb epitopes in the immune response.

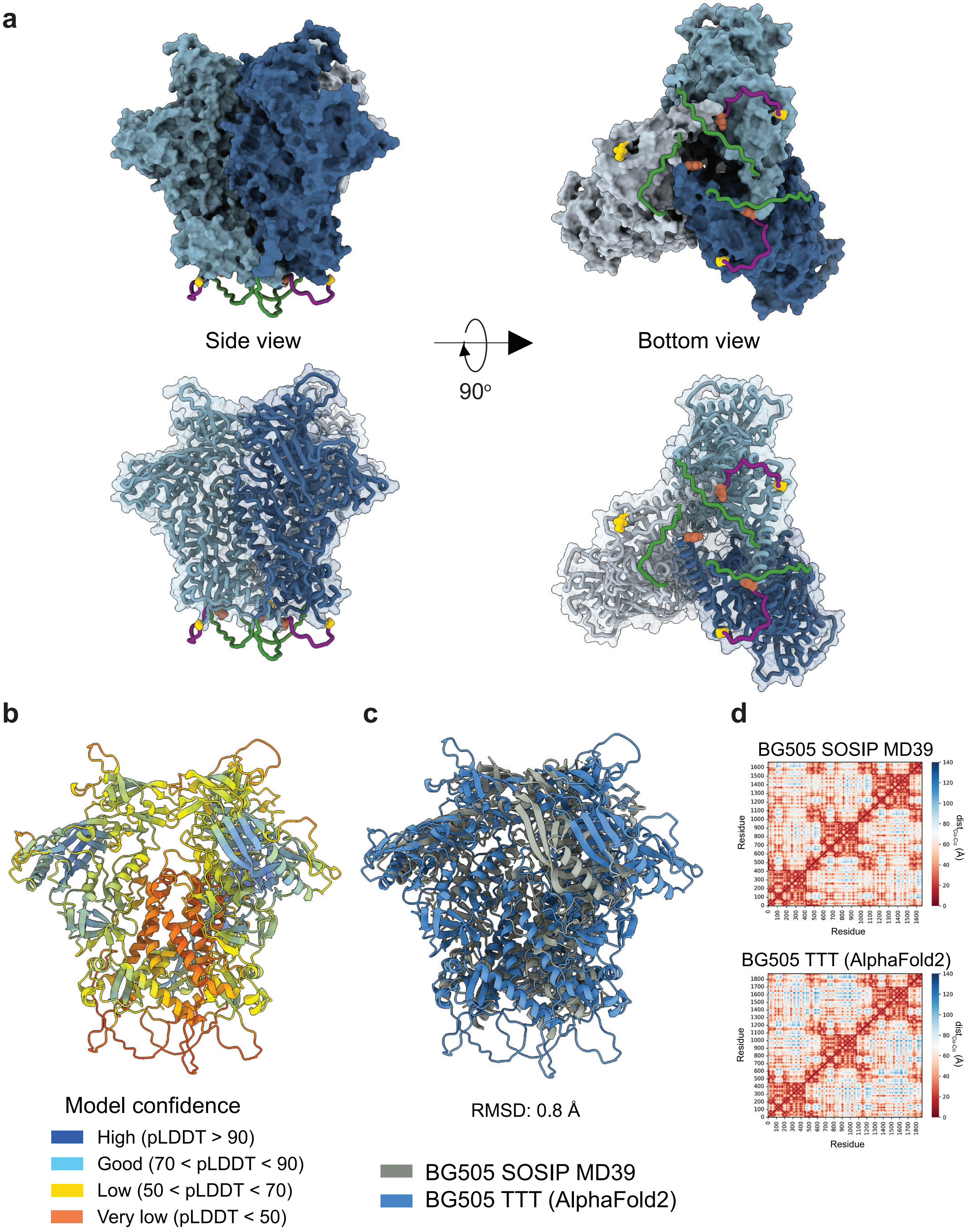


**Fig. S2 AlphaFold2-predicted structure of BG505 TTT. a,b** Predicted BG505 TTT structure, with SC (green) and TTT (purple) linkers at the base of the trimer **(a)** or colored according to the per-residue estimate of the local confidence of the prediction (pLDDT value) **(b)**. **c** Alignment of the predicted BG505 TTT structure to the experimentally determined structure of a native-like BG505 SOSIP MD39 trimer (PDB 7L8D), and the corresponding $C_\alpha$ root-mean-square deviation (RMSD) value. **d** Distance maps of 7L8D and AlphaFold2-predicted BG505 TTT protein structures.

**a**

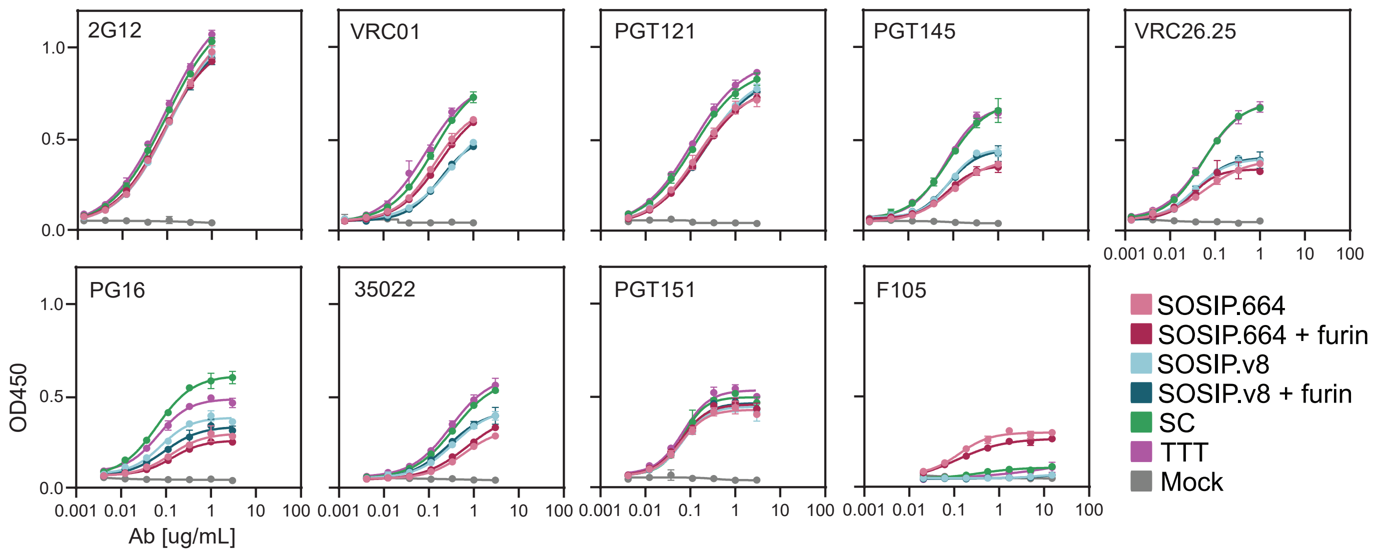


**Fig. S3 StrepTactinXT ELISA with supernatants of HEK293T cells transfected with BG505 SOSIP.664, SOSIP.v8, SC and TTT constructs against a panel of bNAbs (2G12, VRC01, PGT121, PGT145, VRC26.25, PGT151) and non-NAbs (F105).** Dots and error bars represent the average and standard deviation of values obtained in duplicate.

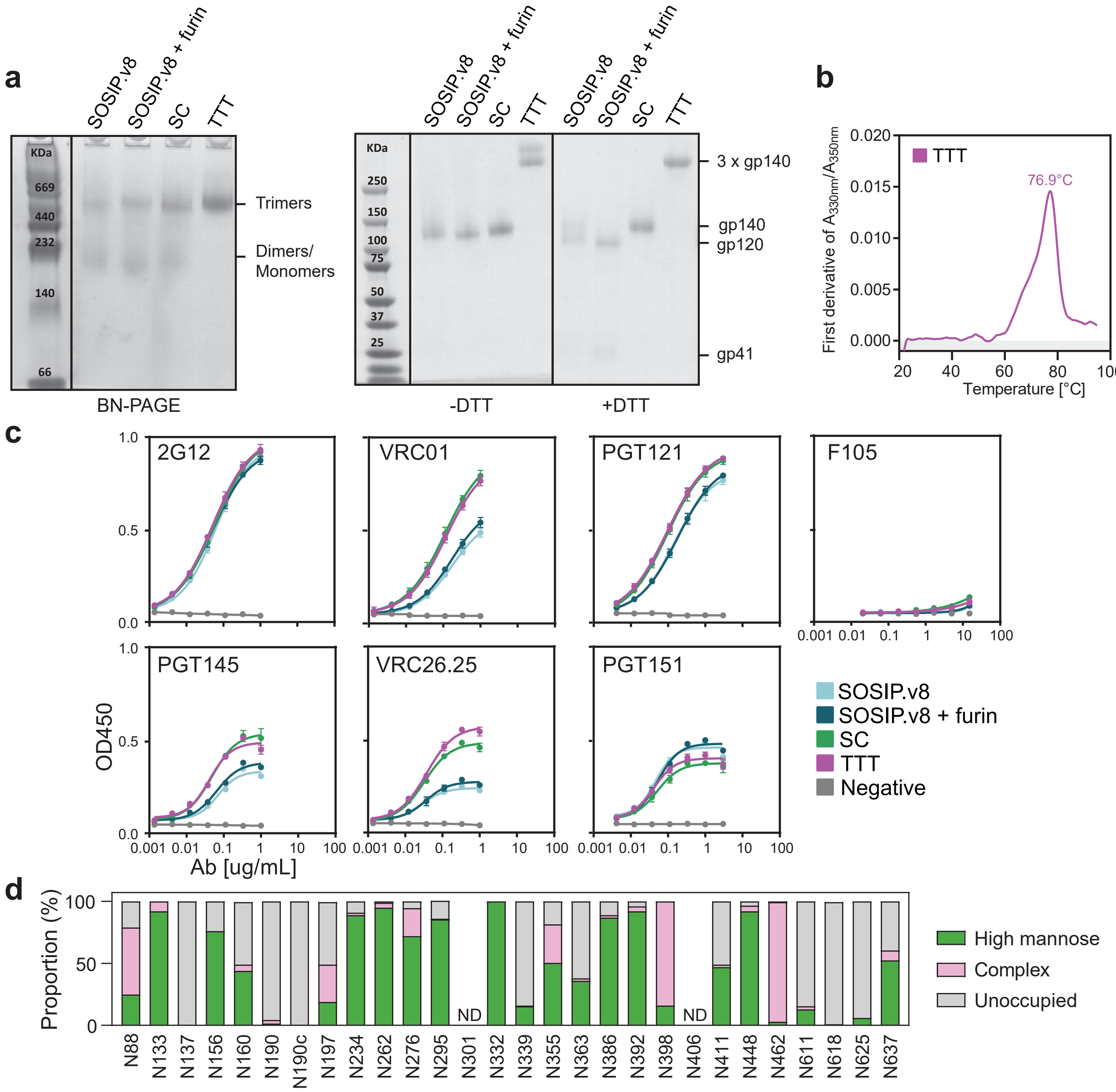


**Fig. S4 Biophysical characterization of BG505 TTT purified protein. a** BN-PAGE, non-reducing (-DTT) and reducing (+DTT) SDS-PAGE analysis of StrepTactinXT-purified SOSIP.v8, SC and TTT protein preparations. The single band above 250 kDa in reducing SDS-PAGE confirmed that BG505 TTT forms covalently linked trimers. **b** Denaturing profile of GNL-purified TTT protein, obtained by nanoDSF and used to determine the $T_m$ value presented on **Table 1. c** StrepTactinXT ELISA with StrepTactinXT-purified SOSIP.v8, SC and TTT protein preparations against a panel of bNAbs (2G12, VRC01, PGT121, PGT145, VRC26.25, PGT151) and non-NAbs (F105). Dots and error bars represent the average and standard deviation of values obtained in duplicate. **d** Site-specific glycan analysis of GNL-purified BG505 TTT protein. Values are specified in **Table S2.** PNGS are displayed as aligned with HxB2. Data could not be determined (ND) for sites N301 and N406. The glycan modifications on the remaining sites were classified into three categories: high mannose (corresponding to any composition containing two HexNAc residues, or three HexNAc and at least 5 hexoses), complex, or unoccupied. The proportion of peptides and glycopeptides corresponding to each of these categories was colored green for high mannose, pink for complex and grey for unoccupied. It is important to note that as the amino acid sequence for all subunits is identical, this data represents an average of the glycosylation across all three protomers that comprise the trimer, and it is not possible to assign particular glycan occupancy and processing states to a particular protomer in the trimer. This analysis revealed that the processing of BG505 TTT was comparable with that of SOSIP trimers[1], and that key glycan bnAb epitopes are conserved across both formats. For instance, the key N332 glycan supersite and the N262 site, which has been shown to **(continues on next page)**

**(continued from previous page)** stabilize the structure of Env, contained almost exclusively oligomannose-type glycans. Furthermore, sites that are occupied by fully processed, complex-type glycans are conserved across both platforms, such as at N398 and N462. Additionally, several sites displayed prominent levels of unoccupied asparagines, most notably on gp41 and in the V1V2 region. These sites are similarly underoccupied on SOSIP trimers[1]. In addition to these sites, the BG505 TTT displayed reduced occupancy at N160, which might be caused by the fact that the TTT was not purified by PGT145, which recognizes oligomannose-type glycans at this site. Additionally, N339, N363 and N411 were more unoccupied on BG505 TTT compared to BG505 SOSIP.

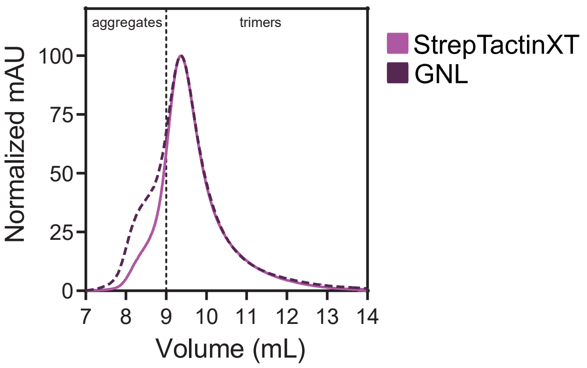


**Fig. S5 SEC profiles of StrepTactinXT- and GNL-purified BG505 TTT protein preparations expressed in HEK293F cells on a Superdex 200 Increase 10/300 GL column.**

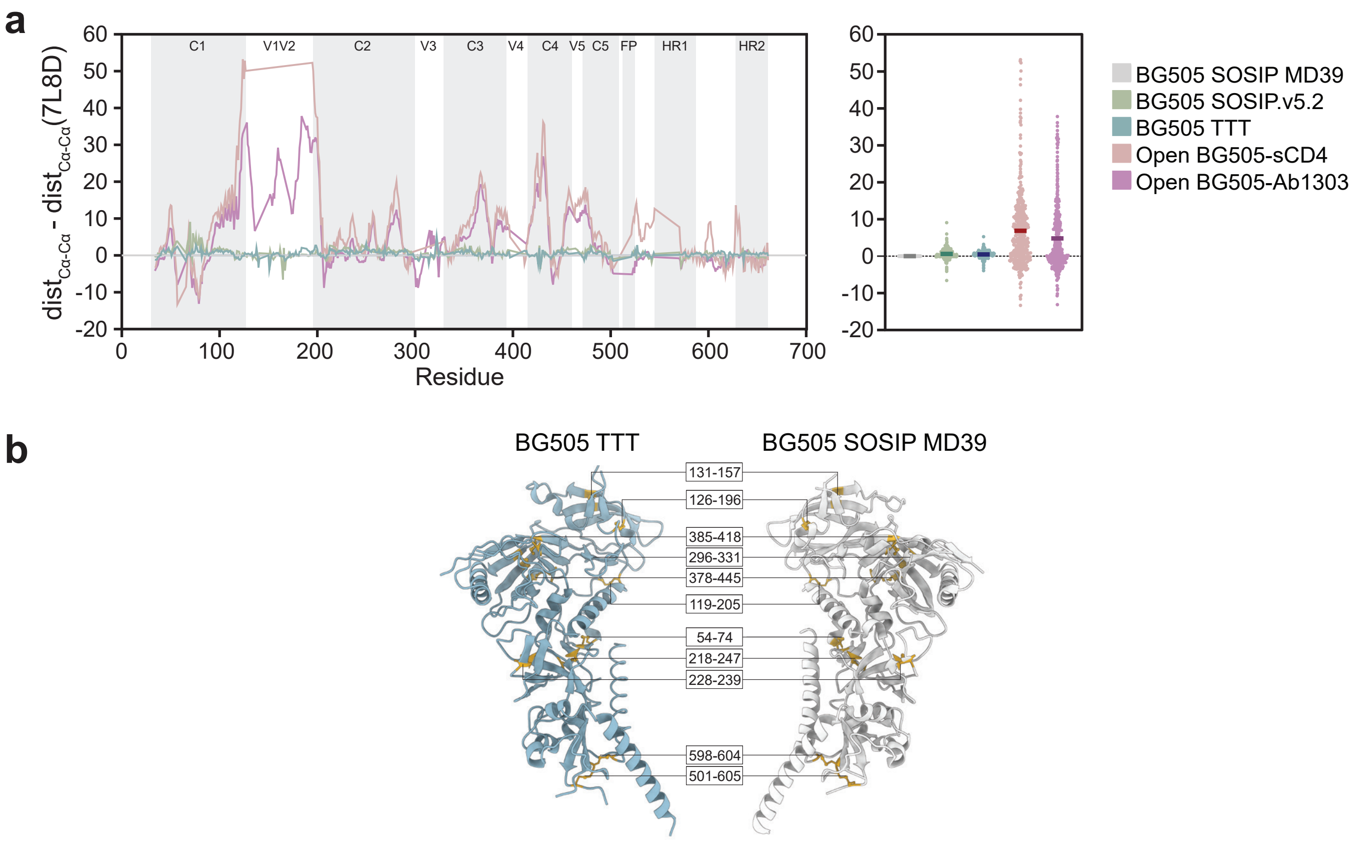


**Fig. S6 X-ray structure of BG505 TTT. a** Interprotomer distances between the $C_\alpha$ atoms of all solved residues in the structures of BG505 TTT, BG505 SOSIP MD39 (PDB 7L8D), BG505 SOSIP v5.2 (PDB 6VO0) and BG505 trimers in an open conformation complexed with sCD4 (PDB 5THR) or the CD4-targeting Ab1303 antibody (PDB 7TFN). **b** Disulfide network of BG505 TTT and BG505 SOSIP MD39 proteins, with Cys residues colored in yellow and numbered according to HxB2.

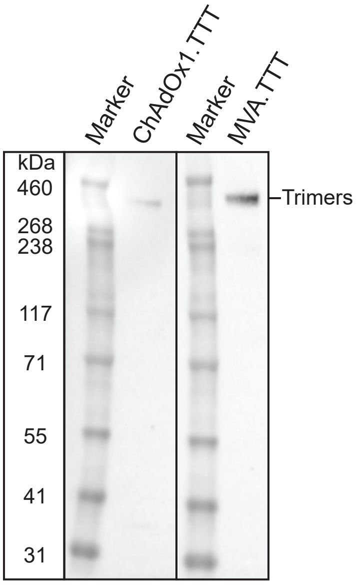


**Fig. S7 Expression of ChAdOx1- and MVA-vectored BG505 TTT in HeLa cells.** TTT expression was confirmed by Western blot analysis of HeLA cell-infected lysate, using an anti-HIV-1 Env monoclonal (ARP3119) primary antibody and a peroxidase-conjugated goat anti-mouse secondary antibody followed by chemiluminescence.

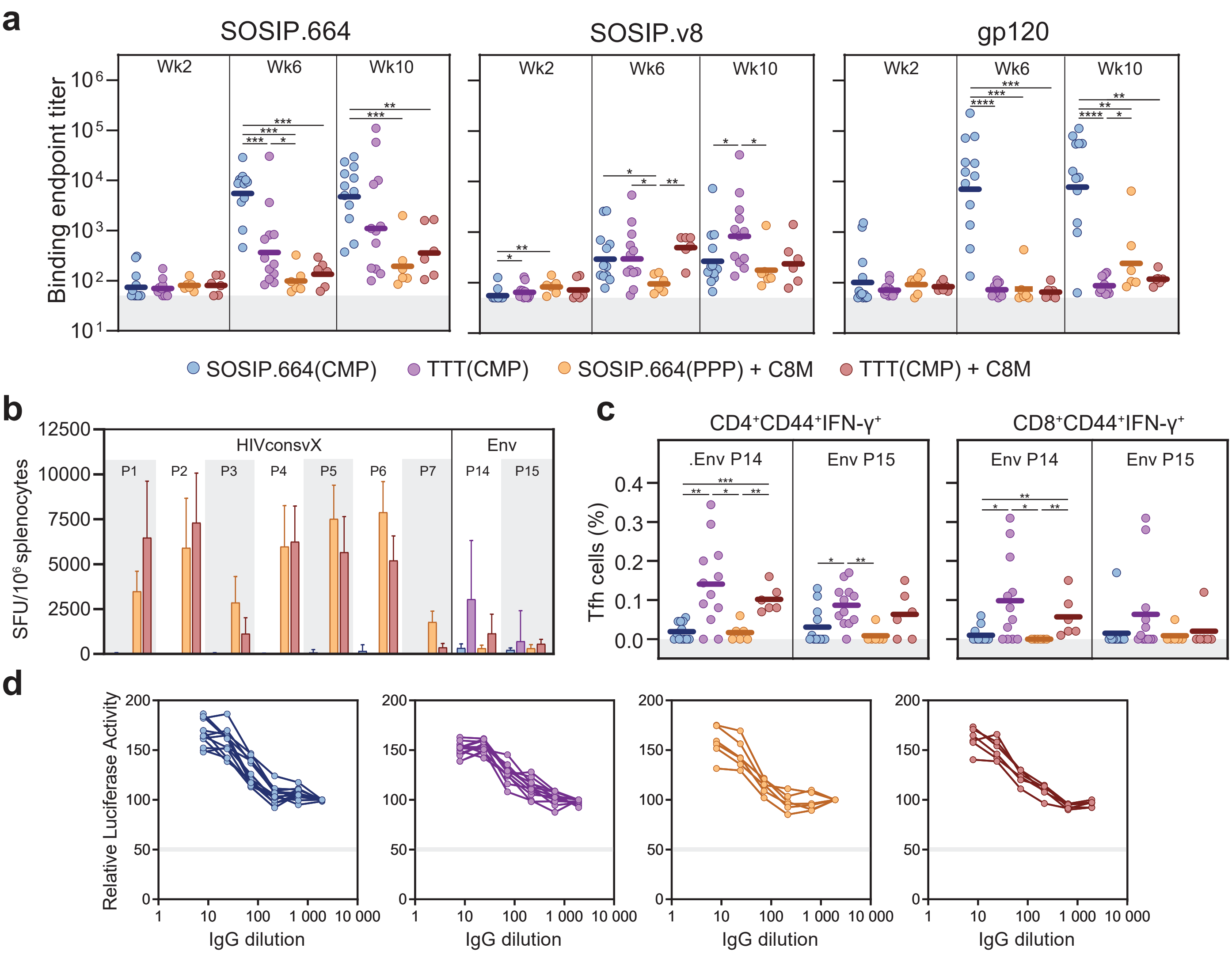


**Fig. S8 Immunogenicity of viral-vectored TTT in mice. a** Endpoint antibody-binding titers of mice sera over time against SOSIP.664, SOSIP.v8 and gp120 proteins, as measured by D7324-capture ELISA. **b** T cell responses against peptide pools covering HIVconsvX (P1-P7) and Env (P14-P15) epitopes, represented as the frequencies of reactive cells, determined using an IFN-γ ELISPOT assay and expressed as spot-producing units (SFU) per $10^6$ splenocytes. **c** Phenotype of vaccine-elicited follicular T cells determined by flow cytometry analysis, indicated by the frequencies of T follicular helper (Tfh) cells ($PD\text{-}1^+CXCR5^+Bcl6^+ICOS^+CCR7^-$) inside the effector-memory ($CD44^+IFN\text{-}\gamma^+$) $CD4^+$ and $CD8^+$ T cell compartments. The Env and T cell immunogen co-immunization, in groups 3 and 4, achieved the goal of inducing strong and broad T cell responses focused predominantly on the vulnerable epitopes of HIV presented by HIVconsvX and minimally reverted to non-protective Env T cells **(b,c)**, while inducing the desired anti-Env Abs **(a)**. Median **(a)** or mean values **(b,c)** are represented by horizontal lines **(a,c)** or bars **(b),** while standard deviations are represented by error bars **(b).** Statistical differences between groups **(a,c)** were determined using a two-tailed Mann-Whitney U test (*p < 0.05, **p < 0.01, ***p < 0.001, ****p < 0.0001). Individual values **(a-c)** are detailed in **Tables S4-S6. d** Neutralization of an autologous BG505/T332N pseudovirus by IgGs purified from week 10 mice sera. Dots represent mean values of replicate samples, determined by a TZM-bl assay.

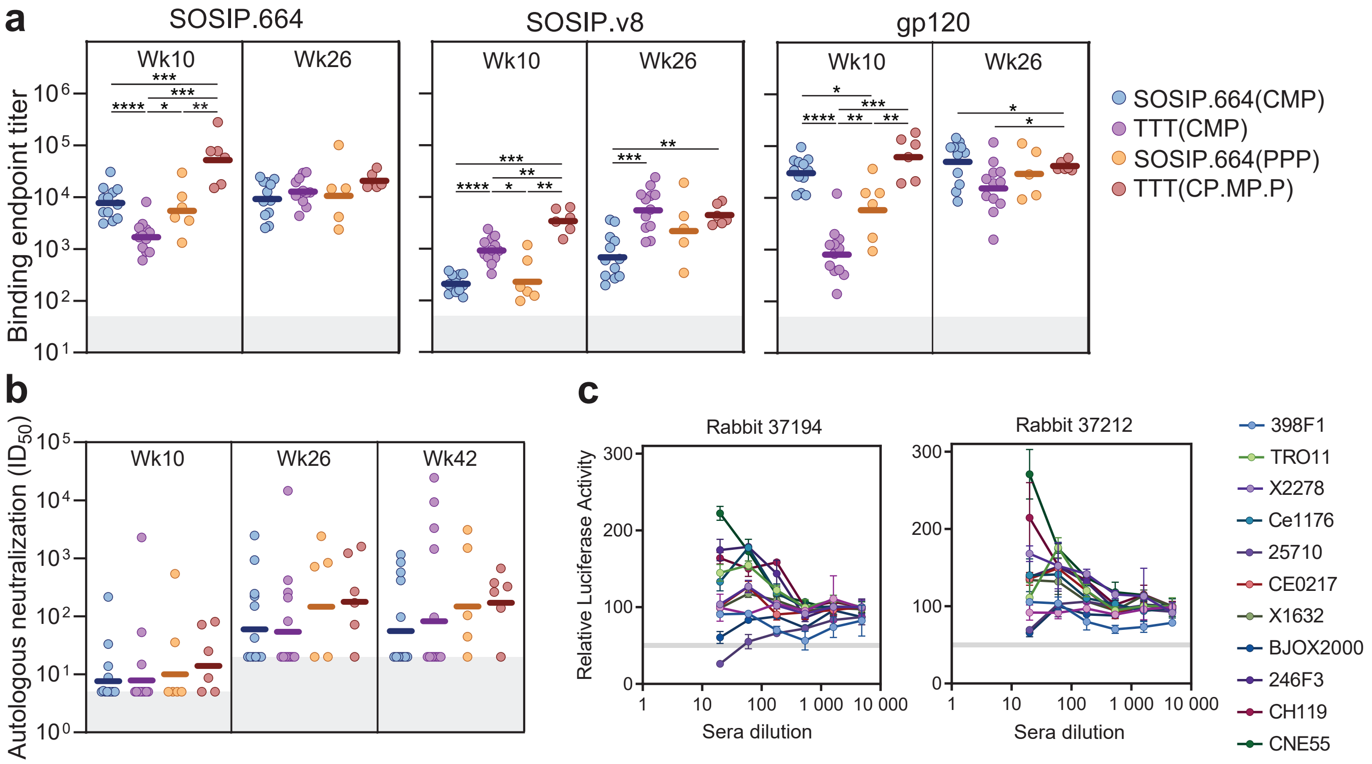


**Fig. S9 Immunogenicity of viral-vectored TTT in rabbits. a** Endpoint antibody binding titers at weeks 10 and 26 against SOSIP.664, SOSIP.v8 and gp120 proteins, as measured by D7324-capture ELISA. **b** Midpoint neutralization titers ($ID_{50}$) at weeks 10, 26 and 42 for sera of the immunized rabbits against a BG505/T332N autologous pseudovirus. **c** Neutralization of a panel of heterologous pseudoviruses by the week 42 sera of rabbits 37194 and 34212. Dots represent mean values of replicate samples determined by a TZM-bl assay. Nine of the viruses used are part of the global panel of HIV-1 Env reference strains[2]. Horizontal lines **(a,b)** represent median **(a)** and geometric mean **(b)** values. Individual values **(a,b)** are detailed in Tables **S7,S8.** Statistical differences between groups **(a,b)**, determined by a two-tailed Mann-Whitney U test, are represented by asterisks (*$p < 0.05$, **$p < 0.01$, ***$p < 0.001$, ****$p < 0.0001$).

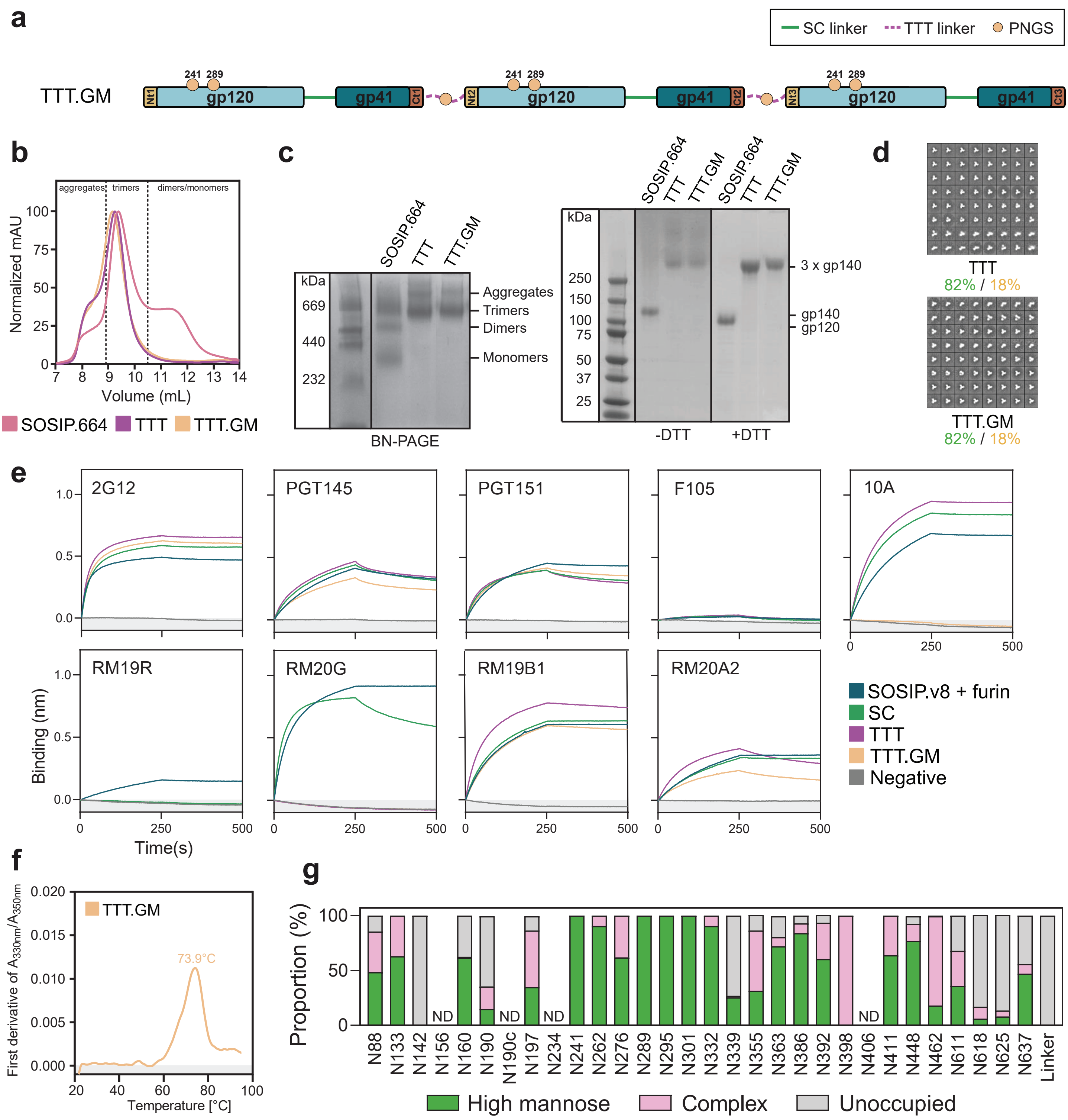


**Fig. S10 Design and biophysical characterization of BG505 TTT.GM construct.** **a** Linear representation of the BG505 TTT.GM construct, derived from BG505 TTT **(Fig. 1a)** by including PNGS motifs at positions 241 and 289 and on the TTT linkers. **b** SEC profile of GNL-purified BG505 SOSIP.664, TTT and TTT.GM protein preparations on a Superdex 200 Increase 10/300 GL column. **c** BN-PAGE, non-reducing (-DTT) and reducing (+DTT) SDS-PAGE analysis of GNL-purified SOSIP.664, TTT and TTT.GM protein preparations. **d** nsEM-generated 2D class averages generated by nsEM analysis of the GNL-purified TTT and TTT.GM proteins. Percentages of native-like trimers (green), malformed trimers (yellow) and monomers/dimers (red) are indicated. **e** ProtA BLI assay with GNL-purified SOSIP, SC, TTT and TTT.GM protein preparations against a panel of bNAbs (2G12, PGT145, PGT151), non-NAbs (F105), 241/289 hole-targeting (10A) and base hole-targeting (RM19R, RM20G, RM19B1 and RM20A2) antibodies. **f** Denaturing profile of GNL-purified TTT.GM protein, obtained by nanoDSF and used to determine the Tm value presented on **Table 1.** **g** Site-specific glycan analysis of GNL-purified TTT.GM protein. Values are specified in **Table S9.** PNGS are displayed as aligned with HxB2. Data could not be determined (ND) for sites N156, N190c, N234 and N406. The glycan **(continues on next page)**

**(continued from previous page)** modifications on the remaining sites were classified into three categories: high mannose (corresponding to any composition containing two HexNAc residues, or three HexNAc and at least 5 hexoses), complex, or unoccupied. The proportion of peptides and glycopeptides corresponding to each of these categories was colored green for high mannose, pink for complex and grey for unoccupied. It is important to note that as the amino acid sequence for all subunits is identical, this data represents an average of the glycosylation across all three protomers that comprise the trimer, and it is not possible to assign particular glycan occupancy and processing states to a particular protomer in the trimer. As expected, the newly incorporated N241 and N289 sites were efficiently populated by oligomannose-type glycans, which explained the 10A binding abrogation. Furthermore, the insertion of these sites appeared to subtly influence the processing of sites around the 241 and 289 positions and increased the abundance of oligomannose-type glycans at N611, as previously reported[3]. The PNGS incorporated in the TTT linkers were, however, largely unoccupied.

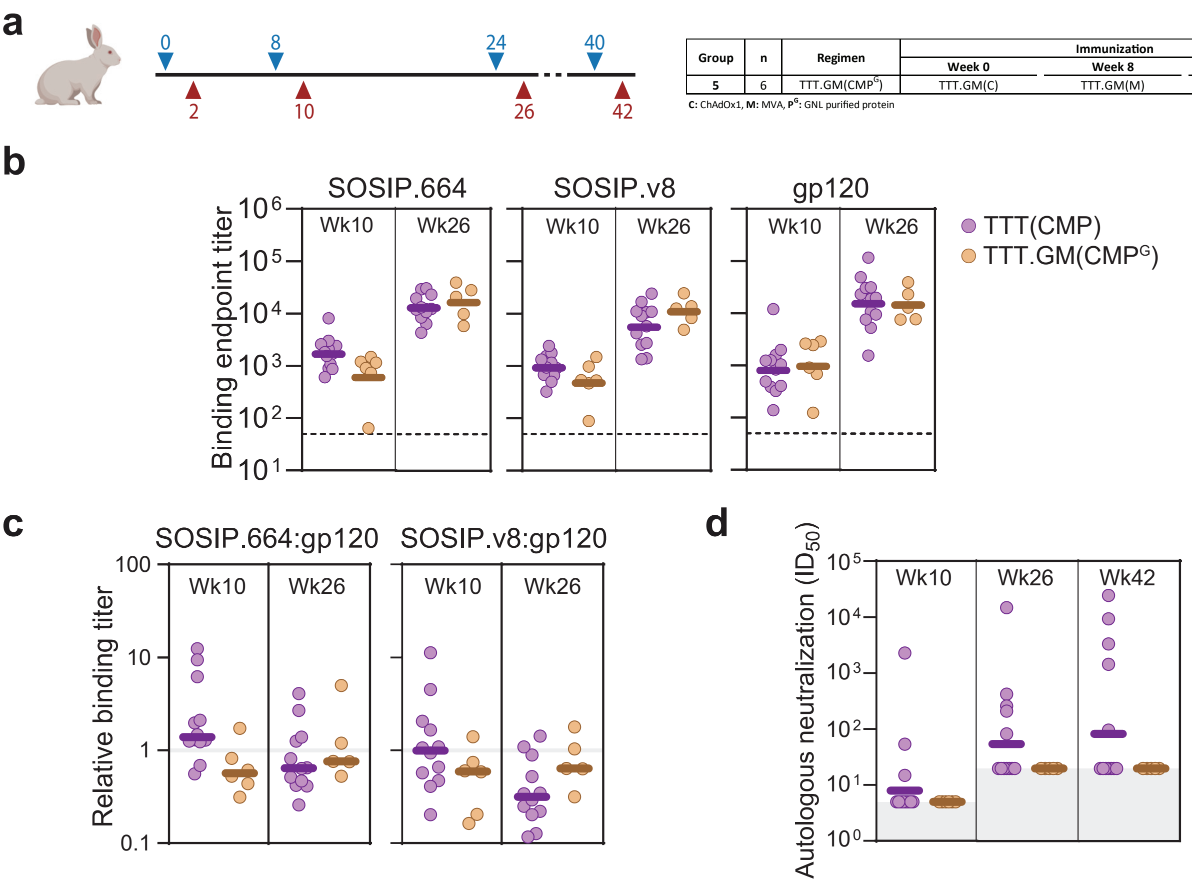


| Group | n | Regimen | Immunization | | |
|---|---|---|---|---|---|
| | | | Week 0 | Week 8 | Week 24 and 40 |
| **5** | 6 | TTT.GM(CMP$^G$) | TTT.GM(C) | TTT.GM(M) | TTT.GM(P$^G$) |

**C:** ChAdOx1, **M:** MVA, **P$^G$:** GNL purified protein

**Fig. S11 Immunogenicity of viral-vectored TTT.GM in rabbits. a** Immunization schedule of rabbits vaccinated with TTT.GM. A group of six NZW rabbits were vaccinated (blue arrows) at weeks 0, 8, 24 and 40 with the immunogens indicated on the table. Antibody responses were evaluated (red arrows) at weeks 10, 26 and 42. **b** Endpoint antibody binding titers at weeks 10 and 26 against SOSIP.664, SOSIP.v8 and gp120 proteins, as measured by D7324-capture ELISA. **c** Ratios between trimer (SOSIP.664 or SOSIP.v8)-binding and gp120-binding titers **d** Midpoint neutralization titers ($ID_{50}$) at weeks 10, 26 and 42 for sera of the immunized rabbits against a BG505/T332N autologous pseudovirus. Horizontal lines represent median **(b,c)** and geometric mean **(d)** values. Individual values **(b,d)** are detailed in **Tables S7,S8.** No statistical differences between groups **(b-d)** were found, as estimated by a two-tailed Mann-Whitney U test.

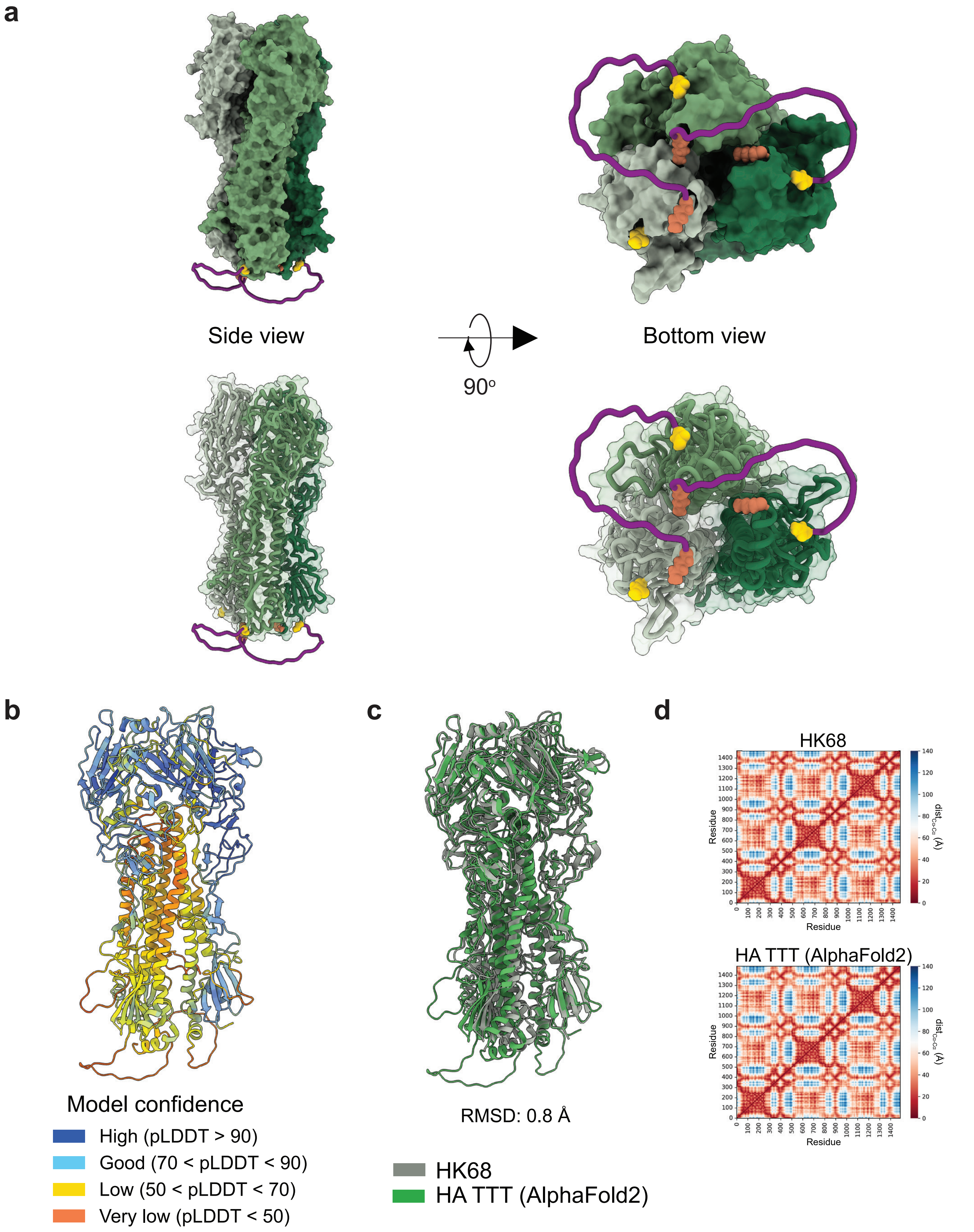


**Fig. S12 AlphaFold2-predicted structure of H3 HA TTT. a,b** Predicted HA TTT structure, with TTT (purple) linkers at the base of the trimer **(a)** or colored according to the per-residue estimate of the local confidence of the prediction (pLDDT value) **(b). c** Alignment of the predicted HA TTT structure to the experimentally determined structure of a native-like HK68 (A/Hong Kong/1/1968(H3N2), PDB 6NHP) HA trimer, and the corresponding $C_\alpha$ root-mean-square deviation (RMSD) value. **d** Distance maps of HK68 and HA TTT predicted structures.

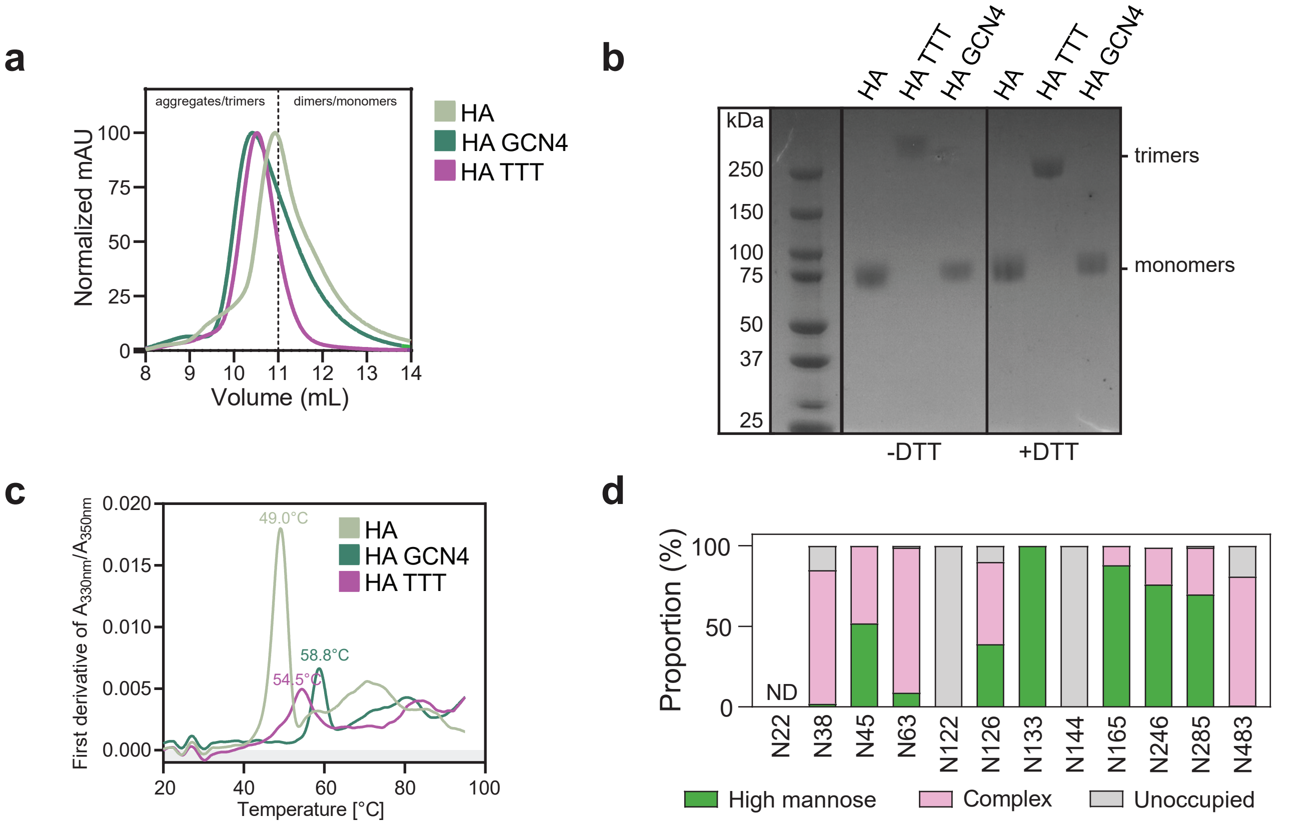


**Fig. S13 Biophysical characterization of the H3 HA TTT protein. a** SEC profiles of StrepTactinXT-purified H3 HA, HA GCN4 and HA TTT protein preparations expressed in HEK293F cells on a Superdex 200 Increase 10/300 GL column. **b** Non-reducing (-DTT) and reducing (+DTT) SDS-PAGE analysis of StrepTactinXT-purified HA, HA GCN4 and HA TTT proteins. The single band above 250 kDa in reducing SDS-PAGE confirmed that HA TTT forms covalently linked trimers. **c** Denaturing profiles of the purified HA, HA GCN4 and HA TTT proteins, obtained by nanoDSF and used to determinate the Tm values presented on **Table 1. d** Site-specific glycan analysis of purified HA TTT. Values are specified in **Table S10.** PNGS are displayed according to H3 numbering. Data could not be determined (ND) for site N22. The glycan modifications on the remaining sites were classified into three categories: high mannose (corresponding to any composition containing two HexNAc residues, or three HexNAc and at least 5 hexoses), complex, or unoccupied. The proportion of peptides and glycopeptides corresponding to each of these categories was colored green for high mannose, pink for complex and grey for unoccupied. It is important to note that as the amino acid sequence for all subunits is identical, this data represents an average of the glycosylation across all three protomers that comprise the trimer, and it is not possible to assign particular glycan occupancy and processing states to a particular protomer in the trimer. This site-specific glycan analysis revealed diverse distributions of oligomannose- and complex-type glycans at different sites. Underprocessed oligomannose glycans were dominant at N133, N165, N246 and N285, while N38, N63 and N483 were occupied mostly with complex glycans. N45 and N126 showed high abundance of both oligomannose and complex glycans. Overall, the glycosylation of the HA TTT corresponds well with that of H3 HA as analyzed previously[4,5]. Both these studies revealed high levels of underprocessed oligomannose-type glycans at N133, N165, N246 and N285, while mixtures or complex glycans were present at other sites. Sites N122 and N144 were not occupied by a glycan on HA TTT, but this is also consistent with previous observations using trimeric HA protein[5], implying that the underoccupancy at these sites is not a result of the TTT format, but rather a consequence of the surrounding primary sequence and/or structure.

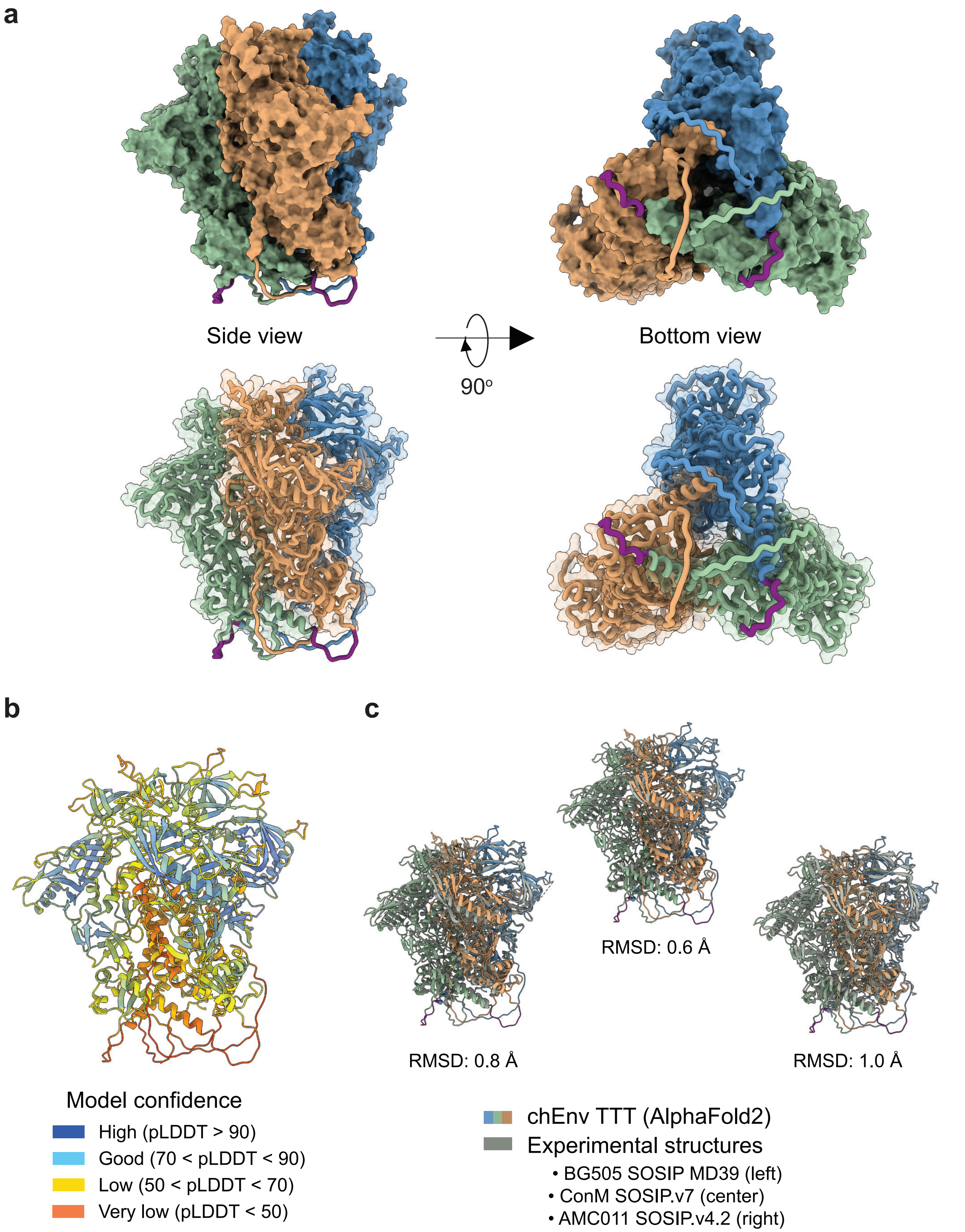


**Fig. S14 AlphaFold2-predicted structure of chEnv TTT.** **a** Predicted chEnv TTT structure, with SC (green) and TTT (purple) linkers at the base of the trimer. BG505, ConM and AMC011 protomers are colored in blue, green and orange, respectively. **b** Predicted structure colored according to the per-residue estimate of the local confidence of the prediction (pLDDT value). **c** Alignment of the predicted chEnv TTT structure to the experimentally determined structures of native-like BG505 SOSIP MD39 (PDB 7L8D), ConM SOSIP.v7 (PDB 6IEQ) and AMC011 SOSIP.v4.2 (PDB 6NC3), and the corresponding $C_\alpha$ root-mean-square deviation (RMSD) values.

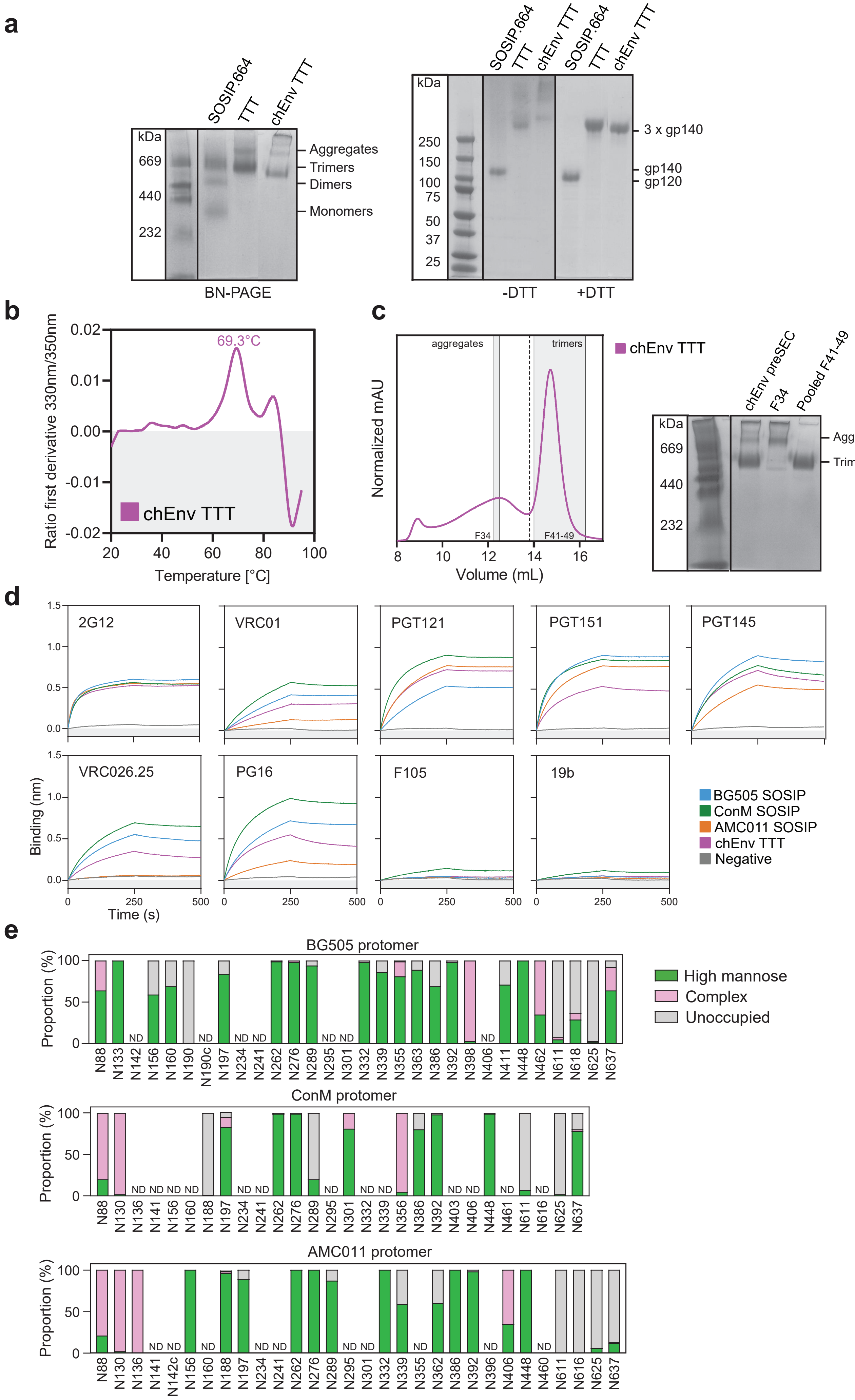
a
SOSIP.664
TTT
chEnv TTT
kDa
669
440
232
Aggregates
Trimers
Dimers
Monomers
BN-PAGE
250
150
100
75
50
37
25
3 x gp140
gp140
gp120
-DTT
+DTT
b
69.3°C
chEnv TTT
Ratio first derivative 330nm/350nm
Temperature [°C]
c
aggregates
trimers
F34
F41-49
Normalized mAU
Volume (mL)
chEnv preSEC
Pooled F41-49
d
2G12
VRC01
PGT121
PGT151
PGT145
VRC026.25
PG16
F105
19b
Binding (nm)
Time (s)
BG505 SOSIP
ConM SOSIP
AMC011 SOSIP
chEnv TTT
Negative
e
BG505 protomer
ConM protomer
AMC011 protomer
Proportion (%)
High mannose
Complex
Unoccupied
ND

**Fig. S15 Biophysical characterization of the chEnv TTT protein. a** BN-PAGE, non-reducing (-DTT) and reducing (+DTT) SDS-PAGE analysis of PGT145-purified chEnv TTT protein. GNL-purified SOSIP.664 and BG505 TTT were included as controls. The single band above 250 kDa in reducing SDS-PAGE confirmed that chEnv TTT forms covalently linked trimers, similarly to BG505 TTT. **b** Denaturing profile of the PGT145-purified chEnv TTT protein, obtained by nanoDSF and used to determine the $T_m$ value presented on Table 1. **c** SEC profile of the PT145-purified chEnv TTT protein expressed in HEK293F cells on a Superose 6 Increase 10/300 GL column (left) and BN-PAGE (right) of the input protein and specific (pools of) fractions. **d** ProtA BLI assay with PGT145+SEC-purified chEnv TTT protein against a panel of bNAbs (2G12, VRC01, PGT121, PGT151, PGT145, VRC026.25, PG16) and non-NAbs (F105, 19b). PGT145-purified BG505 SOSIP.v5.2, ConM SOSIP.v7 and AMC011 SOSIP.v9 were included as homotrimeric controls. The experiment was performed in duplicate, and the curves shown correspond to one of these repetitions. **e** Site-specific glycan analysis of PGT145-purified chEnv TTT protein. Values are specified in **Table S11.** PNGS are displayed as aligned with HxB2. Data could not be determined (ND) for several sites. The glycan modifications on the remaining sites were classified into three categories: high mannose (corresponding to any composition containing two HexNAc residues, or three HexNAc and at least 5 hexoses), complex, or unoccupied. The proportion of peptides and glycopeptides corresponding to each of these categories was colored green for high mannose, pink for complex and grey for unoccupied. This analysis revealed a glycosylation profile similar to the one of BG505 TTT **(Fig. S4d),** with the N332 glycan supersite populated almost exclusively by oligomannose-type glycans and gp41 asparagines prominently unoccupied. Sites N88, N130 and N136 in the N-termini of the second (ConM) and third (AMC011) protomers contained a high proportion of complex-type glycans. Remarkably, this high proportion of complex-type glycans in site N130 has been previously observed in a membrane-bound unmodified AMC011 trimer, but not in a soluble SOSIP counterpart[6].

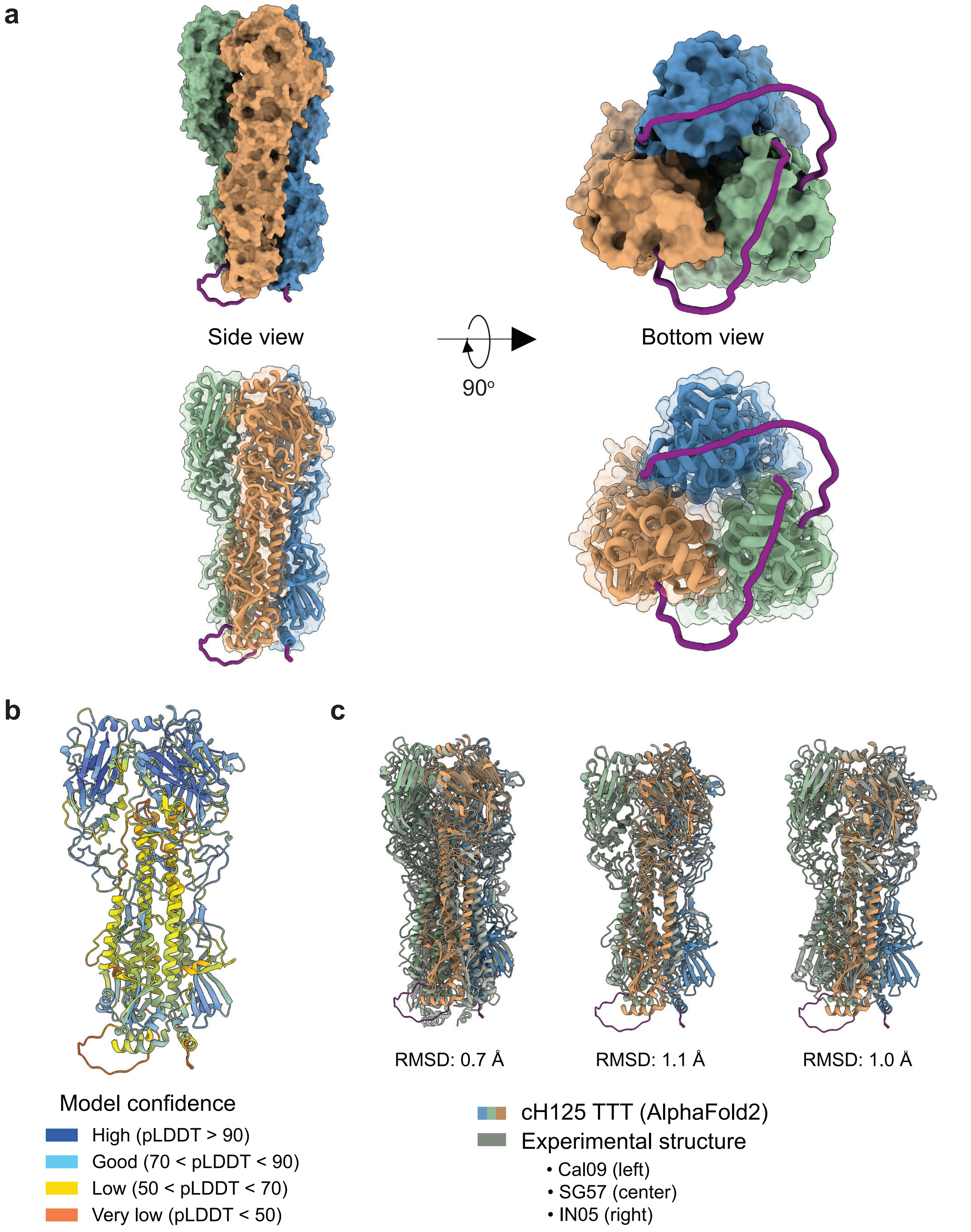


**Fig. S16 AlphaFold2-predicted structure of cH125 TTT. a** Predicted cH125 TTT structure, with TTT (purple) linkers at the base of the trimer and NL09 (A/Netherlands/602/2009(H1N1)), SG57 (A/Singapore/1/1957(H2N2)) and IN05 (A/Indonesia/5/2005(H5N1)) protomers colored in blue, green and orange, respectively. **b** Predicted structure colored according to the per-residue estimate of the local confidence of the prediction (pLDDT value). **c** Alignment of the predicted cH125 TTT structure to the experimentally determined structures of Cal09 (A/California/04/2009(H1N1), PDB 3LZG), SG57 (PDB 2WR7) and IN05 (PDB 4K62) native-like trimers and the corresponding $C_\alpha$ root-mean-square deviation (RMSD) values.

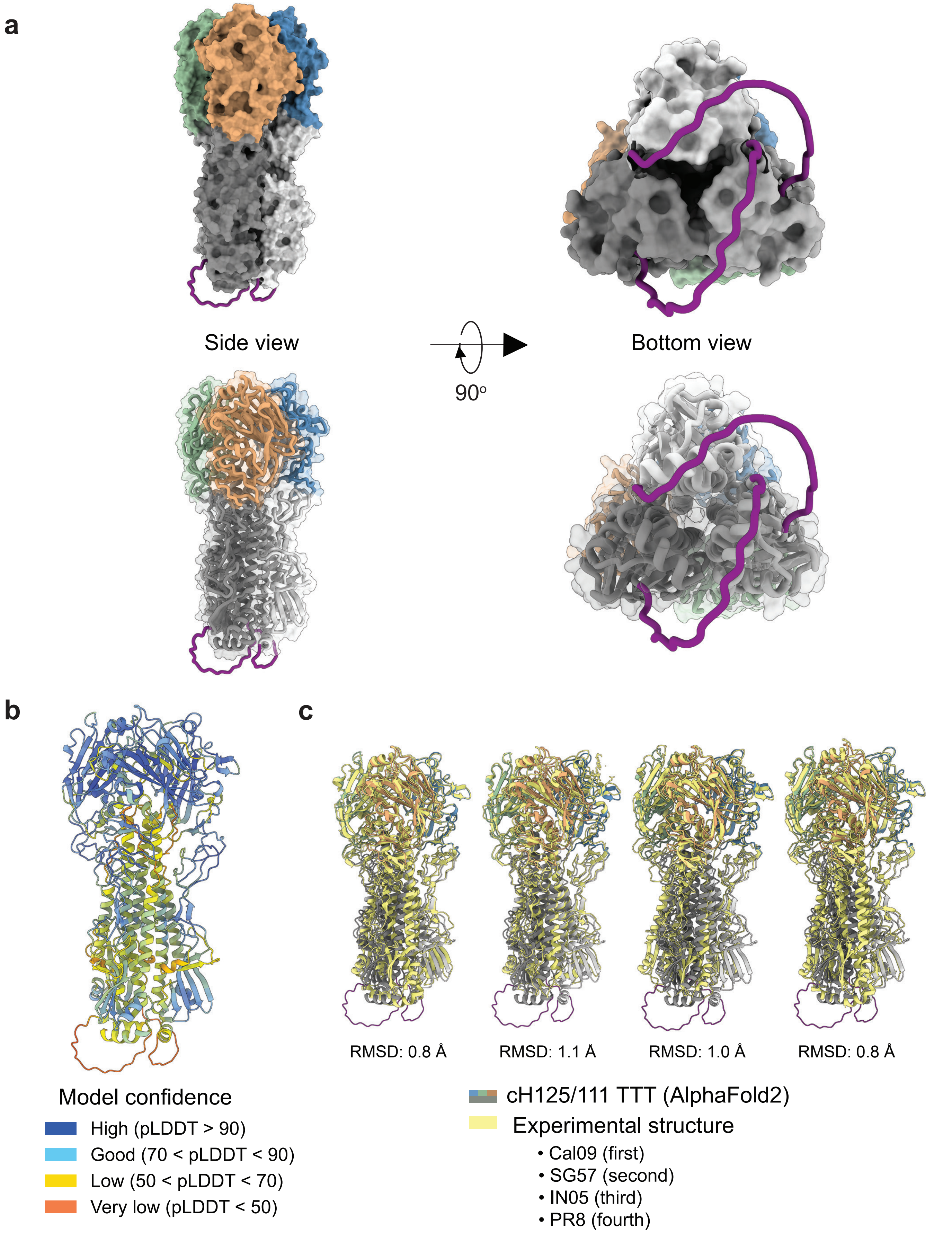


**Fig. S17 AlphaFold2-predicted structure of cH125/111 TTT. a** Predicted cH125/111 TTT structure, with TTT (purple) linkers at the base of the trimer and NL09 (A/Netherlands/602/2009(H1N1)), SG57 (A/Singapore/1/1957(H2N2)), IN05 (A/Indonesia/5/2005(H5N1)) and PR8 (A/Puerto Rico/8/1934/Mount Sinai(H1N1)) regions colored in blue, green, orange and grey, respectively. **b** Predicted structure colored according to the per-residue estimate of the local confidence of the prediction (pLDDT value). **c** Alignment of the predicted cH125/111 TTT structure to the experimentally determined structures of Cal09 (A/California/04/2009(H1N1), PDB 3LZG), SG57 (PDB 2WR7), IN05 (PDB 4K62) and PR8 (PDB 6WCR) native-like trimers and the corresponding $C_\alpha$ root-mean-square deviation (RMSD) values.

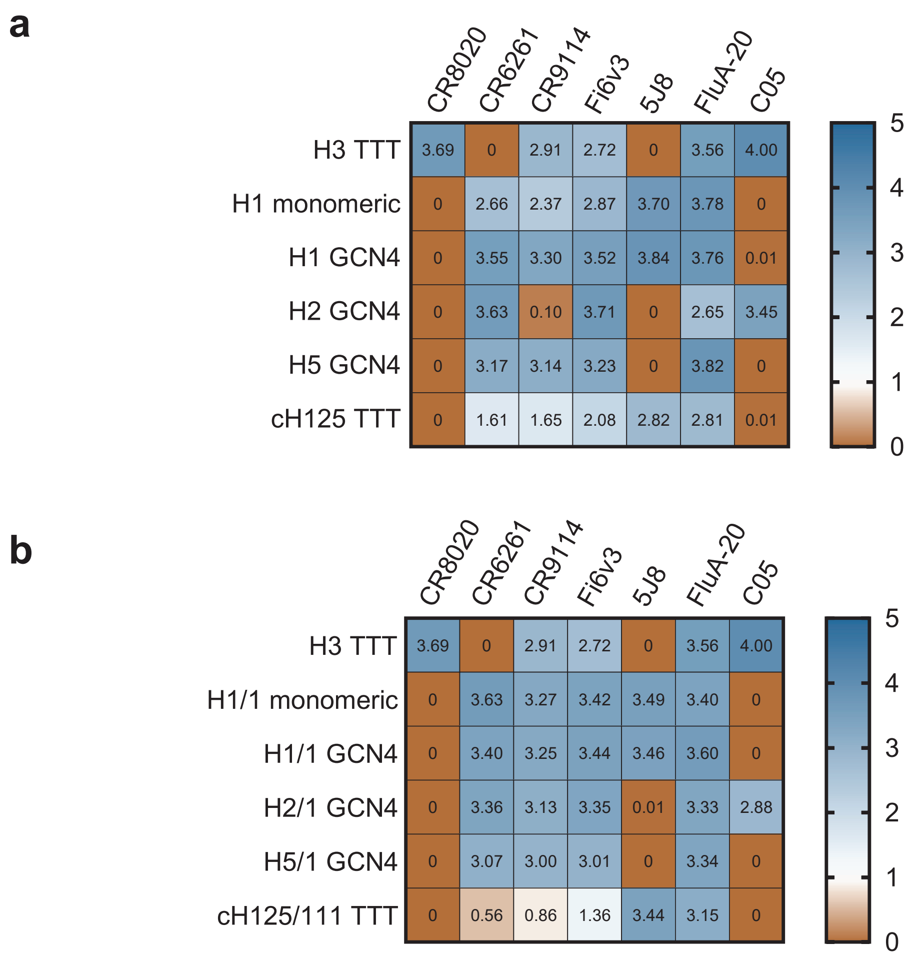


**Fig. S18 StrepTactinXT ELISA with supernatants of HEK293T cells transfected with cH125 TTT and ch125/111 TTT against a panel of stem- (CR8020, CR6261, CR9114, FI6v3) and head-specific (5J8, FluA-20, C05).** Heatmap values represent the area under the curve (AUC) of the ELISA binding curves.

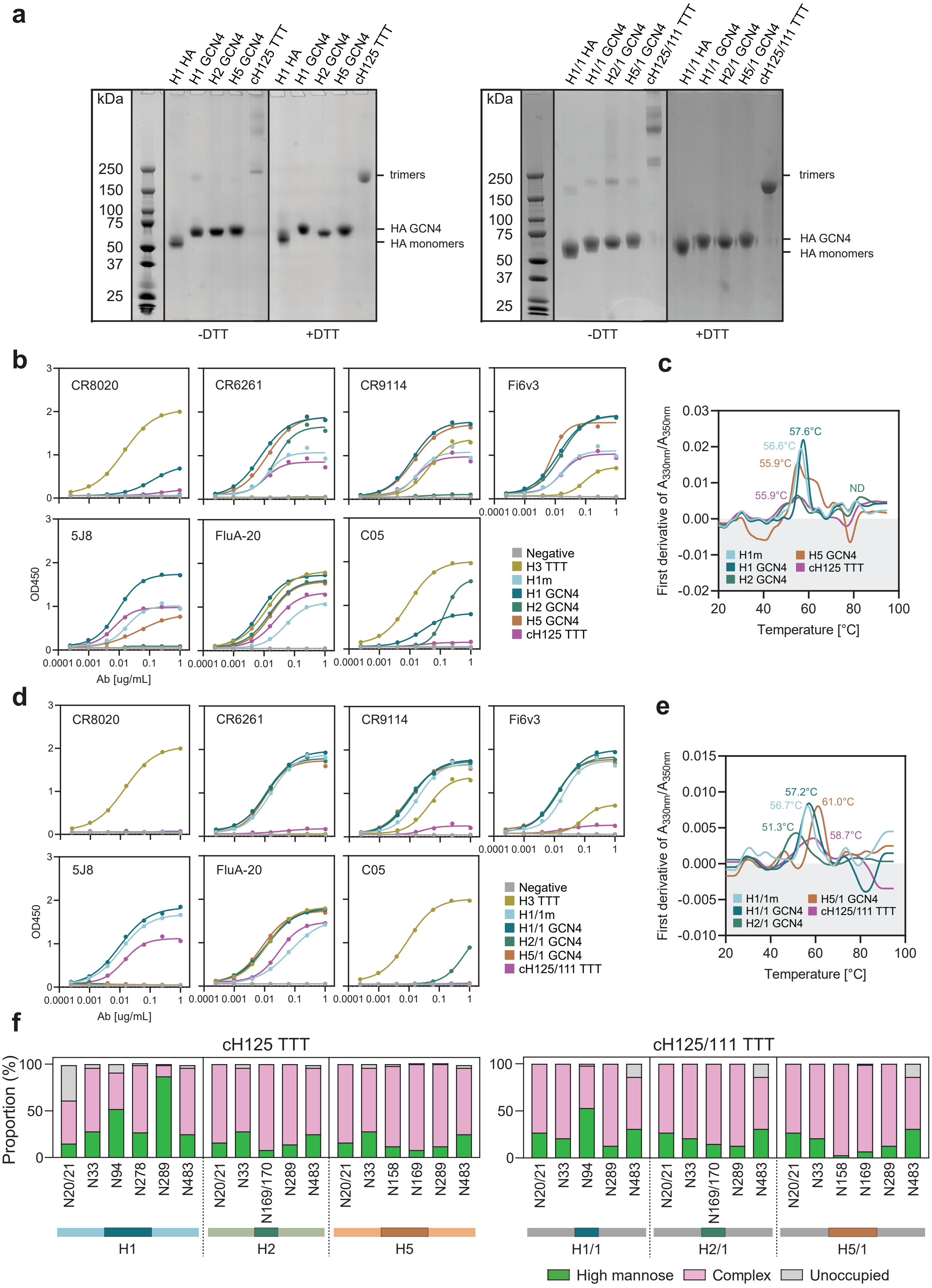
a
H1 HA
H1 GCN4
H2 GCN4
H5 GCN4
cH125 TTT
kDa
250
150
100
75
50
37
25
trimers
HA GCN4
HA monomers
-DTT
+DTT
H1/1 HA
H1/1 GCN4
H2/1 GCN4
H5/1 GCN4
cH125/111 TTT
b
CR8020
CR6261
CR9114
Fi6v3
5J8
FluA-20
C05
OD450
Ab [ug/mL]
Negative
H3 TTT
H1m
H1 GCN4
H2 GCN4
H5 GCN4
cH125 TTT
c
First derivative of A330nm/A350nm
Temperature [°C]
57.6°C
56.6°C
55.9°C
55.9°C
ND
d
H1/1m
H1/1 GCN4
H2/1 GCN4
H5/1 GCN4
cH125/111 TTT
e
57.2°C
56.7°C
61.0°C
51.3°C
58.7°C
f
cH125 TTT
cH125/111 TTT
Proportion (%)
N20/21
N33
N94
N278
N289
N483
N169/170
N158
N169
H1
H2
H5
H1/1
H2/1
H5/1
High mannose
Complex
Unoccupied

**Fig. S19 Biophysical characterization of the chimeric HA TTT proteins. a** BN-PAGE, non-reducing (-DTT) and reducing (+DTT) SDS-PAGE analysis of StrepTactinXT-purified cH125 TTT and cH125/111 TTT proteins and the corresponding monomeric and GCN4-trimerized control proteins. The single band around 250 kDa in reducing SDS-PAGE confirmed that both cH125 TTT and cH125/111 TTT form covalently linked trimers. **b,d** StrepTactinXT ELISA assay with cH125 TTT **(b)** and cH125/111 TTT **(d)** against a panel of stem- (CR8020, CR6261, CR9114, Fi6v3) and head-specific (5J8, FluA-20, C05). **c,e** Denaturing profile of StrepTactinXT-purified cH125 TTT **(c)** and cH125/111 TTT **(e)** proteins, obtained by nanoDSF and used to determine the Tm values presented on **Table 1. f** Site-specific glycan analysis of StreptactinXT-purified cH125 TTT and cH125/111 TTT proteins. Values are specified in **Table S12.** PNGS are displayed according to H3 numbering. The glycan modifications were classified into three categories: high mannose (corresponding to any composition containing two HexNAc residues, or three HexNAc and at least 5 hexoses), complex, or unoccupied. The proportion of peptides and glycopeptides corresponding to each of these categories was colored green for high mannose, pink for complex and grey for unoccupied. The analysis revealed that most sites were effectively populated by glycans, and only the N20/21 site on the first protomer of cH125 TTT and the N483 sites of all three protomers of cH125/111 TTT presented some unoccupancy. The majority of sites were dominated by complex glycans, probably due to the lower density of glycosylation sites compared to the H3 HA TTT protein **(Fig. S13d).**

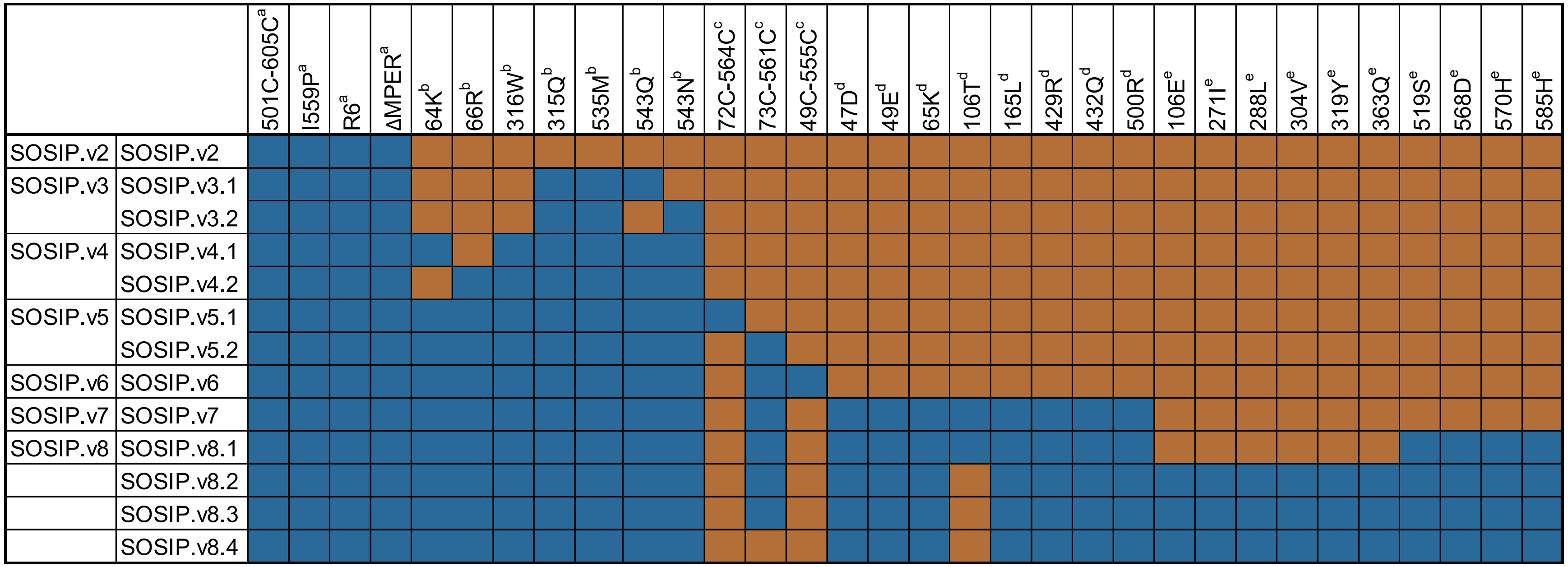


**Table S1 Modifications included in the different SOSIP versions, up to SOSIP.v8.4.** Mutations are described in [a]Sanders et al. 2013[7], [b]Dey et al. 2008[8] and de Taeye et al. 2015[9], [c]de la Peña et al. 2017[10], [d]Guenaga et al. 2015[11], [e]Steichen et al. 2016[12]. The variant 64K (v4.1) is used for BG505 and B41; while the variant 66R (v4.2) is used for the rest of strains. 72C-564C (v5.1) is an alternative disulfide bond that works as well as 73C-561C (v5.2), but we generally use the latter. All the mutations from Guenaga et al. 2015 are naturally present in BG505 wild-type. The 363Q mutation is not used for BG505 SOSIP.v8.4 trimers, as it introduces a glycan hole.

| BG505 TTT | N88 | N133 | N137 | N156 | N160 | N190 | N190c | N197 | N234 | N262 | N276 | N295 | N301 | N332 | N339 | N355 | N363 | N386 | N392 | N398 | N406 | N411 | N448 | N462 | N611 | N618 | N625 | N637 |
|---|---|---|---|---|---|---|---|---|---|---|---|---|---|---|---|---|---|---|---|---|---|---|---|---|---|---|---|---|
| M9Glc | 0 | 0 | 0 | 0 | 0 | 0 | 0 | 0 | 0 | 9 | 0 | 0 | | 0 | 0 | 0 | 0 | 0 | 0 | 0 | | 0 | 0 | 0 | 0 | 0 | 0 | 0 |
| M9 | 0 | 11 | 0 | 63 | 11 | 0 | 0 | 13 | 65 | 66 | 0 | 63 | | 83 | 7 | 0 | 24 | 60 | 9 | 0 | | 0 | 32 | 0 | 0 | 0 | 0 | 0 |
| M8 | 0 | 41 | 0 | 0 | 13 | 0 | 0 | 2 | 14 | 7 | 7 | 13 | | 12 | 6 | 1 | 5 | 10 | 59 | 0 | | 0 | 30 | 0 | 1 | 0 | 0 | 8 |
| M7 | 0 | 11 | 0 | 0 | 6 | 0 | 0 | 1 | 3 | 4 | 13 | 0 | | 0 | 1 | 2 | 2 | 6 | 9 | 0 | | 7 | 10 | 0 | 1 | 0 | 0 | 21 |
| M6 | 3 | 7 | 0 | 0 | 1 | 0 | 0 | 0 | 1 | 2 | 9 | 2 | | 1 | 0 | 2 | 1 | 3 | 3 | 0 | | 8 | 4 | 0 | 1 | 0 | 1 | 4 |
| M5 | 15 | 12 | 0 | 13 | 8 | 1 | 0 | 2 | 5 | 6 | 33 | 3 | | 3 | 1 | 34 | 2 | 7 | 9 | 11 | | 30 | 10 | 1 | 6 | 1 | 2 | 14 |
| M4 | 2 | 6 | 0 | 0 | 3 | 0 | 0 | 0 | 0 | 0 | 3 | 4 | | 0 | 0 | 1 | 0 | 1 | 2 | 0 | | 1 | 3 | 0 | 0 | 0 | 0 | 0 |
| M3 | 0 | 4 | 0 | 0 | 2 | 0 | 0 | 0 | 0 | 0 | 3 | 0 | | 0 | 0 | 1 | 0 | 0 | 1 | 0 | | 0 | 1 | 0 | 0 | 0 | 0 | 0 |
| FM | 0 | 0 | 0 | 0 | 0 | 0 | 0 | 0 | 0 | 0 | 0 | 0 | | 0 | 0 | 0 | 0 | 0 | 0 | 0 | | 0 | 0 | 0 | 0 | 0 | 0 | 0 |
| Hybrid | 4 | 0 | 0 | 0 | 0 | 0 | 0 | 0 | 0 | 0 | 3 | 0 | n.d. | 0 | 0 | 5 | 0 | 1 | 0 | 0 | n.d. | 1 | 2 | 0 | 2 | 0 | 3 | 4 |
| Fhybrid | 0 | 0 | 0 | 0 | 0 | 0 | 0 | 0 | 0 | 0 | 0 | 0 | | 0 | 0 | 5 | 0 | 0 | 0 | 5 | | 0 | 0 | 0 | 0 | 0 | 0 | 1 |
| HexNAc(3)(x) | 6 | 0 | 0 | 0 | 0 | 0 | 0 | 0 | 0 | 0 | 12 | 0 | | 0 | 0 | 1 | 0 | 0 | 0 | 0 | | 0 | 0 | 0 | 0 | 0 | 0 | 1 |
| HexNAc(3)(F)(x) | 0 | 0 | 0 | 0 | 1 | 0 | 0 | 0 | 0 | 1 | 2 | 0 | | 0 | 0 | 9 | 0 | 0 | 0 | 0 | | 1 | 0 | 7 | 0 | 0 | 0 | 0 |
| HexNAc(4)(x) | 14 | 0 | 0 | 0 | 0 | 0 | 0 | 6 | 0 | 0 | 1 | 0 | | 0 | 0 | 0 | 0 | 0 | 0 | 0 | | 0 | 0 | 0 | 0 | 0 | 0 | 0 |
| HexNAc(4)(F)(x) | 11 | 7 | 0 | 0 | 3 | 1 | 0 | 1 | 2 | 2 | 4 | 0 | | 0 | 0 | 12 | 1 | 1 | 1 | 41 | | 1 | 3 | 68 | 1 | 0 | 0 | 6 |
| HexNAc(5)(x) | 12 | 0 | 0 | 0 | 0 | 0 | 0 | 16 | 0 | 0 | 0 | 0 | | 0 | 0 | 0 | 0 | 0 | 0 | 0 | | 0 | 0 | 0 | 0 | 0 | 0 | 0 |
| HexNAc(5)(F)(x) | 10 | 1 | 0 | 0 | 1 | 1 | 0 | 1 | 0 | 0 | 2 | 0 | | 0 | 0 | 8 | 0 | 0 | 0 | 43 | | 0 | 0 | 19 | 2 | 0 | 0 | 1 |
| HexNAc(6+)(x) | 0 | 0 | 0 | 0 | 0 | 0 | 0 | 4 | 0 | 0 | 0 | 0 | | 0 | 0 | 0 | 0 | 0 | 0 | 0 | | 0 | 0 | 0 | 0 | 0 | 0 | 0 |
| HexNAc(6+)(F)(x) | 1 | 0 | 0 | 0 | 0 | 0 | 0 | 1 | 0 | 0 | 0 | 0 | | 0 | 0 | 1 | 0 | 0 | 0 | 0 | | 0 | 0 | 1 | 0 | 0 | 0 | 0 |
| Unoccupied | 21 | 0 | 100 | 24 | 50 | 95 | 100 | 51 | 9 | 1 | 5 | 14 | | 0 | 84 | 18 | 62 | 11 | 4 | 0 | | 51 | 3 | 0 | 84 | 99 | 94 | 39 |
| Core | 0 | 0 | 0 | 0 | 0 | 0 | 0 | 0 | 0 | 1 | 2 | 0 | | 0 | 0 | 1 | 0 | 1 | 2 | 0 | | 1 | 0 | 2 | 0 | 0 | 0 | 0 |

**Table S2 Site-specific glycan analysis of the GNL-purified BG505 TTT protein preparation.** Values indicate the percentage of each type of glycan (rows) at each glycosylation site (columns) and are represented in **Fig. S4d.**

| Data collection | BG505 TTT + PGT124 Fab + 35O22 scFv |
|---|---|
| Beamline | SSRL12-1 |
| Wavelength (Å) | 0.9795 |
| Space group | C2 |
| Unit cell parameters | |
| a, b, c (Å) | 357.6, 245.0, 207.7 |
| α, β, γ (°) | 90, 125.1, 90 |
| Resolution (Å) | 50.0-5.80 (5.90-5.80)[a] |
| Unique reflections | 34,433 (3,202)[a] |
| Redundancy | 6.8 (5.7)[a] |
| Completeness (%) | 97.8 (94.0)[a] |
| $<I/\sigma_I>$ | 12.5 (1.0)[a] |
| $R_{sym}$[b] (%) | 22.1 (>100)[a] |
| $R_{pim}$[b] (%) | 9.1 (79.4)[a] |
| $CC_{1/2}$[c] (%) | 99.2 (66.5)[a] |
| **Refinement statistics** | |
| Resolution (Å) | 40.9-5.80 |
| Reflections (work) | 34,133 |
| Reflections (test) | 3,169 |
| $R_{cryst}$[d] / $R_{free}$[e] (%) | 25.5/29.2 |
| No. of atoms | 29,653 |
| Average *B*-values (Å$^2$) | 359 |
| Wilson *B*-value (Å$^2$) | 224 |
| **RMSD from ideal geometry** | |
| Bond length (Å) | 0.003 |
| Bond angle (°) | 0.57 |
| **Ramachandran statistics (%)[f]** | |
| Favored | 94.6 |
| Outliers | 0.2 |
| **PDB code** | 8TGO |

[a] Numbers in parentheses refer to the highest-resolution shell.
[b] $R_{sym} = \Sigma_{hkl} \Sigma_i | I_{hkl,i} - <I_{hkl}> | / \Sigma_{hkl} \Sigma_i I_{hkl,i}$ and $R_{pim} = \Sigma_{hkl} (1/(n-1))^{1/2} \Sigma_i | I_{hkl,i} - <I_{hkl}> | / \Sigma_{hkl} \Sigma_i I_{hkl,i}$, where $I_{hkl,i}$ is the scaled intensity of the i$^{th}$ measurement of reflection h, k, l, $<I_{hkl}>$ is the average intensity for that reflection, and *n* is the redundancy.
[c] $CC_{1/2}$ = Pearson correlation coefficient between two random half datasets.
[d] $R_{cryst} = \Sigma_{hkl} | F_o - F_c | / \Sigma_{hkl} | F_o | \times 100$, where $F_o$ and $F_c$ are the observed and calculated structure factors, respectively.
[e] $R_{free}$ was calculated as for $R_{cryst}$, but on a test set comprising 10% of the data excluded from refinement.
[f] From MolProbity.

**Table S3 X-ray data collection and refinement statistics.**

| | | | Wk2 | | | Wk6 | | | Wk10 | | |
|---|---|---|---|---|---|---|---|---|---|---|---|
| Group | Regimen | Mouse | SOSIP.664 | SOSIP | gp120 | SOSIP.664 | SOSIP | gp120 | SOSIP.664 | SOSIP | gp120 |
| 1 | SOSIP.664 (CMP) | M1 | <50 | <50 | 68 | 10489 | 137 | 72932 | 23194 | 188 | 113316 |
| | | M2 | <50 | <50 | <50 | 11930 | 579 | 23067 | 4959 | 124 | 6651 |
| | | M3 | <50 | <50 | <50 | 3754 | 77 | 3501 | 3218 | 105 | 11948 |
| | | M4 | 81 | <50 | <50 | 28634 | 2552 | 1379 | 29485 | 7208 | 54414 |
| | | M5 | <50 | <50 | 57 | 9736 | 207 | 12997 | 6034 | 150 | 15939 |
| | | M6 | <50 | <50 | 58 | 4457 | 141 | 15462 | 1741 | 161 | 968 |
| | | M7 | 308 | 50 | 1251 | 8613 | 2617 | 24206 | 18429 | 881 | 55631 |
| | | M8 | <50 | <50 | 52 | 1020 | 480 | 463 | 537 | 207 | 1503 |
| | | M9 | <50 | 126 | 253 | 8592 | 134 | 77461 | 11330 | 482 | 56062 |
| | | M10 | 285 | <50 | 60 | 460 | 114 | 135 | 13412 | 217 | 80076 |
| | | M11 | 124 | <50 | 1525 | 9400 | 218 | 221371 | 367 | 66 | 63 |
| | | M12 | 65 | 75 | 125 | 10297 | 694 | 9165 | 7781 | 886 | 11475 |
| 2 | TTT (CMP) | M13 | 50 | 71 | 74 | 3639 | 547 | 75 | 171 | 329 | 65 |
| | | M14 | 64 | 130 | 55 | 808 | 1585 | 69 | 1242 | 249 | 74 |
| | | M15 | 64 | 57 | 137 | 90 | 177 | 66 | 100 | 304 | 65 |
| | | M16 | 61 | 68 | 77 | 83 | 56 | <50 | 975 | 1120 | 61 |
| | | M17 | 70 | <50 | 56 | 139 | 354 | 55 | 567 | 830 | 136 |
| | | M18 | 72 | <50 | 61 | 169 | 166 | 63 | 137 | 136 | 92 |
| | | M19 | <50 | 55 | 65 | 836 | 426 | 72 | 109639 | 34122 | 160 |
| | | M20 | <50 | 54 | 113 | 283 | 163 | 103 | 56939 | 6092 | 147 |
| | | M21 | 172 | 54 | 58 | 107 | 73 | 86 | 178 | 193 | 70 |
| | | M22 | 70 | 65 | 79 | 678 | 157 | 55 | 9950 | 1015 | 59 |
| | | M23 | 82 | 51 | 67 | 212 | 910 | 112 | 1088 | 1839 | 135 |
| | | M24 | 110 | 114 | 55 | 30145 | 5355 | 98 | 8313 | 2750 | 73 |
| 3 | SOSIP.664 (PPP) C8M | M25 | 83 | 84 | 155 | 60 | 65 | <50 | 113 | 119 | 105 |
| | | M26 | 61 | 89 | <50 | 72 | 148 | <50 | 168 | 124 | 84 |
| | | M27 | 82 | 54 | 112 | 70 | 78 | 54 | 84 | 88 | 102 |
| | | M28 | 126 | 94 | 104 | 123 | 60 | 72 | 249 | 164 | 532 |
| | | M29 | 71 | 63 | 125 | 96 | 110 | 56 | 114 | 150 | 175 |
| | | M30 | 74 | 140 | 58 | 326 | 159 | 446 | 2003 | 1341 | 6445 |
| 4 | TTT (CMP) C8M | M31 | 52 | 138 | 67 | 152 | 775 | 112 | 1581 | 294 | 204 |
| | | M32 | <50 | 57 | 85 | 76 | 783 | 55 | 267 | 196 | 116 |
| | | M33 | 129 | <50 | 98 | 61 | 458 | 51 | 129 | 78 | 121 |
| | | M34 | 129 | <50 | 116 | 203 | 523 | 70 | 1653 | 1427 | 99 |
| | | M35 | 79 | 57 | 72 | 298 | 157 | <50 | 388 | 404 | 81 |
| | | M36 | 88 | 133 | 72 | 168 | 769 | 71 | 104 | 107 | 118 |

| Binding endpoint titers | |
|---|---|
| (white) | <50(LOD) |
| (grey) | 50(LOD)-500 |
| (light yellow) | 501-5000 |
| (orange) | 5001-50000 |
| (dark orange) | >50000 |

**Table S4 Endpoint antibody-binding titers of mice sera over time against SOSIP.664, SOSIP.v8 and gp120 proteins, as measured by D7324-capture ELISA.** Values are represented in **Fig. S8a.**

| | | | HIVconsvX | | | | | | | .664 | |
|---|---|---|---|---|---|---|---|---|---|---|---|
| Group | Regimen | Mouse | P1 | P2 | P3 | P4 | P5 | P6 | P7 | P14 | P15 |
| 1 | SOSIP.664 (CMP) | M1 | 50 | 0 | 0 | 0 | 0 | 0 | 0 | 120 | 95 |
| | | M2 | 0 | 0 | 0 | 0 | 0 | 0 | 0 | 90 | 165 |
| | | M3 | 0 | 0 | 0 | 0 | 0 | 0 | 0 | 175 | 205 |
| | | M4 | 0 | 0 | 0 | 0 | 0 | 150 | 0 | 480 | 180 |
| | | M5 | 0 | 0 | 0 | 0 | 0 | 0 | 0 | 885 | 70 |
| | | M6 | 200 | 0 | 0 | 0 | 0 | 0 | 0 | 580 | 170 |
| | | M7 | 0 | 0 | 200 | 0 | 350 | 800 | 0 | 280 | 135 |
| | | M8 | 0 | 0 | 0 | 0 | 0 | 0 | 0 | 340 | 445 |
| | | M9 | 0 | 0 | 0 | 0 | 0 | 0 | 0 | 375 | 330 |
| | | M10 | 0 | 0 | 0 | 150 | 500 | 1000 | 0 | 130 | 205 |
| | | M11 | 0 | 0 | 0 | 0 | 0 | 0 | 0 | 185 | 250 |
| | | M12 | 0 | 0 | 0 | 0 | 0 | 0 | 0 | 300 | 400 |
| 2 | TTT (CMP) | M13 | 0 | 0 | 0 | 0 | 0 | 0 | 0 | 5015 | 6075 |
| | | M14 | 0 | 0 | 0 | 0 | 0 | 0 | 0 | 6645 | 270 |
| | | M15 | 0 | 0 | 0 | 0 | 0 | 0 | 0 | 1745 | 150 |
| | | M16 | 0 | 5 | 0 | 0 | 0 | 0 | 0 | 8495 | 605 |
| | | M17 | 0 | 0 | 0 | 0 | 0 | 0 | 0 | 3605 | 100 |
| | | M18 | 0 | 0 | 0 | 0 | 0 | 0 | 0 | 50 | 115 |
| | | M19 | 0 | 0 | 0 | 5 | 0 | 0 | 0 | 550 | 95 |
| | | M20 | 0 | 0 | 0 | 0 | 0 | 0 | 0 | 30 | 80 |
| | | M21 | 0 | 0 | 0 | 0 | 0 | 0 | 0 | 35 | 45 |
| | | M22 | 0 | 0 | 10 | 0 | 0 | 0 | 0 | 60 | 120 |
| | | M23 | 0 | 0 | 0 | 0 | 0 | 0 | 0 | 8130 | 600 |
| | | M24 | 0 | 0 | 0 | 0 | 0 | 0 | 0 | 2200 | 145 |
| 3 | SOSIP.664 (PPP) C8M | M25 | 3230 | 5390 | 4685 | 4625 | 8200 | 9035 | 2135 | 100 | 80 |
| | | M26 | 1375 | 2375 | 3995 | 1975 | 8805 | 10195 | 1990 | 365 | 375 |
| | | M27 | 4580 | 7790 | 3145 | 7240 | 8275 | 8195 | 1720 | 395 | 430 |
| | | M28 | 3800 | 3055 | 1810 | 6650 | 8190 | 6930 | 1515 | 520 | 555 |
| | | M29 | 3620 | 9320 | 2860 | 7130 | 7890 | 7680 | 2520 | 395 | 405 |
| | | M30 | 4230 | 7455 | 655 | 8210 | 3705 | 5230 | 725 | 105 | 25 |
| 4 | TTT (CMP) C8M | M31 | 400 | 2085 | 0 | 2440 | 6005 | 6255 | 125 | 425 | 875 |
| | | M32 | 6615 | 8395 | 2535 | 5965 | 6645 | 6065 | 640 | 765 | 645 |
| | | M33 | 8570 | 7015 | 1600 | 6650 | 2380 | 3810 | 620 | 955 | 695 |
| | | M34 | 8365 | 9325 | 1055 | 7655 | 6335 | 5955 | 270 | 3275 | 655 |
| | | M35 | 8645 | 9700 | 405 | 7860 | 8110 | 3080 | 370 | 340 | 225 |
| | | M36 | 6225 | 7290 | 1115 | 6895 | 4485 | 6020 | 150 | 1045 | 270 |

| SFU/$10^6$ splenocytes | |
|---|---|
| (white) | 0-50 |
| (light grey) | 51-500 |
| (light yellow) | 501-1000 |
| (orange) | 1001-5000 |
| (dark orange) | >5000 |

**Table S5 Frequencies of mice-isolated splenocytes reactive to peptide pools covering HIVconsvX (P1-P7) and Env (P14-P15) epitopes.** Frequencies are expressed as spot-producing units (SFU) per $10^6$ splenocytes and represented in **Fig. S8c.**

| | | | CD4$^+$CD44$^+$IFN-γ$^+$ | | CD8$^+$CD44$^+$IFN-γ$^+$ | |
|---|---|---|---|---|---|---|
| **Group** | **Regimen** | **Mouse** | **.664 P14** | **.664 P15** | **.664 P14** | **.664 P15** |
| **1** | **SOSIP.664 (CMP)** | **M1** | 0.000 | 0.000 | 0.000 | 0.000 |
| | | **M2** | 0.000 | 0.000 | 0.000 | 0.000 |
| | | **M3** | 0.000 | 0.010 | 0.000 | 0.000 |
| | | **M4** | 0.045 | 0.000 | 0.020 | 0.000 |
| | | **M5** | 0.055 | 0.000 | 0.060 | 0.000 |
| | | **M6** | 0.045 | 0.000 | 0.040 | 0.000 |
| | | **M7** | 0.000 | 0.000 | 0.000 | 0.000 |
| | | **M8** | 0.015 | 0.130 | 0.000 | 0.170 |
| | | **M9** | 0.045 | 0.070 | 0.000 | 0.000 |
| | | **M10** | 0.000 | 0.050 | 0.000 | 0.000 |
| | | **M11** | 0.000 | 0.000 | 0.000 | 0.000 |
| | | **M12** | 0.025 | 0.110 | 0.000 | 0.010 |
| **2** | **TTT (CMP)** | **M13** | 0.144 | 0.120 | 0.120 | 0.310 |
| | | **M14** | 0.294 | 0.040 | 0.210 | 0.080 |
| | | **M15** | 0.114 | 0.060 | 0.050 | 0.000 |
| | | **M16** | 0.344 | 0.160 | 0.310 | 0.280 |
| | | **M17** | 0.214 | 0.120 | 0.110 | 0.000 |
| | | **M18** | 0.000 | 0.110 | 0.000 | 0.030 |
| | | **M19** | 0.160 | 0.070 | 0.030 | 0.000 |
| | | **M20** | 0.000 | 0.170 | 0.000 | 0.000 |
| | | **M21** | 0.050 | 0.040 | 0.000 | 0.000 |
| | | **M22** | 0.090 | 0.000 | 0.000 | 0.000 |
| | | **M23** | 0.200 | 0.100 | 0.270 | 0.050 |
| | | **M24** | 0.080 | 0.050 | 0.080 | 0.010 |
| **3** | **SOSIP.664 (PPP) C8M** | **M25** | 0.000 | 0.000 | 0.000 | 0.000 |
| | | **M26** | 0.020 | 0.000 | 0.000 | 0.000 |
| | | **M27** | 0.020 | 0.000 | 0.000 | 0.000 |
| | | **M28** | 0.060 | 0.050 | 0.000 | 0.050 |
| | | **M29** | 0.000 | 0.000 | 0.000 | 0.000 |
| | | **M30** | 0.000 | 0.000 | 0.000 | 0.000 |
| **4** | **TTT (CMP) C8M** | **M31** | 0.080 | 0.110 | 0.010 | 0.120 |
| | | **M32** | 0.100 | 0.070 | 0.020 | 0.000 |
| | | **M33** | 0.070 | 0.050 | 0.020 | 0.000 |
| | | **M34** | 0.120 | 0.150 | 0.150 | 0.000 |
| | | **M35** | 0.080 | 0.000 | 0.050 | 0.000 |
| | | **M36** | 0.160 | 0.000 | 0.090 | 0.000 |

| Tfh cells (%) | |
|---|---|
| (white) | 0-0.01 |
| (grey) | 0.011-0.05 |
| (light yellow) | 0.051-0.1 |
| (orange) | 0.11-0.2 |
| (dark orange) | >0.2 |

**Table S6 Frequencies of T follicular helper (Tfh) cells (PD-1$^+$CXCR5$^+$Bcl6$^+$ICOS$^+$CCR7$^-$) inside the effector-memory (CD44$^+$IFN-γ$^+$) CD4$^+$ and CD8$^+$ T cell compartments of mice-isolated splenocytes.** Frequencies are represented in **Fig. S8d.**

| Group | Regimen | Rabbit | Wk10 | | | Wk26 | | |
|---|---|---|---|---|---|---|---|---|
| | | | SOSIP.664 | SOSIP | gp120 | SOSIP.664 | SOSIP | gp120 |
| 1A | SOSIP.664 (CMP) | 37184 | 12191 | 116 | 51929 | 19542 | 595 | 120036 |
| | | 37185 | 3467 | 132 | 12416 | 8589 | 635 | 74675 |
| | | 37186 | 30726 | 182 | 95982 | 12565 | 197 | 71857 |
| | | 37187 | 7247 | 203 | 34014 | 8309 | 1661 | 69168 |
| | | 37188 | 9650 | 270 | 24601 | 22292 | 3613 | 94765 |
| | | 37189 | 3115 | 315 | 11244 | 2746 | 422 | 13058 |
| 1B | SOSIP.664 (CMP$^G$) | 37202 | 9769 | 197 | 48897 | 19829 | 1393 | 96015 |
| | | 37203 | 5150 | 327 | 25881 | 17523 | 3313 | 94316 |
| | | 37204 | 14279 | 310 | 54623 | 2555 | 265 | 8518 |
| | | 37205 | 5114 | 146 | 33514 | 5041 | 292 | 30397 |
| | | 37206 | 15088 | 374 | 44583 | 24623 | 1042 | 143830 |
| | | 37207 | 3922 | 157 | 11319 | 4505 | 300 | 18232 |
| 2A | TTT (CMP) | 37190 | 1716 | 1551 | 139 | 6382 | 1398 | 1563 |
| | | 37191 | 883 | 909 | 1589 | 12282 | 4186 | 18928 |
| | | 37192 | 2415 | 1764 | 390 | 8428 | 2555 | 20111 |
| | | 37193 | 1072 | 779 | 835 | 4342 | 1337 | 5338 |
| | | 37194 | 2071 | 1145 | 1050 | 11946 | 2714 | 23261 |
| | | 37195 | 3037 | 662 | 323 | 29300 | 5687 | 10871 |
| 2B | TTT (CMP$^G$) | 37208 | 868 | 679 | 411 | 12132 | 10492 | 9637 |
| | | 37209 | 2598 | 942 | 1994 | 19573 | 9008 | 30734 |
| | | 37210 | 1829 | 1320 | 1246 | 10707 | 10930 | 7696 |
| | | 37211 | 602 | 320 | 487 | 13199 | 11086 | 31968 |
| | | 37212 | 8207 | 2420 | 11940 | 30531 | 23966 | 118372 |
| | | 37213 | 1548 | 506 | 1236 | 23152 | 16771 | 49400 |
| 3 | SOSIP.664 (PPP) | 37178 | 6335 | 593 | 7576 | 14475 | 4279 | 28326 |
| | | 37179 | 29677 | 1167 | 36364 | 100974 | 18846 | 112702 |
| | | 37180 | 3512 | 99 | 1687 | 4148 | 1316 | 11224 |
| | | 37181 | 10317 | 149 | 12351 | 14733 | 2431 | 77063 |
| | | 37182 | 4059 | 123 | 12401 | ND | ND | ND |
| | | 37183 | 1331 | 181 | 925 | 2361 | 339 | 9322 |
| 4 | TTT (CP.MP.P) | 37196 | 17883 | 2419 | 18842 | 27284 | 8363 | 49040 |
| | | 37197 | 57186 | 6378 | 106234 | 37230 | 7382 | 59086 |
| | | 37198 | 15147 | 1520 | 20701 | 17897 | 4324 | 34182 |
| | | 37199 | 76859 | 3993 | 135124 | 15740 | 2866 | 36042 |
| | | 37200 | 77210 | 3332 | 62310 | 17786 | 3573 | 36404 |
| | | 37201 | 282744 | 5904 | 182871 | 15834 | 3097 | 38028 |
| 5 | TTT.GM (CMP$^G$) | 37214 | 1190 | 966 | 689 | 38943 | 13835 | 7779 |
| | | 37215 | 904 | 473 | 2902 | 9842 | 8234 | 13053 |
| | | 37216 | 64 | 90 | 122 | ND | ND | ND |
| | | 37217 | 1494 | 1478 | 2453 | 20808 | 12474 | 39662 |
| | | 37218 | 744 | 530 | 906 | 27926 | 24225 | 23464 |
| | | 37219 | 1139 | 532 | 2602 | 5825 | 4884 | 7664 |

| Binding endpoint titers | |
|---|---|
| (white) | 50(LOD)-500 |
| (light yellow) | 501-5000 |
| (orange) | 5001-50000 |
| (dark orange) | >50000 |

**Table S7 Endpoint antibody binding titers of rabbit at weeks 10 and 26 against SOSIP.664, SOSIP.v8 and gp120 proteins, as measured by D7324-capture ELISA.** Values are represented in **Fig. S9a.**

| Group | Regimen | Rabbit | Wk10 MLV Neg. Control | Wk10 BG505 Clade A Tier 2 | Wk26 MLV Neg. Control | Wk26 BG505 Clade A Tier 2 | Wk42 MLV Neg. Control | Wk42 BG505 Clade A Tier 2 |
|---|---|---|---|---|---|---|---|---|
| 1A | SOSIP.664 (CMP) | 37184 | <5 | <5 | <20 | <20 | <20 | <20 |
| | | 37185 | <5 | <5 | <20 | <20 | <20 | <20 |
| | | 37186 | <5 | <5 | <20 | <20 | <20 | <20 |
| | | 37187 | 5 | 216 | <20 | 2468 | <20 | 1154 |
| | | 37188 | <5 | <5 | <20 | 934 | <20 | 415 |
| | | 37189 | <5 | <5 | <20 | <20 | <20 | <20 |
| 1B | SOSIP.664 (CMP$^{G}$) | 37202 | 14 | 33 | <20 | 151 | <20 | <20 |
| | | 37203 | 6 | 14 | <20 | 43 | <20 | 855 |
| | | 37204 | <5 | 9 | <20 | 223 | <20 | 572 |
| | | 37205 | <5 | <5 | <20 | <20 | <20 | <20 |
| | | 37206 | <5 | 5 | <20 | 246 | <20 | 97 |
| | | 37207 | <5 | <5 | <20 | <20 | <20 | <20 |
| 2A | TTT (CMP) | 37190 | <5 | <5 | <20 | <20 | <20 | <20 |
| | | 37191 | <5 | <5 | <20 | 420 | <20 | 1442 |
| | | 37192 | <5 | <5 | <20 | <20 | <20 | 96 |
| | | 37193 | <5 | 53 | <20 | 257 | <20 | 3313 |
| | | 37194 | <5 | 15 | <20 | <20 | <20 | 9282 |
| | | 37195 | <5 | <5 | <20 | 210 | <20 | <20 |
| 2B | TTT (CMP$^{G}$) | 37208 | <5 | <5 | <20 | <20 | <20 | <20 |
| | | 37209 | <5 | <5 | 40 | 82 | <20 | <20 |
| | | 37210 | <5 | <5 | <20 | <20 | <20 | <20 |
| | | 37211 | <5 | <5 | <20 | <20 | <20 | <20 |
| | | 37212 | <5 | 2277 | <20 | 14611 | <20 | 24440 |
| | | 37213 | <5 | <5 | <20 | <20 | <20 | <20 |
| 3 | SOSIP.664 (PPP) | 37178 | <5 | <5 | <20 | 717 | <20 | 3097 |
| | | 37179 | <5 | 541 | <20 | 2398 | <20 | 1509 |
| | | 37180 | <5 | <5 | <20 | <20 | <20 | <20 |
| | | 37181 | <5 | 36 | <20 | 834 | <20 | 104 |
| | | 37182 | <5 | <5 | ND | ND | ND | ND |
| | | 37183 | <5 | <5 | <20 | <20 | <20 | 45 |
| 4 | TTT (CP.MP.P) | 37196 | <5 | <5 | <20 | 1598 | <20 | 376 |
| | | 37197 | <5 | 25 | <20 | 1223 | <20 | 326 |
| | | 37198 | 30 | 72 | <20 | <20 | <20 | <20 |
| | | 37199 | <5 | 81 | <20 | 172 | <20 | 671 |
| | | 37200 | <5 | <5 | <20 | 267 | <20 | 140 |
| | | 37201 | <5 | 9 | <20 | 72 | <20 | 264 |
| 5 | TTT.GM (CMP$^{G}$) | 37214 | <5 | <5 | <20 | <20 | <20 | <20 |
| | | 37215 | <5 | <5 | <20 | <20 | <20 | <20 |
| | | 37216 | <5 | <5 | ND | ND | ND | ND |
| | | 37217 | <5 | <5 | <20 | <20 | <20 | <20 |
| | | 37218 | <5 | <5 | <20 | <20 | <20 | <20 |
| | | 37219 | <5 | <5 | <20 | <20 | <20 | <20 |

| Neutralization |
|---|
| LOD-40 |
| 41-100 |
| 101-1000 |
| >1001 |

**Table S8 Midpoint neutralization titers ($ID_{50}$) over time for sera of the immunized rabbits against an autologous BG505/T332N pseudovirus.** $ID_{50}$ values were determined in a TZM-bl neutralization assay and are represented in **Fig. S9b.**

| BG505 TTT.GM | N88 | N133 | N137 | N156 | N160 | N190 | N190c | N197 | N234 | N241 | N262 | N276 | N289 | N295 | N301 | N332 | N339 | N355 | N363 | N386 | N392 | N398 | N406 | N411 | N448 | N462 | N611 | N618 | N625 | N637 | Linker |
|---|---|---|---|---|---|---|---|---|---|---|---|---|---|---|---|---|---|---|---|---|---|---|---|---|---|---|---|---|---|---|---|
| M9Glc | 0 | 0 | 0 | n.d. | 0 | 0 | n.d. | 0 | n.d. | 0 | 2 | 0 | 0 | 0 | 0 | 0 | 0 | 0 | 0 | 0 | 0 | 0 | n.d. | 0 | 0 | 0 | 0 | 0 | 0 | 0 | 0 |
| M9 | 0 | 0 | 0 | | 10 | 0 | | 1 | | 0 | 40 | 0 | 50 | 0 | 0 | 47 | 15 | 0 | 35 | 23 | 0 | 0 | | 0 | 22 | 0 | 1 | 0 | 0 | 0 | 0 |
| M8 | 1 | 0 | 0 | | 36 | 0 | | 12 | | 27 | 30 | 1 | 0 | 0 | 0 | 23 | 6 | 1 | 21 | 23 | 31 | 0 | | 12 | 17 | 0 | 1 | 0 | 1 | 3 | 0 |
| M7 | 4 | 0 | 0 | | 10 | 0 | | 4 | | 24 | 8 | 5 | 43 | 0 | 0 | 2 | 1 | 1 | 5 | 9 | 5 | 0 | | 13 | 8 | 0 | 7 | 1 | 0 | 5 | 0 |
| M6 | 13 | 0 | 0 | | 1 | 1 | | 3 | | 21 | 6 | 16 | 0 | 0 | 0 | 4 | 0 | 2 | 2 | 6 | 3 | 0 | | 0 | 7 | 0 | 2 | 0 | 3 | 7 | 0 |
| M5 | 15 | 55 | 0 | | 2 | 10 | | 11 | | 28 | 4 | 26 | 7 | 100 | 100 | 10 | 2 | 19 | 6 | 16 | 14 | 0 | | 37 | 17 | 15 | 12 | 1 | 1 | 25 | 0 |
| M4 | 0 | 7 | 0 | | 0 | 0 | | 1 | | 0 | 0 | 6 | 0 | 0 | 0 | 1 | 0 | 2 | 0 | 2 | 2 | 0 | | 0 | 3 | 0 | 3 | 0 | 0 | 1 | 0 |
| M3 | 0 | 0 | 0 | | 0 | 1 | | 1 | | 0 | 0 | 1 | 0 | 0 | 0 | 1 | 0 | 0 | 1 | 2 | 3 | 0 | | 1 | 1 | 0 | 0 | 0 | 0 | 0 | 0 |
| FM | 0 | 0 | 0 | | 0 | 1 | | 0 | | 0 | 0 | 0 | 0 | 0 | 0 | 0 | 0 | 0 | 0 | 0 | 0 | 0 | | 0 | 0 | 0 | 6 | 1 | 0 | 0 | 0 |
| Hybrid | 15 | 0 | 0 | | 1 | 1 | | 2 | | 0 | 0 | 1 | 0 | 0 | 0 | 1 | 0 | 3 | 1 | 3 | 0 | 0 | | 0 | 2 | 0 | 3 | 0 | 2 | 6 | 0 |
| Fhybrid | 0 | 0 | 0 | | 0 | 1 | | 2 | | 0 | 0 | 5 | 0 | 0 | 0 | 1 | 0 | 2 | 1 | 0 | 2 | 0 | | 0 | 1 | 1 | 2 | 0 | 0 | 1 | 0 |
| HexNAc(3)(x) | 5 | 0 | 0 | | 0 | 0 | | 1 | | 0 | 0 | 3 | 0 | 0 | 0 | 0 | 0 | 1 | 0 | 0 | 0 | 0 | | 0 | 0 | 0 | 5 | 2 | 5 | 0 | 0 |
| HexNAc(3)(F)(x) | 1 | 1 | 0 | | 0 | 2 | | 3 | | 0 | 0 | 4 | 0 | 0 | 0 | 1 | 0 | 8 | 1 | 0 | 3 | 43 | | 5 | 2 | 1 | 1 | 0 | 0 | 1 | 0 |
| HexNAc(4)(x) | 11 | 0 | 0 | | 0 | 0 | | 8 | | 0 | 0 | 0 | 0 | 0 | 0 | 0 | 0 | 0 | 0 | 0 | 0 | 0 | | 0 | 0 | 0 | 0 | 0 | 0 | 0 | 0 |
| HexNAc(4)(F)(x) | 10 | 34 | 0 | | 1 | 8 | | 3 | | 0 | 6 | 31 | 0 | 0 | 0 | 4 | 1 | 29 | 5 | 4 | 21 | 0 | | 30 | 11 | 42 | 13 | 7 | 0 | 5 | 0 |
| HexNAc(5)(x) | 6 | 0 | 0 | | 0 | 0 | | 23 | | 0 | 0 | 0 | 0 | 0 | 0 | 0 | 0 | 0 | 0 | 0 | 0 | 0 | | 0 | 0 | 0 | 0 | 0 | 0 | 0 | 0 |
| HexNAc(5)(F)(x) | 2 | 0 | 0 | | 0 | 6 | | 2 | | 0 | 3 | 0 | 0 | 0 | 0 | 4 | 0 | 8 | 2 | 1 | 6 | 57 | | 0 | 1 | 34 | 8 | 0 | 0 | 2 | 0 |
| HexNAc(6+)(x) | 1 | 0 | 0 | | 0 | 0 | | 10 | | 0 | 0 | 0 | 0 | 0 | 0 | 0 | 0 | 0 | 0 | 0 | 0 | 0 | | 0 | 0 | 0 | 0 | 0 | 0 | 0 | 0 |
| HexNAc(6+)(F)(x) | 1 | 2 | 0 | | 0 | 1 | | 0 | | 0 | 1 | 0 | 0 | 0 | 0 | 0 | 0 | 9 | 0 | 0 | 0 | 0 | | 0 | 0 | 3 | 3 | 0 | 0 | 0 | 0 |
| Unoccupied | 14 | 0 | 100 | | 38 | 64 | | 13 | | 0 | 0 | 0 | 0 | 0 | 0 | 0 | 73 | 14 | 19 | 7 | 7 | 0 | | 0 | 7 | 1 | 32 | 83 | 87 | 44 | 100 |
| Core | 0 | 0 | 0 | | 0 | 3 | | 1 | | 0 | 0 | 0 | 0 | 0 | 0 | 1 | 0 | 1 | 1 | 4 | 3 | 0 | | 0 | 1 | 2 | 1 | 1 | 0 | 0 | 0 |

**Table S9 Site-specific glycan analysis of the GNL-purified BG505 TTT.GM protein preparation.** Values indicate the percentage of each type of glycan (rows) at each glycosylation site (columns) and are represented in **Fig. S10g.**

| HA TTT | N22 | N38 | N45 | N63 | N122 | N126 | N133 | N144 | N165 | N246 | N285 | N483 |
|---|---|---|---|---|---|---|---|---|---|---|---|---|
| M9Glc | n.d. | 0 | 0 | 0 | 0 | 0 | 0 | 0 | 0 | 0 | 0 | 0 |
| M9 | | 0 | 0 | 0 | 0 | 0 | 0 | 0 | 8 | 20 | 2 | 0 |
| M8 | | 0 | 0 | 0 | 0 | 1 | 4 | 0 | 13 | 17 | 5 | 0 |
| M7 | | 0 | 0 | 0 | 0 | 1 | 0 | 0 | 13 | 2 | 8 | 0 |
| M6 | | 0 | 6 | 0 | 0 | 2 | 0 | 0 | 10 | 2 | 6 | 0 |
| M5 | | 2 | 27 | 7 | 0 | 30 | 96 | 0 | 38 | 24 | 44 | 0 |
| M4 | | 0 | 0 | 0 | 0 | 0 | 0 | 0 | 0 | 0 | 0 | 0 |
| M3 | | 0 | 0 | 0 | 0 | 0 | 0 | 0 | 0 | 0 | 0 | 0 |
| FM | | 0 | 0 | 0 | 0 | 0 | 0 | 0 | 0 | 0 | 0 | 1 |
| Hybrid | | 0 | 19 | 1 | 0 | 2 | 0 | 0 | 5 | 6 | 5 | 0 |
| Fhybrid | | 0 | 0 | 0 | 0 | 3 | 0 | 0 | 1 | 4 | 0 | 0 |
| HexNAc(3)(x) | | 0 | 29 | 1 | 0 | 1 | 0 | 0 | 1 | 2 | 5 | 0 |
| HexNAc(3)(F)(x) | | 3 | 1 | 2 | 0 | 10 | 0 | 0 | 3 | 3 | 8 | 0 |
| HexNAc(4)(x) | | 0 | 7 | 1 | 0 | 2 | 0 | 0 | 1 | 5 | 0 | 0 |
| HexNAc(4)(F)(x) | | 38 | 4 | 36 | 0 | 20 | 0 | 0 | 4 | 7 | 9 | 40 |
| HexNAc(5)(x) | | 0 | 7 | 3 | 0 | 0 | 0 | 0 | 0 | 2 | 0 | 0 |
| HexNAc(5)(F)(x) | | 29 | 0 | 45 | 0 | 19 | 0 | 0 | 3 | 5 | 0 | 36 |
| HexNAc(6+)(x) | | 0 | 0 | 0 | 0 | 0 | 0 | 0 | 0 | 0 | 0 | 0 |
| HexNAc(6+)(F)(x) | | 9 | 0 | 0 | 0 | 0 | 0 | 0 | 0 | 0 | 0 | 2 |
| Unoccupied | | 15 | 0 | 1 | 100 | 10 | 0 | 100 | 0 | 0 | 1 | 19 |
| Core | | 3 | 0 | 2 | 0 | 0 | 0 | 0 | 1 | 0 | 6 | 2 |

**Table S10 Site-specific glycan analysis of the StreptactinXT-purified H3 HA TTT protein preparation.** Values indicate the percentage of each type of glycan (rows) at each glycosylation site (columns) and are represented in **Fig. S13d.**

| BG505 protomer | N88 | N133 | N142 | N156 | N160 | N190 | N190c | N197 | N234 | N241 | N262 | N276 | N289 | N295 | N301 | N332 | N339 | N355 | N363 | N386 | N392 | N398 | N406 | N411 | N448 | N462 | N611 | N618 | N625 | N637 |
|---|---|---|---|---|---|---|---|---|---|---|---|---|---|---|---|---|---|---|---|---|---|---|---|---|---|---|---|---|---|---|
| M9Glc | 0 | 0 | n.d. | 0 | 0 | 0 | n.d. | 0 | n.d. | n.d. | 0 | 0 | 0 | n.d. | n.d. | 0 | 0 | 0 | 0 | 0 | 0 | 0 | n.d. | 0 | 0 | 0 | 0 | 0 | 0 | 0 |
| M9 | 0 | 0 | | 0 | 0 | 0 | | 10 | | | 67 | 1 | 22 | | | 8 | 14 | 1 | 8 | 11 | 16 | 0 | | 0 | 62 | 0 | 0 | 0 | 0 | 0 |
| M8 | 1 | 0 | | 15 | 3 | 0 | | 48 | | | 28 | 24 | 25 | | | 2 | 7 | 4 | 2 | 3 | 27 | 0 | | 3 | 18 | 0 | 0 | 0 | 0 | 4 |
| M7 | 3 | 0 | | 5 | 3 | 0 | | 16 | | | 4 | 19 | 15 | | | 0 | 1 | 6 | 0 | 1 | 4 | 0 | | 0 | 6 | 0 | 0 | 0 | 0 | 10 |
| M6 | 15 | 0 | | 0 | 3 | 0 | | 7 | | | 0 | 14 | 20 | | | 0 | 0 | 7 | 0 | 1 | 2 | 0 | | 4 | 11 | 0 | 0 | 0 | 0 | 8 |
| M5 | 16 | 11 | | 0 | 4 | 0 | | 0 | | | 0 | 31 | 7 | | | 0 | 0 | 42 | 0 | 2 | 3 | 3 | | 7 | 2 | 0 | 0 | 0 | 0 | 7 |
| M4 | 3 | 0 | | 0 | 1 | 0 | | 0 | | | 0 | 1 | 2 | | | 0 | 0 | 2 | 0 | 0 | 1 | 0 | | 0 | 0 | 0 | 0 | 0 | 0 | 1 |
| M3 | 0 | 0 | | 0 | 0 | 0 | | 0 | | | 0 | 0 | 1 | | | 0 | 0 | 0 | 0 | 0 | 0 | 0 | | 0 | 0 | 0 | 0 | 0 | 0 | 0 |
| FM | 0 | 0 | | 0 | 0 | 0 | | 0 | | | 0 | 0 | 0 | | | 0 | 0 | 0 | 0 | 0 | 0 | 1 | | 0 | 0 | 0 | 0 | 0 | 0 | 0 |
| Hybrid | 25 | 0 | | 0 | 0 | 0 | | 0 | | | 0 | 7 | 0 | | | 0 | 0 | 9 | 0 | 0 | 0 | 0 | | 0 | 0 | 0 | 0 | 0 | 0 | 5 |
| Fhybrid | 0 | 0 | | 0 | 0 | 0 | | 0 | | | 0 | 0 | 0 | | | 0 | 0 | 6 | 0 | 0 | 0 | 0 | | 0 | 0 | 0 | 0 | 0 | 0 | 4 |
| HexNAc(3)(x) | 5 | 0 | | 0 | 0 | 0 | | 0 | | | 0 | 1 | 0 | | | 0 | 0 | 1 | 0 | 0 | 0 | 0 | | 0 | 0 | 0 | 0 | 0 | 0 | 0 |
| HexNAc(3)(F)(x) | 0 | 0 | | 0 | 0 | 0 | | 0 | | | 0 | 0 | 0 | | | 0 | 0 | 7 | 0 | 0 | 0 | 24 | | 0 | 0 | 6 | 0 | 1 | 0 | 7 |
| HexNAc(4)(x) | 26 | 0 | | 0 | 0 | 0 | | 0 | | | 0 | 1 | 0 | | | 0 | 0 | 0 | 0 | 0 | 0 | 0 | | 0 | 0 | 0 | 0 | 0 | 0 | 0 |
| HexNAc(4)(F)(x) | 2 | 0 | | 0 | 0 | 0 | | 0 | | | 0 | 0 | 0 | | | 0 | 0 | 6 | 0 | 0 | 0 | 29 | | 0 | 0 | 33 | 0 | 4 | 0 | 19 |
| HexNAc(5)(x) | 3 | 0 | | 0 | 0 | 0 | | 0 | | | 0 | 0 | 0 | | | 0 | 0 | 0 | 0 | 0 | 0 | 0 | | 0 | 0 | 0 | 0 | 0 | 0 | 0 |
| HexNAc(5)(F)(x) | 1 | 0 | | 0 | 0 | 0 | | 0 | | | 0 | 0 | 0 | | | 0 | 0 | 5 | 0 | 0 | 0 | 44 | | 0 | 0 | 25 | 3 | 3 | 0 | 2 |
| HexNAc(6+)(x) | 0 | 0 | | 0 | 0 | 0 | | 0 | | | 0 | 0 | 0 | | | 0 | 0 | 0 | 0 | 0 | 0 | 0 | | 0 | 0 | 0 | 0 | 0 | 0 | 0 |
| HexNAc(6+)(F)(x) | 0 | 0 | | 0 | 0 | 0 | | 0 | | | 0 | 0 | 0 | | | 0 | 0 | 0 | 0 | 0 | 0 | 0 | | 0 | 0 | 0 | 0 | 0 | 0 | 0 |
| Unoccupied | 0 | 0 | | 41 | 31 | 100 | | 16 | | | 1 | 0 | 6 | | | 2 | 14 | 1 | 11 | 31 | 2 | 0 | | 29 | 0 | 0 | 92 | 63 | 97 | 8 |
| Core | 0 | 89 | | 39 | 55 | 0 | | 3 | | | 0 | 1 | 2 | | | 87 | 64 | 3 | 78 | 51 | 46 | 0 | | 58 | 1 | 35 | 5 | 28 | 1 | 26 |

| conM protomer | N88 | N130 | N136 | N141 | N156 | N160 | N188 | N197 | N234 | N241 | N262 | N276 | N289 | N295 | N301 | N332 | N339 | N356 | N386 | N392 | N403 | N406 | N448 | N461 | N611 | N616 | N625 | N637 |
|---|---|---|---|---|---|---|---|---|---|---|---|---|---|---|---|---|---|---|---|---|---|---|---|---|---|---|---|---|
| M9Glc | 0 | 0 | n.d. | n.d. | n.d. | n.d. | 0 | 0 | n.d. | n.d. | 0 | 0 | 0 | n.d. | 0 | n.d. | n.d. | 0 | 0 | 0 | n.d. | n.d. | 0 | n.d. | 0 | n.d. | 0 | 0 |
| M9 | 0 | 0 | | | | | 0 | 0 | | | 66 | 2 | 1 | | 0 | | | 0 | 23 | 4 | | | 80 | | 0 | | 0 | 11 |
| M8 | 0 | 0 | | | | | 0 | 0 | | | 27 | 54 | 5 | | 0 | | | 1 | 29 | 8 | | | 13 | | 0 | | 0 | 18 |
| M7 | 0 | 0 | | | | | 0 | 0 | | | 6 | 21 | 6 | | 4 | | | 0 | 13 | 0 | | | 3 | | 0 | | 0 | 12 |
| M6 | 5 | 0 | | | | | 0 | 0 | | | 0 | 9 | 3 | | 6 | | | 0 | 9 | 0 | | | 1 | | 0 | | 0 | 14 |
| M5 | 15 | 2 | | | | | 0 | 2 | | | 0 | 10 | 5 | | 6 | | | 2 | 5 | 1 | | | 1 | | 2 | | 0 | 10 |
| M4 | 0 | 0 | | | | | 0 | 0 | | | 0 | 1 | 0 | | 40 | | | 0 | 1 | 0 | | | 0 | | 0 | | 0 | 1 |
| M3 | 0 | 0 | | | | | 0 | 0 | | | 0 | 0 | 0 | | 5 | | | 0 | 0 | 0 | | | 0 | | 0 | | 0 | 0 |
| FM | 0 | 0 | | | | | 0 | 0 | | | 0 | 0 | 0 | | 0 | | | 0 | 0 | 0 | | | 0 | | 0 | | 0 | 0 |
| Hybrid | 0 | 0 | | | | | 0 | 1 | | | 0 | 2 | 0 | | 8 | | | 0 | 0 | 0 | | | 0 | | 0 | | 0 | 7 |
| Fhybrid | 0 | 0 | | | | | 0 | 1 | | | 0 | 0 | 0 | | 9 | | | 2 | 0 | 0 | | | 0 | | 0 | | 0 | 2 |
| HexNAc(3)(x) | 0 | 1 | | | | | 0 | 0 | | | 0 | 0 | 0 | | 0 | | | 0 | 0 | 0 | | | 0 | | 0 | | 0 | 0 |
| HexNAc(3)(F)(x) | 0 | 11 | | | | | 0 | 3 | | | 0 | 0 | 0 | | 11 | | | 6 | 0 | 0 | | | 0 | | 0 | | 0 | 0 |
| HexNAc(4)(x) | 14 | 0 | | | | | 0 | 0 | | | 0 | 0 | 0 | | 0 | | | 0 | 0 | 0 | | | 0 | | 0 | | 0 | 0 |
| HexNAc(4)(F)(x) | 53 | 50 | | | | | 0 | 7 | | | 0 | 0 | 0 | | 8 | | | 27 | 0 | 0 | | | 0 | | 0 | | 0 | 2 |
| HexNAc(5)(x) | 0 | 0 | | | | | 0 | 0 | | | 0 | 0 | 0 | | 0 | | | 0 | 0 | 0 | | | 0 | | 0 | | 0 | 0 |
| HexNAc(5)(F)(x) | 13 | 36 | | | | | 0 | 2 | | | 0 | 0 | 0 | | 0 | | | 61 | 0 | 0 | | | 0 | | 0 | | 0 | 0 |
| HexNAc(6+)(x) | 0 | 0 | | | | | 0 | 0 | | | 0 | 0 | 0 | | 0 | | | 0 | 0 | 0 | | | 0 | | 0 | | 0 | 0 |
| HexNAc(6+)(F)(x) | 0 | 0 | | | | | 0 | 0 | | | 0 | 0 | 0 | | 0 | | | 0 | 0 | 0 | | | 1 | | 0 | | 0 | 0 |
| Unoccupied | 0 | 0 | | | | | 99 | 6 | | | 1 | 0 | 80 | | 0 | | | 0 | 20 | 2 | | | 0 | | 93 | | 98 | 20 |
| Core | 0 | 0 | | | | | 0 | 77 | | | 0 | 0 | 0 | | 3 | | | 0 | 1 | 84 | | | 0 | | 5 | | 1 | 1 |

| AMC011 protomer | N88 | N130 | N136 | N141 | N142c | N156 | N160 | N188 | N197 | N234 | N241 | N262 | N276 | N289 | N295 | N301 | N332 | N339 | N355 | N362 | N386 | N392 | N396 | N406 | N448 | N460 | N611 | N616 | N625 | N637 |
|---|---|---|---|---|---|---|---|---|---|---|---|---|---|---|---|---|---|---|---|---|---|---|---|---|---|---|---|---|---|---|
| M9Glc | 0 | 0 | 0 | n.d. | n.d. | 0 | n.d. | 0 | 0 | n.d. | n.d. | 0 | 0 | 0 | n.d. | n.d. | 0 | 0 | n.d. | 0 | 0 | 0 | n.d. | 0 | 0 | n.d. | 0 | 0 | 0 | 0 |
| M9 | 0 | 0 | 0 | | | 0 | | 1 | 47 | | | 10 | 0 | 45 | | | 3 | 37 | | 46 | 29 | 4 | | 0 | 63 | | 0 | 0 | 0 | 0 |
| M8 | 0 | 0 | 0 | | | 0 | | 7 | 34 | | | 6 | 63 | 19 | | | 0 | 15 | | 8 | 8 | 8 | | 0 | 18 | | 0 | 0 | 0 | 1 |
| M7 | 0 | 0 | 0 | | | 0 | | 20 | 7 | | | 2 | 23 | 11 | | | 0 | 0 | | 2 | 3 | 0 | | 0 | 6 | | 0 | 0 | 0 | 1 |
| M6 | 0 | 0 | 0 | | | 0 | | 20 | 0 | | | 3 | 1 | 6 | | | 0 | 5 | | 1 | 1 | 0 | | 0 | 10 | | 0 | 0 | 0 | 4 |
| M5 | 21 | 2 | 0 | | | 0 | | 31 | 0 | | | 1 | 13 | 3 | | | 1 | 2 | | 1 | 2 | 1 | | 0 | 2 | | 0 | 0 | 6 | 3 |
| M4 | 0 | 0 | 0 | | | 0 | | 0 | 0 | | | 0 | 1 | 1 | | | 0 | 0 | | 0 | 0 | 0 | | 0 | 0 | | 0 | 0 | 0 | 1 |
| M3 | 0 | 0 | 0 | | | 0 | | 0 | 0 | | | 2 | 0 | 0 | | | 0 | 0 | | 0 | 0 | 0 | | 0 | 0 | | 0 | 0 | 0 | 0 |
| FM | 0 | 0 | 0 | | | 0 | | 0 | 0 | | | 0 | 0 | 0 | | | 0 | 0 | | 0 | 0 | 0 | | 35 | 0 | | 0 | 0 | 0 | 0 |
| Hybrid | 0 | 0 | 0 | | | 0 | | 12 | 0 | | | 0 | 0 | 0 | | | 0 | 0 | | 0 | 0 | 0 | | 0 | 0 | | 0 | 0 | 0 | 1 |
| Fhybrid | 0 | 0 | 0 | | | 0 | | 3 | 0 | | | 0 | 0 | 0 | | | 0 | 0 | | 0 | 0 | 0 | | 0 | 0 | | 0 | 0 | 0 | 0 |
| HexNAc(3)(x) | 0 | 1 | 90 | | | 0 | | 1 | 0 | | | 0 | 0 | 0 | | | 0 | 0 | | 0 | 0 | 0 | | 0 | 0 | | 0 | 0 | 0 | 0 |
| HexNAc(3)(F)(x) | 0 | 11 | 10 | | | 0 | | 1 | 0 | | | 0 | 0 | 0 | | | 0 | 0 | | 0 | 0 | 0 | | 45 | 0 | | 0 | 0 | 0 | 0 |
| HexNAc(4)(x) | 16 | 0 | 0 | | | 0 | | 0 | 0 | | | 0 | 0 | 0 | | | 0 | 0 | | 0 | 0 | 0 | | 0 | 0 | | 0 | 0 | 0 | 0 |
| HexNAc(4)(F)(x) | 41 | 50 | 0 | | | 0 | | 0 | 0 | | | 0 | 0 | 0 | | | 0 | 0 | | 0 | 0 | 0 | | 20 | 0 | | 0 | 0 | 0 | 1 |
| HexNAc(5)(x) | 0 | 0 | 0 | | | 0 | | 0 | 0 | | | 0 | 0 | 0 | | | 0 | 0 | | 0 | 0 | 0 | | 0 | 0 | | 0 | 0 | 0 | 0 |
| HexNAc(5)(F)(x) | 22 | 36 | 0 | | | 0 | | 0 | 0 | | | 0 | 0 | 0 | | | 0 | 0 | | 0 | 0 | 0 | | 0 | 0 | | 0 | 0 | 0 | 0 |
| HexNAc(6+)(x) | 0 | 0 | 0 | | | 0 | | 0 | 0 | | | 0 | 0 | 0 | | | 0 | 0 | | 0 | 0 | 0 | | 0 | 0 | | 0 | 0 | 0 | 0 |
| HexNAc(6+)(F)(x) | 0 | 0 | 0 | | | 0 | | 0 | 0 | | | 0 | 0 | 0 | | | 0 | 0 | | 0 | 0 | 0 | | 0 | 0 | | 0 | 0 | 0 | 0 |
| Unoccupied | 0 | 0 | 0 | | | 0 | | 1 | 11 | | | 0 | 0 | 13 | | | 0 | 41 | | 40 | 0 | 2 | | 0 | 0 | | 100 | 100 | 94 | 87 |
| Core | 0 | 0 | 0 | | | 100 | | 2 | 0 | | | 75 | 0 | 1 | | | 96 | 0 | | 1 | 57 | 84 | | 0 | 1 | | 0 | 0 | 0 | 0 |

**Table S11 Site-specific glycan analysis of the PGT145-purified chEnv TTT protein preparation.** Values indicate the percentage of each type of glycan (rows) at each glycosylation site (columns) and are represented in **Fig. S15e.**

| cH125 TTT | N20/21[1] | N20/21[2,3] | N33[1,2,3] | N94[1] | N158[3] | N169[3] | N169/170[2] | N278[1] | N289[1] | N289[2] | N289[3] | N483[1,2,3] |
|---|---|---|---|---|---|---|---|---|---|---|---|---|
| M9Glc | 0 | 0 | 0 | 0 | 0 | 0 | 0 | 0 | 0 | 0 | 0 | 0 |
| M9 | 0 | 0 | 0 | 0 | 0 | 0 | 0 | 0 | 2 | 0 | 0 | 0 |
| M8 | 0 | 1 | 0 | 5 | 0 | 0 | 0 | 6 | 12 | 1 | | 1 |
| M7 | 0 | 3 | 2 | 15 | 1 | 0 | 0 | 5 | 6 | 1 | 0 | 1 |
| M6 | 0 | 1 | 1 | 5 | 1 | 0 | 0 | 1 | 6 | 1 | 1 | 1 |
| M5 | 7 | 10 | 22 | 23 | 8 | 7 | 6 | 13 | 51 | 9 | 10 | 19 |
| M4 | 7 | 0 | 0 | 0 | 1 | 0 | 1 | 0 | 3 | 1 | 1 | 1 |
| M3 | 2 | 0 | 1 | 0 | 1 | 0 | 0 | 0 | 0 | 0 | 0 | 1 |
| FM | 0 | 0 | 2 | 0 | 1 | 1 | 1 | 0 | 0 | 0 | 0 | 1 |
| Hybrid | 0 | 0 | 0 | 2 | 0 | 0 | 0 | 1 | 6 | 1 | 0 | 1 |
| Fhybrid | 0 | 0 | 0 | 1 | 0 | 0 | 0 | 0 | 0 | 0 | 0 | 0 |
| HexNAc(3)(x) | 0 | 1 | 1 | 1 | 0 | 0 | 0 | 2 | 2 | 1 | 0 | 1 |
| HexNAc(3)(F)(x) | 2 | 0 | 4 | 2 | 4 | 3 | 3 | 2 | 0 | 4 | 2 | 4 |
| HexNAc(4)(x) | 0 | 1 | 1 | 2 | 0 | 0 | 1 | 2 | 2 | 2 | 0 | 3 |
| HexNAc(4)(F)(x) | 2 | 20 | 32 | 23 | 25 | 40 | 40 | 40 | 2 | 38 | 29 | 28 |
| HexNAc(5)(x) | 0 | 1 | 0 | 1 | 0 | 0 | 0 | 0 | 0 | 1 | 1 | 1 |
| HexNAc(5)(F)(x) | 3 | 15 | 16 | 10 | 29 | 46 | 33 | 19 | 0 | 23 | 29 | 13 |
| HexNAc(6+)(x) | 0 | 0 | 0 | 0 | 0 | 0 | 0 | 0 | 0 | 0 | 0 | 1 |
| HexNAc(6+)(F)(x) | 0 | 33 | 12 | 0 | 5 | 0 | 4 | 0 | 0 | 6 | 12 | 11 |
| Unoccupied | 38 | 0 | 4 | 9 | 2 | 1 | 0 | 2 | 1 | 0 | 1 | 3 |
| Core | 39 | 14 | 1 | 0 | 22 | 2 | 10 | 5 | 4 | 12 | 13 | 9 |

[1]Present in the H1 protomer [2]Present in the H2 protomer [3]Present in the H5 protomer

| cH125/111 TTT | N20/21[1,2,3] | N33[1,2,3] | N94[1] | N158[3] | N169[3] | N169/170[2] | N289[1,2,3] | N483[1,2,3] |
|---|---|---|---|---|---|---|---|---|
| M9Glc | 0 | 0 | 0 | 0 | 0 | 0 | 0 | 0 |
| M9 | 0 | 2 | 0 | 0 | 0 | 0 | 0 | 1 |
| M8 | 0 | 3 | 0 | 0 | 0 | 2 | 1 | 3 |
| M7 | 2 | 2 | 4 | 0 | 1 | 2 | 0 | 4 |
| M6 | 1 | 1 | 5 | 0 | 1 | 2 | 1 | 4 |
| M5 | 14 | 5 | 27 | 2 | 3 | 8 | 6 | 13 |
| M4 | 7 | 2 | 0 | 1 | 1 | 1 | 0 | 0 |
| M3 | 0 | 1 | 0 | 0 | 0 | 0 | 0 | 0 |
| FM | 0 | 0 | 0 | 0 | 0 | 0 | 0 | 0 |
| Hybrid | 0 | 2 | 17 | 0 | 0 | 1 | 3 | 4 |
| Fhybrid | 3 | 2 | 0 | 0 | 0 | 0 | 1 | 2 |
| HexNAc(3)(x) | 0 | 1 | 0 | 0 | 0 | 0 | 1 | 2 |
| HexNAc(3)(F)(x) | 1 | 4 | 3 | 1 | 2 | 0 | 4 | 3 |
| HexNAc(4)(x) | 3 | 0 | 5 | 0 | 0 | 0 | 1 | 1 |
| HexNAc(4)(F)(x) | 36 | 34 | 25 | 40 | 44 | 51 | 47 | 32 |
| HexNAc(5)(x) | 0 | 0 | 1 | 0 | 0 | 1 | 0 | 0 |
| HexNAc(5)(F)(x) | 18 | 20 | 10 | 50 | 39 | 28 | 22 | 13 |
| HexNAc(6+)(x) | 0 | 0 | 0 | 0 | 0 | 0 | 0 | 0 |
| HexNAc(6+)(F)(x) | 15 | 16 | 0 | 5 | 5 | 3 | 10 | 4 |
| Unoccupied | 0 | 0 | 2 | 0 | 1 | 0 | 0 | 14 |
| Core | 0 | 3 | 0 | 0 | 2 | 1 | 3 | 0 |

[1]Present in the H1/1 protomer [2]Present in the H2/1 protomer [3]Present in the H5/1 protomer.

**Table S12 Site-specific glycan analysis of the StrepTactinXT-purified cH125 and cH125/111 TTT protein preparations.** Values indicate the percentage of each type of glycan (rows) at each glycosylation site (columns) and are represented in **Fig. S19d.**

| Protein | +CR9114 | +5J8 | +2B05 | 2D class averages | 3D model | Comments |
|---|---|---|---|---|---|---|
| H1 | Full | Partial | - | 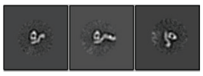 | - | Sample contains mostly monomers. All particles bind CR9114, but only a portion bind 5J8. |
| H1 GCN4 | 3/3 | 1/3 | - | 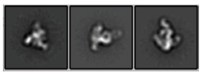 | - | - |
| | - | - | 3/3 | 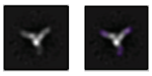 | - | **Evidence of cH125/TTT chimerism:** particles bind 3x2B05 Fabs (H1 head-specific) |
| H2 GCN4 | - | - | - | 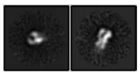 | - | - |
| | 0/3 | - | - | 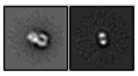 | - | No CR9114 binding, consistent with ELISA data. |
| | - | 0/3 | - | 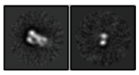 | - | No 5J8 binding, consistent with ELISA data. |
| | - | - | 0/3 | 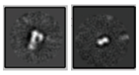 | - | **Evidence of cH125/TTT chimerism:** no 2B05 binding (H1 head-specific). |
| H5 GCN4 | 3/3 | - | - | - | 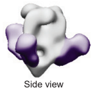  | - |
| | - | 0/3 | - | 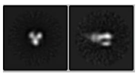 | - | No 5J8 binding, consistent with ELISA data. |
| | - | - | 0/3 | 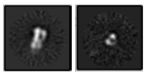 | - | **Evidence of cH125/TTT chimerism:** no 2B05 binding (H1 head-specific). |
| cH125 TTT | 3/3 | 0/3 | - | 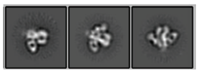 | 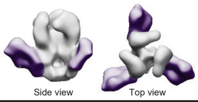  | No 5J8 binding observed, in contrast with ELISA data. |
| H1/1 | Full | Partial | - | 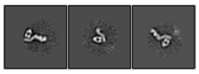 | - | Sample contains mostly monomers. All particles bind CR9114, but only a portion bind 5J8. |
| H1/1 GCN4 | 3/3 | Partial | - | 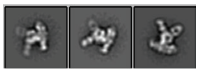 | - | Sample contains only trimers, with full occupancy of CR9114 epitopes, but partical occupancy of 5J8 epitopes (1-2x5J8 per particle). |
| H2/1 GCN4 | 3/3 | 1/3 | - | 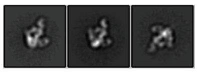 | 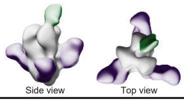  | Most particles bound by 1x5J8 Fab, in contrast with ELISA data. |
| H5/1 GCN4 | 3/3 | 0/3 | - | 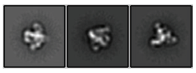 | 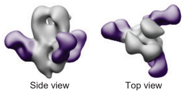  | - |
| | - | 0/3 | - | 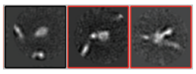 | - | 5J8 induced trimer degradation. |
| cH125/111 TTT | 3/3 | 0/3 | - | 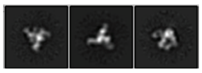 | 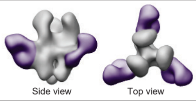  | No 5J8 binding observed, in contrast with ELISA data. |
| | - | - | 1/3 | 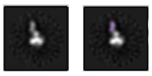 | - | **Evidence of cH125/TTT chimerism:** particles bind 1x2B05 Fab (H1 head-specific). |

**Table S13 nsEM data of cH125 TTT, cH125/111 TTT and the monomeric and GCN4-trimerized controls.** Proteins were imaged independently or complexed with CR9114, 5J8 or 2B05 fabs. Columns 2-4 indicate the occupancy of epitopes for the corresponding complexed antibody.
